# Piezo inhibition prevents *and* rescues scarring by targeting the adipocyte to fibroblast transition

**DOI:** 10.1101/2023.04.03.535302

**Authors:** Michelle F. Griffin, Heather E. Talbott, Nicholas J. Guardino, Jason L. Guo, Amanda F. Spielman, Kellen Chen, Jennifer B.L. Parker, Shamik Mascharak, Dominic Henn, Norah Liang, Megan King, Asha C. Cotterell, Khristian E. Bauer-Rowe, Darren B. Abbas, Nestor M. Diaz Deleon, Dharshan Sivaraj, Evan J. Fahy, Mauricio Downer, Deena Akras, Charlotte Berry, Jessica Cook, Natalina Quarto, Ophir D. Klein, H. Peter Lorenz, Geoffrey C. Gurtner, Michael Januszyk, Derrick C. Wan, Michael T. Longaker

**Affiliations:** Department of Surgery, Division of Plastic and Reconstructive Surgery, Stanford University School of Medicine; Stanford, CA 94305, USA; Institute for Stem Cell Biology and Regenerative Medicine, Stanford University School of Medicine; Stanford, CA 94305, USA; Department of Orofacial Sciences and Program in Craniofacial Biology, University of California San Francisco, San Francisco, CA 94143, USA; Department of Pediatrics, Cedars-Sinai Medical Center, Los Angeles, CA 90048, USA

**Keywords:** adipocyte, adipose tissue, adipocyte plasticity, fibroblast, fibrosis, scarring, mechanotransduction

## Abstract

While past studies have suggested that plasticity exists between dermal fibroblasts and adipocytes, it remains unknown whether fat actively contributes to fibrosis in scarring. We show that adipocytes convert to scar-forming fibroblasts in response to *Piezo*-mediated mechanosensing to drive wound fibrosis. We establish that mechanics alone are sufficient to drive adipocyte-to-fibroblast conversion. By leveraging clonal-lineage-tracing in combination with scRNA-seq, Visium, and CODEX, we define a “mechanically naïve” fibroblast-subpopulation that represents a transcriptionally intermediate state between adipocytes and scar-fibroblasts. Finally, we show that *Piezo1* or *Piezo2*-inhibition yields regenerative healing by preventing adipocytes’ activation to fibroblasts, in both mouse-wounds and a novel human-xenograft-wound model. Importantly, *Piezo1*-inhibition induced wound regeneration even in *pre-existing* established scars, a finding that suggests a role for adipocyte-to-fibroblast transition in wound remodeling, the least-understood phase of wound healing. Adipocyte-to-fibroblast transition may thus represent a therapeutic target for minimizing fibrosis via *Piezo*-inhibition in organs where fat contributes to fibrosis.

## Introduction

Skin scarring is an acute fibrotic process that occurs following any injury to the adult dermis. Scarring affects over 100 million patients every year in the U.S. and can cause both visual disfigurement and functional impairment (e.g., growth restriction, joint contraction).(desJardins-Park et al., 2021; Griffin et al., 2020; Gurtner et al., 2008) The most downstream mediators of scarring are dermal fibroblasts, which deposit the excess, fibrotic extracellular matrix (ECM) comprising a scar.(Griffin et al., 2021; Gurtner et al., 2008) The potential contributions of subcutaneous adipose tissue to skin fibrosis are relatively undefined but of growing interest.(Sakers et al., 2022) Recent studies have revealed lineage plasticity between adipocytes and fibroblasts, suggesting that fibroblasts may be able to differentiate into adipocytes(Plikus et al., 2017) and vice versa.(Shook et al., 2020) Clinical and experimental correlates also implicate adipose tissue in the balance between fibrosis and regeneration: adipose tissue is lost in multiple fibrotic skin conditions (e.g., systemic sclerosis,(Marangoni and Lu, 2017) radiation fibrosis(Poglio et al., 2009)), whereas adipogenic differentiation has been associated with presence of neogenic hair follicles in large wounds.(Plikus et al., 2017) However, much remains unknown about adipocyte dynamics, including the function, molecular drivers, and effects of modulating adipocyte-to-fibroblast conversion in scarring.

Tissue mechanical forces are a critical mediator of scarring. This has long been recognized by surgeons, who incise along lines of relaxed skin tension (“Langer’s lines”) to minimize postoperative scarring.(Wong et al., 2010) Experimentally, increasing tension across wounds increases scarring in mice;(Aarabi et al., 2007) conversely, inhibiting wound mechanics - using either a tension-offloading elastomeric dressing or small molecule inhibitors of mechanotransduction – has been shown to reduce scarring in both animal models and human clinical trials.(Longaker et al., 2014; Talbott et al., 2022) While it is well established that fibroblasts are highly mechanosensitive – for instance, we recently showed that mechanical forces activate a subset of dermal fibroblasts to adopt a pro-fibrotic phenotype(Mascharak et al., 2021a) – relatively little is known about mechanoresponsiveness of dermal adipocytes. Rare *in vitro* studies suggest that adipocytes modify gene expression in response to substrate mechanics,(Hossain et al., 2010; Yuan et al., 2015) but understanding of adipocyte mechanical signaling remains extremely limited, and the effects of wound mechanical cues on adipocytes are entirely unknown.

Here, we apply multiple *in vitro* and *in vivo* models to accomplish lineage tracing of adipocytes and study their phenotypic response to substrate/wound mechanics. We show for the first time that mechanics can dramatically alter wound adipocyte fate: mechanical cues promote conversion of adipocytes into pro-fibrotic fibroblasts within wounds, via an intermediate, “mechanically naïve” fibroblast state. Further, blocking adipocyte mechanosensing, via blockade of *Piezo1* or *Piezo2* mechanosensitive ion channel components, prevents adipocyte-to-fibroblast transition and significantly decreases fibrosis, promoting wound regeneration, even in existing scars. We utilize a multi-omic approach including scRNA-seq and spatial genomic and proteomic analyses to define the mechanosensitive and mechanically naïve fibroblast populations during wound repair and existing scars. Collectively, this study elucidates the role and molecular drivers of adipocyte-to-fibroblast conversion in wound repair and scarring, which may represent a novel therapeutic target for preventing scarring and other fibroses.

## Results

### Adipocytes transition to ECM-producing fibroblasts during wound repair and contribute to scarring

To trace adipocyte fate in wounds, we first developed an adipocyte transplantation and wounding model (**Fig. S1A**). We transplanted Tomato^+^ adipocytes from *R26^mTmG^* mice into the dorsal dermis of non-fluorescent wildtype recipient mice, then wounded within the cell-engrafted region. Wounds were stented using silicone rings to prevent rapid contraction,(Mascharak et al., 2021a; Mascharak et al., 2022) and excised skin was saved as an unwounded cell-engrafted control. In unwounded skin, few Tomato^+^ cells were observed (**Fig. S1B**, dotted orange arrows). In contrast, in wounds harvested at postoperative day (POD) 14 (the time of complete wound re-epithelialization), a greater number of Tomato^+^ cells were observed; interestingly, these cells had phenotypic properties of both adipocyte and fibroblast identity, expressing adiponectin (Adipoq; mature adipocyte marker) and type 1 collagen (Col1; not substantially expressed by adipocytes at homeostasis) (**Fig. S1C**, solid white arrows), suggesting that these cells may be in the process of transitioning from adipocytes to fibroblasts. We also observed Tomato^+^ cells that were Adipoq^-^/Col1^+^ (**Fig. S1C**, dotted white arrows) and a significant increase in the number of transplanted (Tomato^+^) cells expressing Col1 by POD 14 (**Fig. S1D**), consistent with adipocytes having differentiated into fibroblasts in healing wounds.

While these results strongly suggested that adipocytes in wounds transitioned to fibroblasts, it was possible that a small number of contaminating fibroblasts could have been inadvertently engrafted and then proliferated within wounds to produce the observed Tomato^+^ fibroblasts. To address this potential concern, we generated, tamoxifen-induced, and wounded *Adipoq^Cre-ERT^;R26^mTmG^* mice (**Fig. 1A and S1E**). In this model, mature adipocytes (*Adipoq^+^*) and their progeny express green fluorescent protein (GFP), allowing robust identification of wound fibroblasts that arose from adipocytes (GFP^+^); all other cells express Tomato. On histology, GFP expression in uninjured skin was confined to large, round subcutaneous cells consistent with mature adipocytes; over the course of wound healing, GFP^+^ cells increased in prevalence throughout the dermis (**Fig. 1B-C**). Fluorescence-activated cell sorting (FACS; based on lineage depletion as previously published(Leavitt et al., 2017)) showed that GFP^+^ (adipocyte-derived) fibroblasts were rare in unwounded skin but increased in wounds over time, with adipocyte-derived fibroblasts (ADFs) comprising ∼10% of wound fibroblasts by POD 14 (**Fig. 1D and S1F**). Further supporting fibroblast identity, GFP^+^ cells in POD 14 wounds did not express endothelial or immune markers (**Fig. S2A-B**), largely failed to express adipocyte markers (e.g., perilipin), and instead expressed typical scar fibroblast markers (e.g., collagens, alpha-smooth muscle actin [*α*SMA]; **Fig. S2C**, **top and middle rows**). These ADFs also colocalized with known fibroblast signaling factors such as transforming growth factor beta (TGF*β*) (**Fig. S2C**, **bottom row**). Collectively, these results were consistent with ADFs losing their adipocyte identity over the course of wound healing and transitioning to activated, ECM-producing fibroblasts.

**Figure 1:**
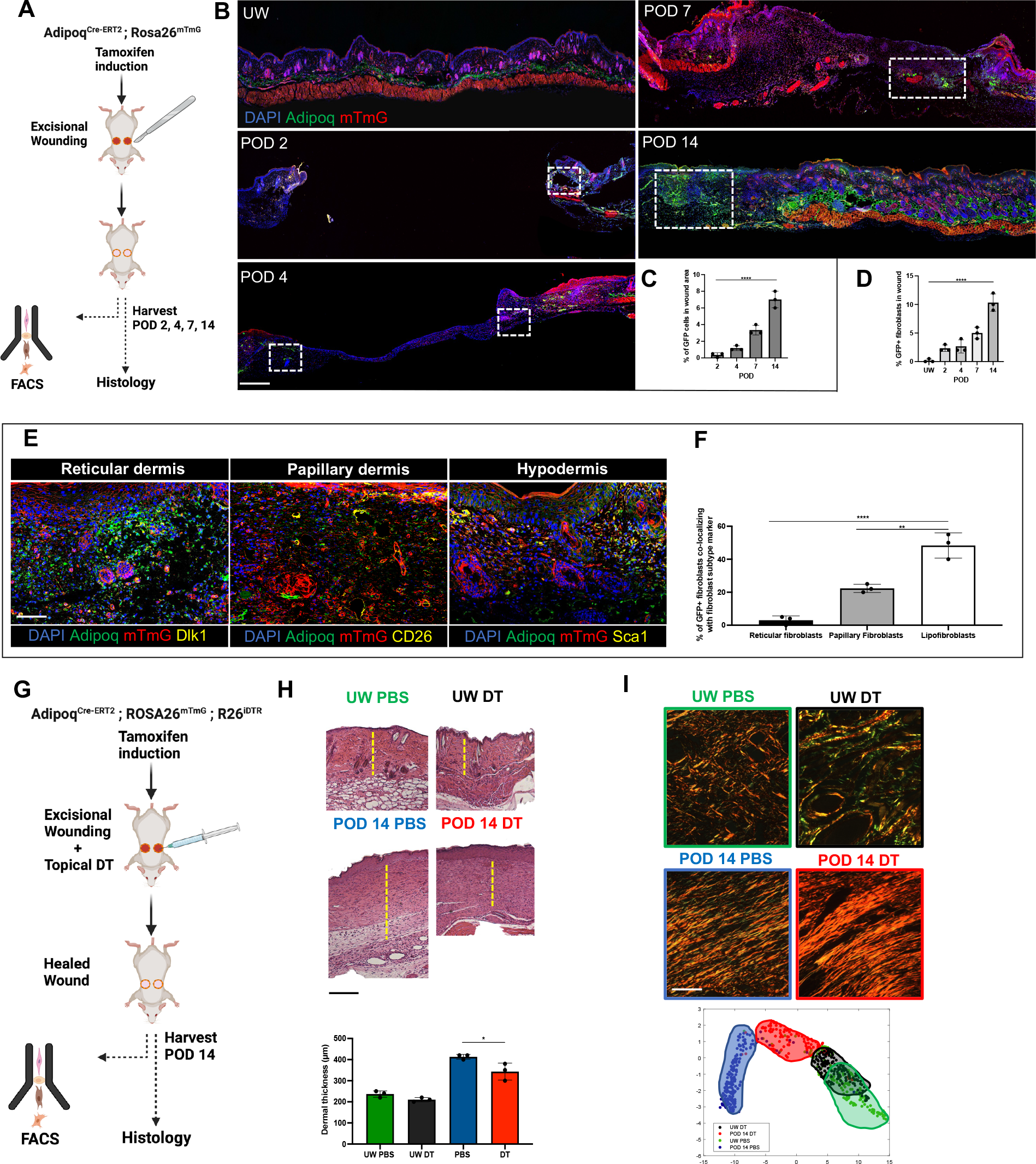
Adipocytes transition to fibroblasts in wounds and contribute to scarring. (**A**) Schematic of *Adipoq^Cre-ERT^;R26^mTmG^* wounding experiments for adipocyte lineage tracing. (**B**) Fluorescent histology of unwounded skin (UW) or wounds at indicated postoperative day (POD). White dotted regions highlight GFP^+^ cells within wounds. (**C**) Quantification of GFP^+^ wound cells as percentage of wound area. (**D**) Quantification of GFP^+^ fibroblasts as percentage of all fibroblasts on fluorescence-activated cell sorting (FACS; see *Fig. S1F* for FACS strategy). (**E**) Fluorescent histology of POD 14 wounds with immunofluorescent (IF) staining for indicated fibroblast subtype markers (yellow fluorescent signal). (**F**) Quantification of percentage of GFP^+^ wound cells colocalizing with given fibroblast subtype marker by area. (**G**) Schematic of *Adipoq^Cre- ERT^;R26^mTmG^;R26^iDTR^* wounding experiments for adipocyte ablation. (**H**) Hematoxylin and eosin (H&E) staining (top) and dermal thickness measurements (bottom) of skin (UW) and POD 14 wounds treated with PBS (control) or Dipheria Toxin (DT) (to ablate adipocytes). Yellow dotted lines show dermal thickness in representative sections. (**I**) Picrosirius red histology (top) and uniform manifold approximation and projection (UMAP) mapping of quantified ECM ultrastructure parameters (bottom; each dot represents one histology image) for PBS- and DT-treated skin and wounds. (B, E) DAPI (4′,6-diamidino-2-phenylindole), nuclear counterstain (blue fluorescent signal); mTmG, Tomato^+^ cells (red fluorescent signal; background color of non-Cre- expressing cells in *mTmG* construct); Adipoq, GFP^+^ cells (green fluorescent signal; *Adipoq^Cre^* lineage-positive cells). (**C, D, F, H**) Data shown as mean ± standard deviation (S.D.). *\*P* ≤ 0.05, \*\**P* ≤ 0.01, \*\*\*\**P* ≤ 0.0001. Scale bars, 500 µm (**B**), 25 µm (**E**), 150 µm (**H**), 20 µm (**I**).

Unwounded skin contains multiple anatomically distinct fibroblast populations with unique surface markers, which we and others have implicated in differing wound phenotypes and fibrogenic properties.(Driskell et al., 2013; Mascharak et al., 2021a) We used histology and multi- color FACS (**Fig. S1G**) profiling of unwounded skin and wounds in *Adipoq^Cre-ERT^;R26^mTmG^* mice to separately identify fibroblasts of the papillary dermis (CD26^+^Sca1^-^), reticular dermis (Dlk1^+^Sca1^-^), and hypodermis (lipofibroblasts; Dlk1^+/-^Sca1^+^). The majority of ADFs were Sca1^+^ fibroblasts (lipofibroblasts); a smaller number expressed papillary and reticular dermal fibroblast markers (**Fig. 1E-F and S1G-H**).

We next developed *Adipoq^Cre-ERT^;R26^mTmG^;R26^iDTR^* mice to enable selective ablation of mature adipocytes following tamoxifen induction (**Fig. 1G**). Confirming efficacy of ablation, unwounded skin and POD 14 wounds treated with phosphate-buffered saline (PBS) vehicle control contained abundant subdermal adipocytes and GFP^+^ ADFs, while those treated with diphtheria toxin (DT) had minimal to no subdermal fat and no GFP^+^ cells in wounds (**Fig. 1H, top panels, and Fig. S3A**). Further, DT-treated scars had significantly reduced dermal thickness (**Fig. 1H**, **bottom panel**), less dense extracellular matrix (ECM), and decreased Col1 and *α*-SMA content compared to PBS control scars (**Fig. S3B-C**). Quantitative analysis of ECM ultrastructure(Mascharak et al., 2021a; Mascharak et al., 2022) revealed that ECM of DT-treated wounds was less scar-like and more closely resembled unwounded skin, compared to the distinct scar ECM structure of control (PBS) wounds (**Fig. 1I**). Scars, despite being more densely collagenous than skin, are mechanically inferior, only ever regaining up to 80% of the strength of unwounded skin;(Mascharak et al., 2021b; Mascharak et al., 2022) adipocyte-ablated wounds had mechanical properties intermediate between those of unwounded skin and control scars (**Fig. S3D**).

### Adipocyte-derived fibroblasts comprise distinct “mechanically naïve” and “mechanically activated” subpopulations

While complete adipocyte ablation did reduce scarring, this effect fell short of complete regeneration. Although our results supported a role for adipocytes in *driving* fibrosis, prior studies and clinical correlates suggest that adipocytes may *mitigate* fibrosis in certain settings;(Almadori et al., 2019) we thus wondered whether there may exist multiple dermal adipocyte subtypes with distinct functional roles (i.e., some pro-scarring, some pro-regenerative). If this were the case, a more optimal strategy may be to specifically target only those pro-scarring adipocytes, rather than blanket ablation of all adipocytes. We hypothesized that a subset of adipocytes may respond to wound-specific cues to adopt a pro-fibrotic fibroblast phenotype, and sought to robustly characterize the dynamics and drivers of adipocyte-to-fibroblast transition.

We first examined the cellular proliferation dynamics of adipocytes’ contributions to scarring. EdU analysis revealed that adipocytes were minimally proliferative in both wounds and unwounded skin, whereas EdU incorporation appeared in ADFs by POD 4 and increased by POD 7 and 14 (**Fig. S4A-C**), suggesting an initial adipocyte-to-fibroblast transition followed by proliferation of ADFs post-transition. We next used *R26^VT2/GK3^* (“Rainbow”) reporter mice to study adipocyte clonal dynamics in wounds. In these mice, all cells initially express GFP; Cre induces stochastic *R26^VT2/GK3^* cassette recombination, resulting in random expression of one of three distinct fluorophores (mCerulean [blue], mOrange [orange], or mCherry [red]). Following recombination, that cell and its progeny will express the same color, allowing histologic identification of clonal expansion (clusters of multiple, identically colored cells). We performed tamoxifen induction and splinted excisional wounding of *Adipoq^Cre-ERT^;R26^VT2/GK3^* mice (**Fig. 2A**). POD 14 *Adipoq^Cre-ERT^;R26^VT2/GK3^* wounds contained numerous ADFs (mOrange^+^, mCerulean^+^, or mCherry^+^ cells that colocalized with Col1) and clusters of same-colored ADFs (**Fig. 2B-C**). Collectively, these results supported that individual adipocytes give rise to a larger number of fibroblasts via polyclonal expansion, suggesting that a subset of adipocytes may possibly serve as pro-fibrotic fibroblast progenitors in the injury setting.

**Figure 2:**
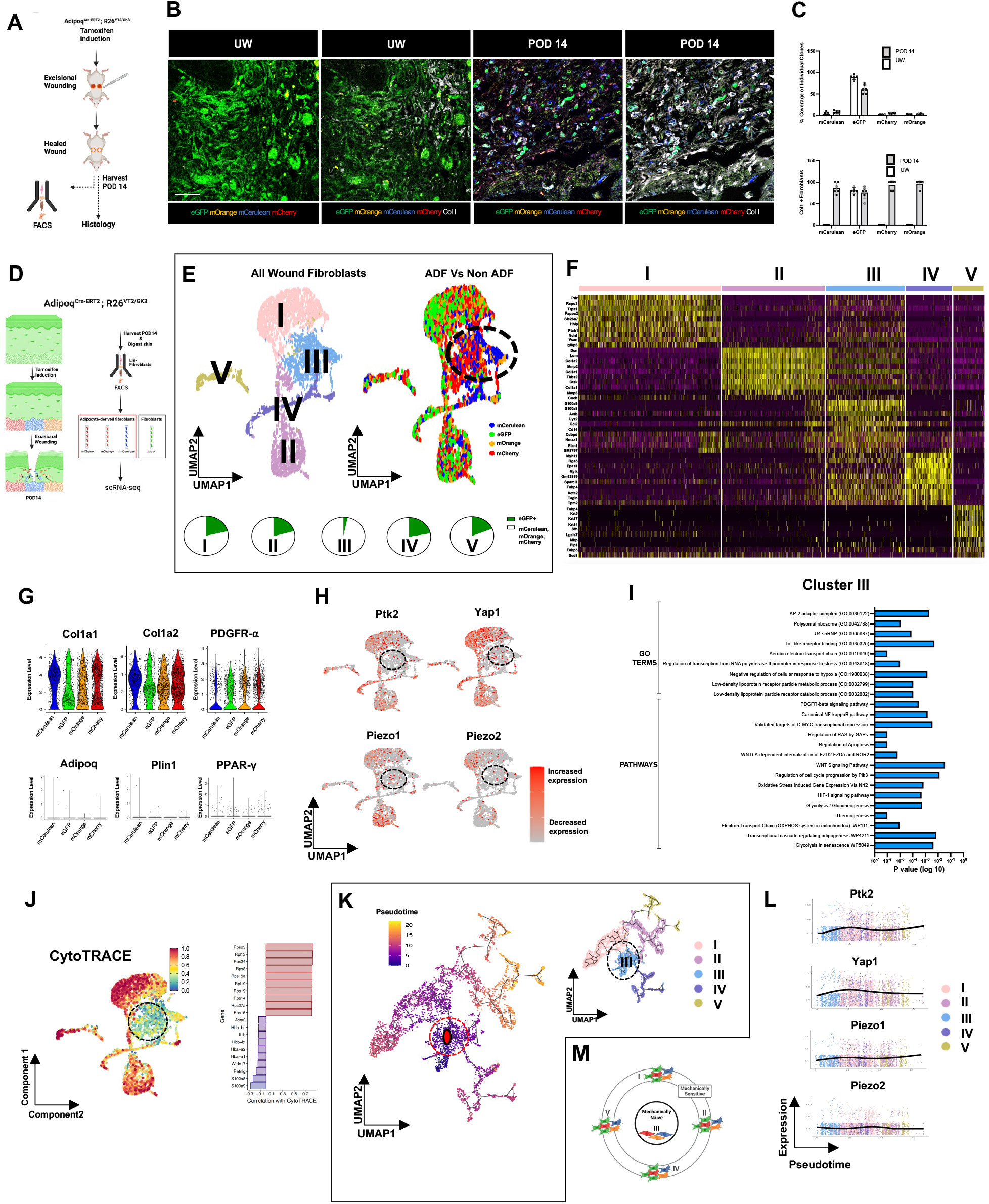
Clonal and single-cell transcriptomic analysis reveal differentiation dynamics of adipocyte-derived fibroblasts. (**A**) Schematic of *Adipoq^Cre-ERT^;R26^VT2/GK3^* (Rainbow) mouse wounding experiments for clonal analysis of adipocyte-derived cells. (**B**) Fluorescent histology with Col 1 IF staining (white signal) of Rainbow skin and postoperative day 14 (POD 14) wounds. eGFP, enhanced green fluorescent protein (green signal; Cre^-^/non-adipocyte lineage-derived cells in Rainbow construct); mOrange/mCerulean/mCherry, membrane-bound orange/cerulean/cherry fluorescent proteins (orange, blue, and red signals, respectively; collectively represent Cre^+^/adipocyte lineage-derived cells in Rainbow construct). Scale bar, 25µm. (**C**) Quantification of wound area percent coverage (top) and percentage of Col1^+^ fibroblast wound area (bottom) by Rainbow clone color in unwounded skin versus POD 14 wounds. (**D**) Schematic of Rainbow construct for analysis of adipocytes and their derivatives (left) and experimental approach for FACS-isolating and separately sequencing adipocyte-derived (mOrange^+^, mCerulean^+^, and mCherry^+^) and non-adipocyte-derived (eGFP^+^) fibroblasts (right). (**E**) Top, UMAP of Rainbow fibroblast scRNA-seq data, color coded by Seurat cluster (top left; overlaid Roman numerals I-V indicate Seurat cluster identity) or Rainbow clone color (top right). Bottom, relative representation of adipocyte-derived (mOrange, mCerulean, or mCherry^+^; collectively, white) versus non-adipocyte-derived (eGFP^+^; green) cells in each Seurat cluster. (**F**) Heatmap showing top differentially expressed genes for each Seurat fibroblast cluster. (**G**) Violin plots showing expression of genes characteristic to fibroblasts (top) or adipocytes (bottom) in fibroblasts of each Rainbow clone color. (**H**) Fibroblast scRNA-seq UMAP colored by expression level for known mechanical signaling genes. (**I**) Top GO pathway analysis results for genes characteristic to cells in Seurat cluster III (fibroblast cluster containing almost exclusively adipocyte-derived cells). (**J**) CytoTRACE analysis showing predicted order (differentiation potential scores; left) and genes with strongest positive and negative correlation with these scores (right). (**K**) Pseudotime analysis of Rainbow fibroblast scRNA-seq data; plots colored by relative pseudotime value (left) and Seurat cluster identity (right). Red point within cluster III (left plot) indicates root for pseudotime calculations. (**L**) Expression of mechanical signaling genes across pseudotime. Overlaid lines represent regression fit to gene expression of cells over pseudotime; colors indicate Seurat cluster identity of each cell. (**M**) Schematic depicting proposed identities of adipocyte-derived (red, orange, or blue) and non-adipocyte-derived (green) wound fibroblasts based on scRNA-seq analysis. We propose that Seurat cluster III represents a “mechanically naïve” population of ADFs that expresses lower levels of mechanical signaling genes and is transcriptomically more similar to adipocytes, whereas the other Seurat clusters (I, II, IV, and V) represent more typical “mechanically sensitive” wound fibroblasts, comprising both ADFs and non-adipocyte-derived wound fibroblasts, which highly express mechanical activation and fibrosis genes. (**E, H, J, K**) Dotted circled regions indicate Seurat cluster III. (**C**) Data shown as mean ± standard deviation (S.D.).

Next, to interrogate ADF heterogeneity and molecular signaling, we FACS-isolated fibroblasts expressing each Rainbow color, then subjected each population to single-cell RNA-sequencing (scRNA-seq) (**Fig. 2D and S5A**). Rainbow construct specificity was confirmed by the absence of reporter recombination without tamoxifen induction (**Fig. S5B**). Louvain-based (Seurat) clustering identified five transcriptionally distinct fibroblast clusters (denoted with Roman numerals I-V; **Fig. 2E-F and S5C**); notably, one of these clusters (cluster III) comprised almost exclusively adipocyte-derived fibroblasts (**Fig. 2E, bottom, and S5D**), suggesting that this cluster identity was unique to cells of adipocyte origin. Transcriptomic analysis revealed that fibroblasts, regardless of clone color, expressed negligible adipocyte genes and instead expressed typical wound fibroblast genes (**Fig. 2G and S5E**). We and others have previously implicated mechanical signaling (e.g., canonical focal adhesion kinase [FAK]/Yes-associated protein [YAP]-mediated mechanotransduction) as modulating wound fibrosis;(Chen et al., 2021b; Mascharak et al., 2021a; Mascharak et al., 2021b) interestingly, scRNA-seq revealed enrichment for FAK (Ptk2) and YAP (Yap1), as well as the mechanosensitive ion channel components Piezo1 and Piezo2,(He et al., 2018; Holt et al., 2021; Jin et al., 2020; Pethő et al., 2019) in all fibroblast clusters *except* cluster III (the cluster containing almost entirely adipocyte-derived fibroblasts; **Fig. 2H and S5F**). Further interrogation of cluster III using Gene Ontology (GO) pathway analysis revealed enrichment for non-traditional fibroblast pathways including lipoprotein particle receptor-related processes, thermogenesis, and adipogenesis (**Fig. 2I**). In stark contrast, all other fibroblast clusters were enriched for numerous “typical” fibrosis-related pathways, including mechanical activation terms (e.g., “focal adhesion,” “cell-matrix adhesion mediator activity”) and fibrosis-related terms (e.g., “collagen-containing extracellular matrix”; **Fig. S5G**). GeneTrail analysis further highlighted that Seurat fibroblast clusters I, II, IV, and V showed comparative upregulation of collagen processes, collagen-activated pathways, focal adhesion assembly, and leukocyte migration (**Fig. S5H**). Enrichment for typical fibrosis GO terms was observed across all Rainbow clone colors (**Fig. S5I**). One population in particular (cluster II) exhibited a transcriptional profile consistent with a mechanosensitive fibroblast subpopulation previously described by our group, which clonally proliferates toward the wound center following skin injury(Foster et al., 2021) (**Fig. S5J**).

We also analyzed cells using CytoTRACE (a computational method for predicting cell differentiation states based on relative transcriptional diversity(Gulati et al., 2020)), which revealed that cluster III represented a dramatically distinct differentiation state from all other fibroblast clusters (**Fig. 2J**). Pseudotime analysis supported the presence of clear differentiation trajectories originating in cluster III cells and progressing outward to all other clusters (**Fig. 2K and S5K**); differentiation along these trajectories was associated with increased expression of known mechanical signaling genes and pro-fibrotic markers (**Fig. 2L and S5L**). We then applied RNA velocity analysis (scVelo), which compares expression of unspliced pre-mRNA and mature spliced mRNA to infer directional information for transcriptional dynamics within the dermis.(Bergen et al., 2020) This approach again predicted differentiation trajectories originating from cluster III cells (**Fig. S5M-P**). RNA velocity analysis using root cell prediction and velocity length further supported that this differentiation trajectory starts from cluster III (**Fig. S5O**). Collectively, these results led us to conclude that cluster III may represent an intermediate differentiation state, in which adipocytes have transitioned to a fibroblast fate but remain “mechanically naïve” and may retain some hallmarks of adipocyte transcriptional identity; continued wound mechanical stimuli could then lead to further differentiation to a fully mechanically sensitive/activated and pro-fibrotic scar fibroblast state (**Fig. 2M**).

#### Mechanical forces are sufficient to drive adipocyte-to-fibroblast conversion in a process involving Piezo1 and Piezo2 signaling

We were interested to find transcriptional evidence for the relevance of mechanosignaling in ADFs. It is well known that tissue mechanical forces activate fibroblasts to drive scarring.(Mascharak et al., 2021b) However, while *in vitro* evidence suggests that adipocytes alter phenotype in response to mechanical environment,(Hossain et al., 2010; Yuan et al., 2015) whether mechanics modulate differentiation of wound adipocytes remains unknown. We sought to determine whether mechanical signaling could serve as a “switch” to guide adipocytes toward a fibrogenic (in response to mechanical strain) versus adipogenic (without strain or with blocked mechanotransduction) fate. As traditional floating adipocyte culture does not allow alterations of mechanical environment and fails to replicate cells’ native 3D environment, we adapted a published 3D culture system(Chen et al., 2021a) in which cell-seeded hydrogels undergo controlled stretching, permitting precise mechanomodulation, for mouse adipocytes (**Fig. 3A**, **top panel**). In the absence of applied stretch, mouse adipocytes had widespread expression of adiponectin and absent fibroblast markers; remarkably, when stretch was applied, a large proportion of adipocytes adopted a fibrogenic phenotype, losing adiponectin and instead expressing Col1 (**Fig. 3A**, **bottom panels**). When small molecule inhibitors were used to block key mechanosignaling genes (Piezo1 [P1i], Piezo2 [P2i], FAK [FAKi], or YAP [YAPi]; **Fig. S6A**), the stretch-induced phenotypic switch was abrogated (**Fig. 3A and S6B-E**). Collectively, these findings suggested that mechanical stress, communicated at the cellular level via mechanosensation pathways including *Piezo1* and *Piezo2*, was sufficient to drive adipocyte-to-fibroblast conversion.

**Figure 3:**
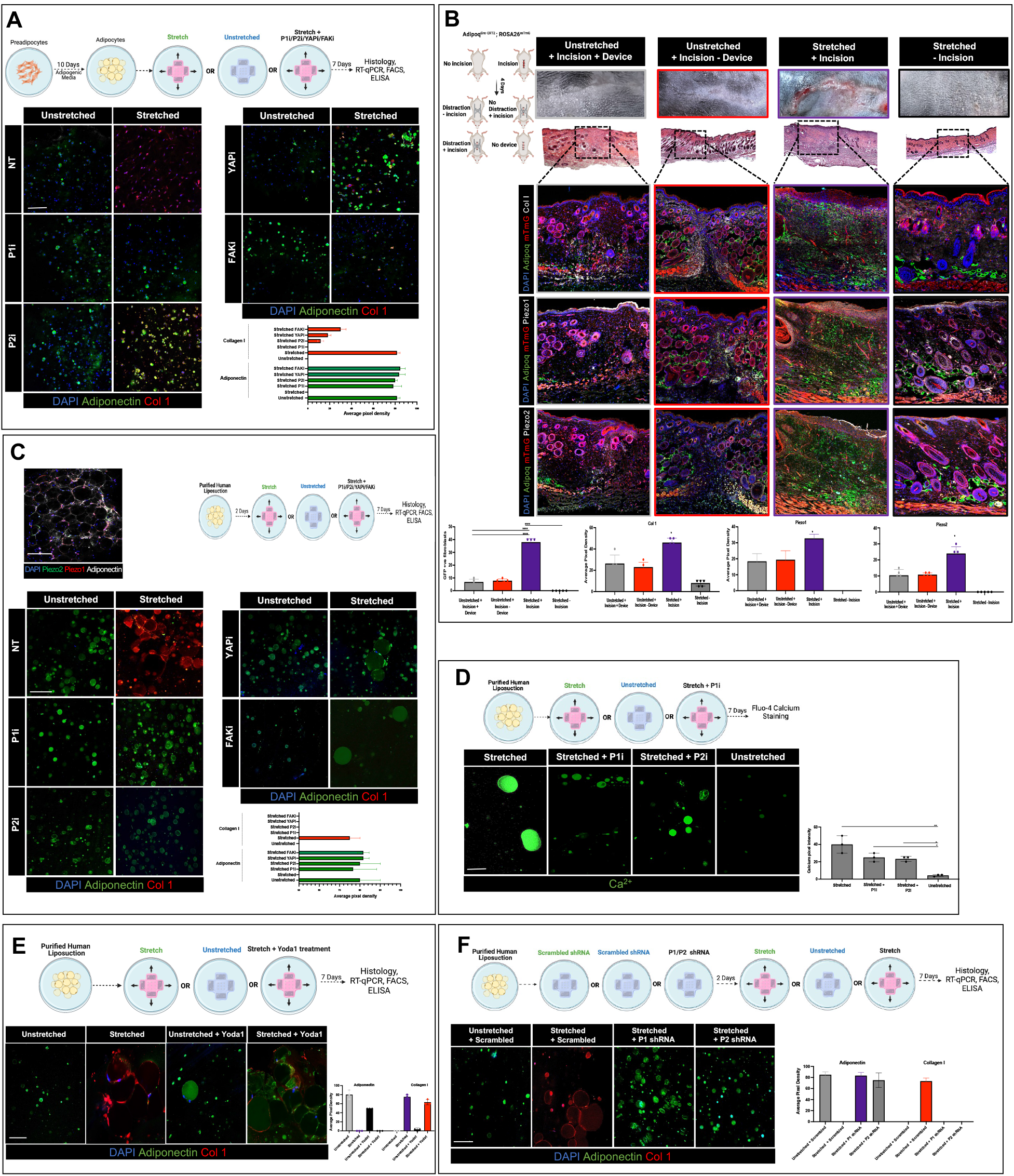
Adipocytes transition to fibroblasts in response to mechanical stimuli *in vitro* and *in vivo*. (**A**) Top, schematic of mouse adipocyte culture and mechanomodulation experiments. Bottom left, immunofluorescence (IF) staining of mouse adipocytes cultured with or without mechanical stretch, in the presence of indicated mechanosignaling inhibitors (Piezo1 inhibitor [P1i], Piezo2 inhibitor [P2i], YAP inhibitor [YAPi], and FAK inhibitor [FAKi]) or no inhibitor (no treatment, NT), with IF staining for adiponectin (green fluorescent signal) and Col1 (red signal). Bottom right, quantification of adiponectin and Col 1 expression from IF staining. (**B**) Top left, schematic of mouse hypertrophic scarring (HTS) model experiments with incisional wounding and mechanical distraction (stretching). Right panels: gross photographs (top row), H&E histology (second row; black dotted regions show wounded/stretched region of skin), fluorescent histology (third through fifth rows), and IF quantification (bottom row) of experimental conditions in mouse HTS model. Fluorescent histology shows Tomato background fluorescence (mTmG, red signal) and GFP from *Adipoq* lineage-derived cells (Adipoq, green signal) with IF staining for Col 1 (third row), Piezo1 (fourth row), or Piezo2 (fifth row; all IF staining, white signal). Bottom, quantification of ADFs (left panel) and Col 1, Piezo1, and Piezo2 expression (right three panels; *\*P* ≤ 0.05 vs. all other conditions) from IF staining. (**C**) Top left, whole mount histology of human adipose tissue with IF staining for Piezo1 (red signal), Piezo2 (green), and adiponectin (white). Top right, schematic of human adipocyte culture and mechanomodulation experiments. Bottom, as in (A) bottom, but with human adipocytes. (**D**) Top, schematic of human adipocyte culture to evaluate cell calcium content. Bottom left, IF staining of human adipocytes cultured in indicated conditions showing calcium staining (green signal). Bottom right, quantification of calcium staining. (**E**) Schematic (top), IF staining (as in (**A**) and (**C**); bottom left), and IF quantification (bottom right) of human adipocyte culture with Yoda1 (Piezo1 agonist). (**F**) As in (E), but with Piezo1 (P1 shRNA), Piezo2 (P2 shRNA), or scrambled shRNA (Scrambled). Data shown as mean ± S.D. (B, D) *\*P* ≤ 0.05, \*\**P* ≤ 0.01, *\*\*\*\*P* ≤ 0.0001. Scale bars, 100 µm (**A**), 150 µm (**B**, second row), 200 µm (**B**, third through fifth rows), 100 µm (**C-F**).

It was next critical to confirm these findings *in vivo*. We applied a model that subjects mouse incisional wounds to increased tension, producing scars that resemble human hypertrophic scars (HTS),(Aarabi et al., 2007) in *Adipoq^Cre-ERT^;R26^mTmG^* mice (**Fig. 3B**, **top left**) to directly determine whether mechanical loading promotes adipocyte-to-fibroblast conversion in wounds *in vivo*. When wounds were mechanically stretched, scars were more fibrotic and had substantially increased ADFs, Col1, and mechanical signaling factors including Piezo1 and Piezo2 (**Fig. 3B and S6F-G**).

Given that significant differences may exist between mouse and human adipocytes, we sought to validate our mouse findings in human cells. IF staining revealed that Piezo1 and Piezo2 were expressed in human adipocytes while, interestingly, FAK and YAP were minimally expressed (**Fig. 3C**, **top left, and S7A**). We applied our mechanomodulatory culture system to human adipocytes (**Fig. 3C, top right**) and found that, similar to mouse adipocytes, stretching caused these cells to express Col1 and lose adipocyte markers; this transition was again abrogated by inhibiting mechanosignaling (**Fig. 3C, bottom, and S7B-E**). Interestingly, stretching also induced substantial hypertrophy of human adipocytes (previously linked with pathological remodeling/fibrosis of adipose tissue(Vishvanath and Gupta, 2019)), which was reversed with P1i and P2i but not FAKi or YAPi (**Fig. 3C, bottom**). Consistent with stretching driving activation of calcium-permeable Piezo mechanosensitive ion channels,(Matsunaga et al., 2021; Romac et al., 2018) we observed increased calcium staining with stretching that decreased with P1i or P2i (**Fig. 3D**). Our inhibitor experiments suggested that mechanosignaling was necessary for adipocyte-fibroblast transition in response to stretch; conversely, applying Yoda1 (a Piezo1 agonist(Syeda et al., 2015)) was sufficient to drive adipocyte hypertrophy even in the absence of applied stretch (**Fig. 3E and S7F-G**). Given possible off-target effects of small molecules, we also used short hairpin RNA (shRNA) to knockdown Piezo1 and Piezo2 (**Fig. 3F**, **top, and S7H**), which blocked stretch-induced adipocyte hypertrophy and adipocyte-to-fibroblast transition (**Fig. 3F and S7H- J**).

### Blocking Piezo mechanosignaling in adipocytes reduces scarring and fibrosis

Given the dramatic effects of Piezo1 and Piezo2 inhibition on adipocyte differentiation into fibroblasts *in vitro*, we next examined whether blocking adipocyte mechanosensing *in vivo* could prevent adipocyte-to-fibroblast transition in wounds and yield reduced scarring. We focused on Piezo1 and Piezo2 because, compared to FAK or YAP, their expression was more adipocyte-specific (**Fig. S7A**) and their inhibition reversed both hypertrophy and gene expression changes in stretched adipocytes, compared to gene expression only with FAK/YAP (**Fig. 3C**). Further, genetic lineage tracing and *in situ* hybridization confirmed that Piezo1 and Piezo2 expressing cells were present and lost adipocyte/gained fibroblast markers in wounds (**Fig. S8A-D**).

We treated wounds in tamoxifen-induced *Adipoq^Cre-ERT^;R26^VT2/GK3^* mice with P1i or P2i or PBS (vehicle control) via local injection into the wound edges at POD 0 (**Fig. 4A**). Doses were optimized based on *in vitro* dosing and confirmed with toxicity testing (**Fig. S6H-I**). P1i and P2i did not significantly affect time to complete wound re-epithelialization (**Fig. S9A**). Grossly and histologically, Piezo inhibitor-treated wounds yielded less-apparent scars, with significantly decreased scar thickness, increased adipocytes, and (particularly in P1i wounds) numerous structures morphologically consistent with unerupted hair follicles (HF; **Fig. 4B**). While control wounds had ECM distinct from unwounded skin’s (consistent with fibrotic scars), Piezo-inhibited wounds had ECM features overlapping unwounded skin’s (**Fig. 4C**), suggesting that mechanotransduction inhibition led to regeneration of normal skin-like ECM. ADFs were significantly reduced in P2i and absent in P1i wounds (**Fig. 4D-E and S9B, top**), suggesting that Piezo inhibition prevented adipocyte-to-fibroblast conversion. P1i and P2i wounds had reduced Col1 and *α*-SMA and less dense ECM at both POD 14 and POD 30 (early remodeling phase of wound repair), while Oil Red O staining and IF for cytokeratins (CK) 14/19 (markers of regenerating appendages) demonstrated presence of putative regenerating sebaceous glands/HF (**Fig. 4E-F and S9B-D**). Congruent with our previous study(Mascharak et al., 2021a), where we found YAPi inhibition to allow for hair regrowth at POD 30, P1i/P2i allowed for hair follicle formation at POD 30 (**Fig. S9E)**. Finally, mechanical testing confirmed that P1i (and, to a lesser extent, P2i) reverted scars to more unwounded-like mechanical properties (**Fig. 4G**).

**Figure 4:**
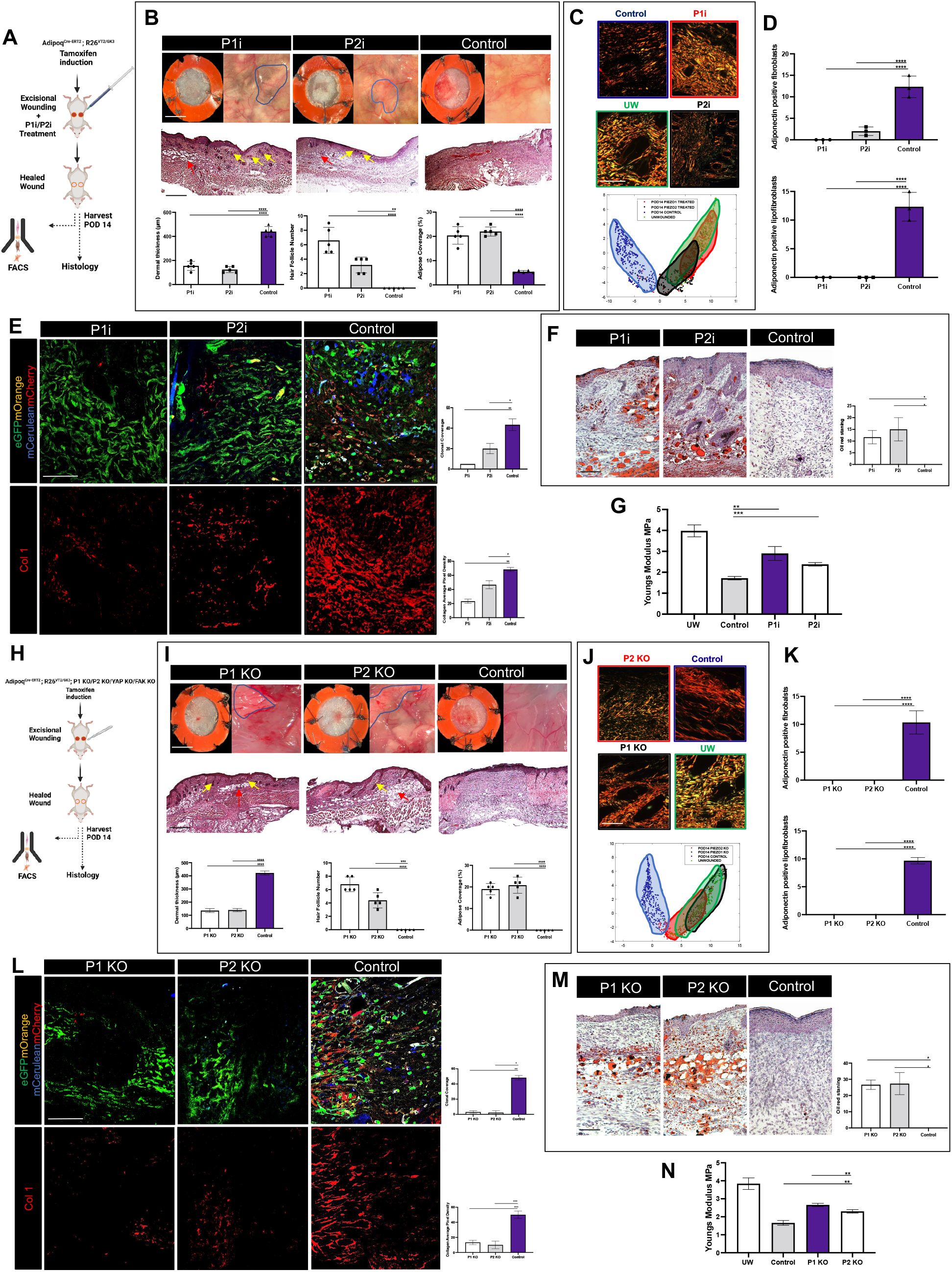
Adipocyte-targeted mechanosignaling blockade in wounds via small molecule inhibition and genetic knockout of Piezo1 (P1i) and Piezo2 (P2i) reduces scarring and fibrosis. (A) Schematic of *Adipoq^Cre-ERT^;R26^VT2/GK3^* wounding with small molecule P1i or P2i treatment. (B) Gross photos (top row; paired photos of top [left image] and underside [right image] of wounds) and H&E histology (middle row) of wounds treated with P1i, P2i, or control (PBS). Blue outlined regions and red arrows, fat accumulation; yellow arrows, regenerating secondary elements (e.g., hair follicles [HFs]). Bottom row, quantification of dermal thickness (left), number of HFs in wounded area (middle), or adipose coverage as percent of wound area (right) for each treatment condition. (C) Picrosirius red histology (top) and UMAP of quantified extracellular matrix (ECM) ultrastructure parameters (bottom; each dot represents one histology image) for indicated conditions. (D) Quantification of percentage of fibroblasts (top) or lipofibroblasts (bottom) that are *Adipoq* lineage-derived (eGFP^-^ and mOrange^+^, mCerulean^+^, or mCherry^+^) by fluorescence-activated cell sorting (FACS). (E) Left, fluorescent histology of wounds showing Rainbow clone colors (top; eGFP, green signal; mOrange, orange signal; mCerulean, blue signal; mCherry, red signal) or Col 1 immunofluorescence (IF) staining (bottom, red signal). Right, quantified percent wound area coverage by adipocyte-derived clones (top) and Col 1 expression by IF (bottom). (F) Left, Oil Red O staining (red) for lipid/oil glands (left); right, quantification of red staining density. (G) Young’s modulus, calculated from tensile strength testing, by wound condition. (H) Schematic depicting mouse wounding for transgenic, adipocyte-targeted KO of mechanosignaling genes. (I-N) As in (B-G), but with genetic P1 or P2 Knockout (KO) versus control (P1/P2 wildtype). Data shown as mean ± S.D. *\*P* ≤ 0.05, \*\**P* ≤ 0.01, *\*\*\*P* ≤ 0.001, *\*\*\*\*P* ≤ 0.0001. Scale bars, (A) 3mm 250 µm (B, I), 20 µm (C, J), 25 µm (E, F, L, M).

While these results suggested that P1i or P2i was sufficient to prevent adipocyte-to-fibroblast conversion and reduce wound fibrosis, small molecules may have off-target effects. Thus, we developed *Adipoq^Cre-ERT^;R26^VT2/GK3^;Piezo1^fl/+^* (*Piezo1^fl/+^*) and *Adipoq^Cre- ERT^;R26^VT2/GK3^;Piezo2^fl/+^* (*Piezo2^fl/+^*) mice and administered tamoxifen to induce both Rainbow reporter combination and Piezo1 or Piezo2 deletion in adipocytes prior to splinted excisional wounding; we similarly generated adipocyte-targeted YAP and FAK KO mice to compare the efficacy of ablating these canonical mechanotransduction molecules in adipocytes (**Fig. 4H**). None of these genes’ (P1, P2, YAP, FAK) deletion in adipocytes significantly affected time to wound closure (**Fig. S10A**). Similar to the effects of P1i and P2i, Piezo1 and Piezo2 KO yielded significantly reduced scarring with fat accumulation and regeneration of dermal appendages in wounds (**Fig. 4I-M**). Notably, tensile testing revealed that Piezo KO wounds had mechanical properties similar to those of unwounded skin rather than scars (**Fig. 4N**). Supporting our *in vitro* evidence suggesting that YAP and FAK inhibition were less effective at preventing mechanically-activated adipocyte changes compared to Piezo inhibition, fibrosis was mitigated to a much lesser extent in adipocyte-targeted YAP and FAK KO wounds (**Fig. S10B-F**). Collectively, these results demonstrated that adipocyte-specific Piezo1 or Piezo2 blockade prevented adipocyte-to-fibroblast conversion and had anti-scarring/pro-regenerative effects in wound healing.

### Piezo blockade prevents activation of “mechanically naïve” to “mechanically activated” adipocyte-derived fibroblasts

In order to more deeply interrogate the mechanism(s) by which Piezo inhibition prevented scarring and induced wound regeneration, we performed scRNA-seq of cells from five wound healing conditions (all wounds at POD 14): wounds treated with small molecule Piezo inhibitors (P1i or P2i); wounds in Piezo1 KO mice; control wounds (wildtype mice without treatment); or unwounded skin (**Fig. 5A**). Sequenced cells included all expected wound cell types, including fibroblasts, keratinocytes, endothelial cells, inflammatory cells, and a small number of adipocytes (**Fig. 5B and S11A-C**). Following sequencing, we performed *in silico* selection for fibroblasts (based on individual sequenced cell transcriptomic profiles; **Fig. 5B**); we confirmed that these cells expressed canonical fibroblast genes and had minimal expression of adipocyte identity genes (**Fig. S11B**). Six transcriptionally distinct fibroblast clusters (designated clusters 0-5) were identified by Louvain-based clustering (Seurat; **Fig. 5C-D**). To determine how these populations related to those identified by our original scRNA-seq fibroblast dataset (**Fig. 2E**), we applied an anchor-based label transfer approach to project the current six clusters (**Fig. 5C**) onto the original five scRNA-seq clusters (**Fig. 2E**) in a K-nearest neighbor-based fashion (**Fig. 5E**, **top**).(Stuart et al., 2019) These projections supported strong similarities between each cluster from the original dataset and specific clusters in the current dataset, most notably between our previous cluster III (the “mechanically naïve” subpopulation from **Fig. 2**) and new cluster 5 (**Fig. 5E**). Supporting that the newly-identified cluster 5 shared the unique transcriptional program previously identified as the “mechanically naïve” fibroblast cluster in **Fig. 2**, GO pathway analysis of cluster 5 revealed terms related to lipoproteins, visceral fat, and lipid functions; in contrast, all other clusters were enriched for typical pro-fibrotic fibroblast terms such as focal adhesion and ECM-related terms as well as mechanical signaling genes (**Fig. 5F and S11D**). Notably, cluster 5 was less represented in control (scarring) wounds and more highly represented in Piezo inhibition and KO wound cells as well as unwounded skin (**Fig. 5G and S11E**); P1i wounds in particular also contained relatively more adipocytes and fewer fibroblasts (**Fig. 5H**). Cluster 5 was distinguished by lower expression of key mechanical signaling genes (**Fig. S11F**), consistent with its putatively “mechanically naïve” identity. Using RNA velocity analysis, we again identified a differentiation trajectory originating from mechanically naïve cells (cluster 5; **Fig. 5I and S11G-I**) which we then used as a starting point for pseudotime analysis (**Fig. 5J and S11J-K**). Both analyses further supported cellular transition from “mechanically naïve” fibroblasts to more traditional fibroblast subpopulations (**Fig. S11G-I**). Overall, these results were again consistent with the presence of two distinct differentiation states of adipocyte-derived fibroblasts: a mechanically naïve population (cluster 5), which is enriched in states of inhibited adipocyte mechanotransduction; and a more mechanically primed population, which is relatively enriched in typical scarring wound conditions and is more fibrotic (**Fig. 5K**).

**Figure 5:**
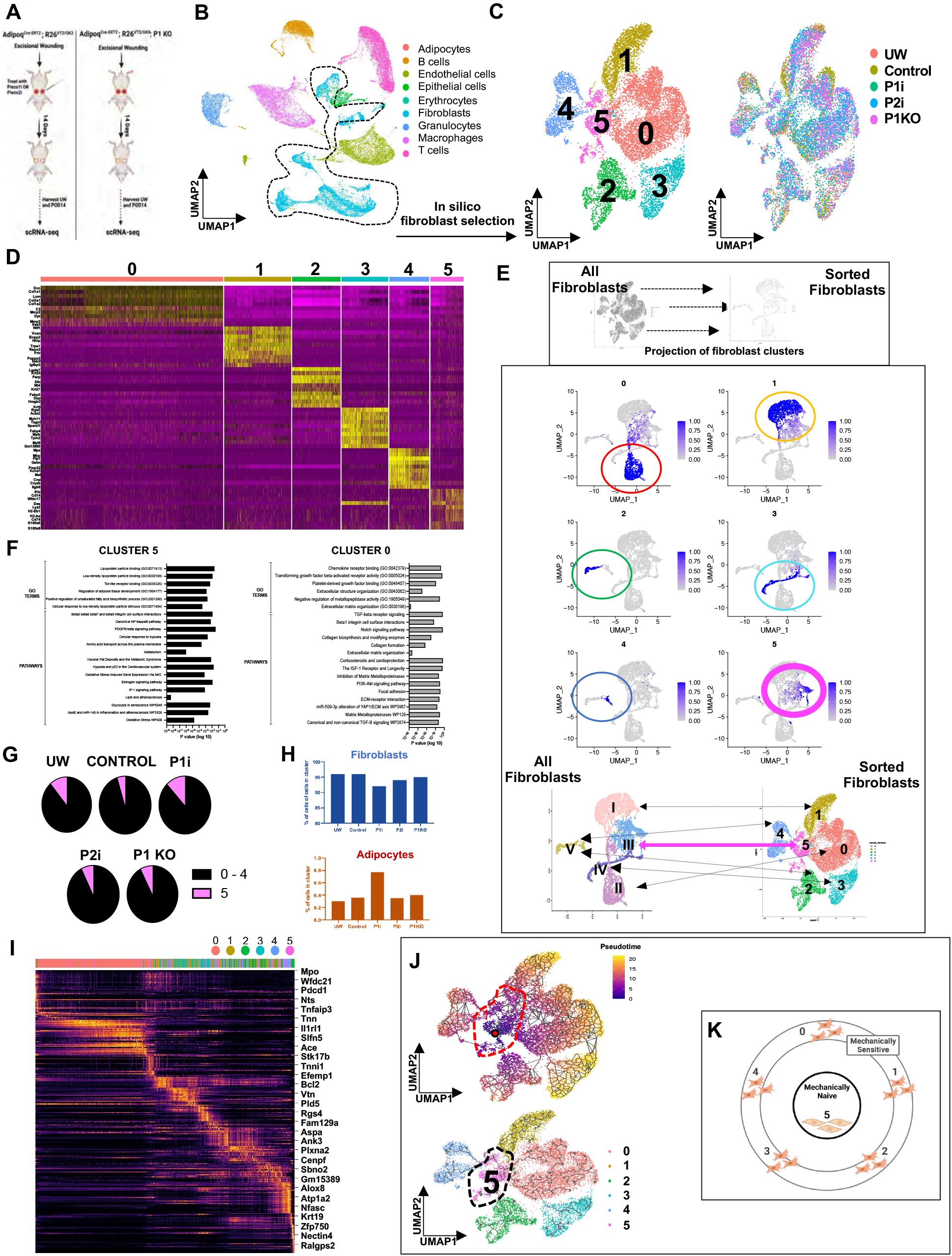
Adipocyte mechanosignaling (Piezo) blockade influences differentiation dynamics of “mechanically naïve” versus “mechanically sensitive” adipocyte-derived fibroblasts. (A) Schematic of wounding experiments with Piezo1 inhibitor (P1i), Piezo2 inhibitor (P2i), or Piezo1 knockout (P1 KO) and scRNA-seq analysis. (B) UMAP of scRNA-seq data from all wound cells, colored by cell type. Black dotted region indicates cells transcriptomically classified as fibroblasts (i.e., *in silico* selection) that were used for downstream analysis. (C) UMAP plots of fibroblasts colored by either Seurat cluster (0-5; left) or experimental condition (right). (D) Heatmap showing top differentially expressed genes for each Seurat fibroblast cluster. (E) Top, schematic depicting approach for projecting the six clusters derived from the current dataset (left) on the original five scRNA-seq clusters from Fig. 2 (right). Center, label transfer projection of Fig. 5C fibroblast subpopulations onto Fig. 2 embedding. Colored circles highlight the spatial locations of Fig. 5C subpopulations. Bottom, summary of correlations between fibroblast clusters from Fig. 2 to Fig. 5C; arrows indicate Seurat cluster from original dataset that most closely corresponds to each cluster in new dataset (heavy pink arrow indicates correlation between Fig. 2 cluster III and Fig. 5 cluster 5). (F) Gene Ontology (GO) pathway analysis for indicated Seurat clusters in (all wound conditions) scRNA-seq dataset. (G) Relative representation of fibroblasts belonging to clusters 0- 4 versus cluster 5 from each experimental condition. (H) Quantification of percentage of all sequenced wound cells from each condition that were considered fibroblasts (top) or adipocytes (bottom) based on transcriptomic profiles. (I) scVelo heatmap highlighting genes with high correlation with velocity pseudotime, indexed by Seurat cluster. (J) Top, pseudotime analysis of fibroblasts from all wound conditions, with plots colored by relative pseudotime value (top) and Seurat cluster identity (bottom). (K) Schematic depicting proposed identities of “mechanically naïve” (cluster 5) versus typical “mechanically sensitive” (clusters 0-4) wound fibroblasts based on scRNA-seq analysis.

Finally, we analyzed our scRNA-seq data using CellChat, a computational method for inferring cell-cell interactions(Jin et al., 2021) that we previously applied to study dynamics of wound fibrosis versus regeneration. Examining all cell-cell interactions revealed fewer overall interactions with cluster 5 (“mechanically naïve”) fibroblasts compared to other fibroblast clusters (especially cluster 0) (**Fig. S12A-B**). Extensive cell signaling was found between fibroblasts and adipocytes, including a unique pattern of signaling between cluster 5 fibroblasts and adipocytes that was not present with other fibroblast clusters (**Fig. S12C**). Following P1i treatment, we identified an increase in signaling between cluster 5 fibroblasts and most other cell types, compared to decreased signaling of other fibroblast clusters (**Fig. S12D-E**). P1i treatment also enhanced signaling of known adipocyte pathways including Leptin (Lep),(Harris, 2014) Chemerin,(Goralski et al., 2007) and PD-L1(Ingram et al., 2017) (**Fig. S12F**). Lastly, P1i treatment enhanced interactions between cluster 5 fibroblasts and adipocytes (**Fig. S12G**), suggesting that Piezo inhibition alters crosstalk between adipocytes and fibroblasts upon wounding.

### Piezo inhibition reduces fibrosis in existing mouse scars

While wound re-epithelialization is completed relatively quickly (around 2 weeks), wounds continue to heal beneath the surface for months to years in a process known as scar remodeling.(Foster et al., 2021; Gurtner et al., 2008) Remodeling is the longest and least well understood phase of wound repair but is likely critical to ultimate outcomes. We postulated that ADFs could continue to play a role in scarring during the remodeling process; if this were the case, we wondered whether ADF activity could be targeted during wound remodeling, via P1i in existing, actively remodeling scars, to ultimately reduce scarring and drive regeneration (**Fig. 6A, C**). Thus, we administered P1i to existing scars at POD 30 (early remodeling period), POD 75 (mid- remodeling), and POD 120 (late remodeling), then harvested tissue one month later at POD 60, POD 105, and 150 respectively (**Fig. 6A, C**). Remarkably, treatment with P1i was sufficient to induce near-complete wound regeneration by POD 60, POD 105, and POD 150 with return of hair follicles (**Fig. 6B, D**), including full recovery of unwounded-like ECM architecture, compared to untreated wounds which remained scar-like on histology (**Fig. 6E-F**, **S13A-C, S14-S16**). Of note, the hair follicles at POD 105 and 150 were less organized than those at POD 60 following P1i treatment suggesting the regeneration is less complete (**Fig. 6E, S13A**). ADFs were significantly reduced in late P1i-treated wounds (**Fig. S13D-H**), suggesting that P1i was still acting on ADFs even when given at this later timepoint. Treatment of POD 30 scars, with YAPi did not induce regeneration suggesting that adipocytes play a more significant role in the remodeling phase of wound healing than fibroblasts (**S17A-C**). Overall, these findings suggested that ADFs continue to play a role during the remodeling phase, and that P1i could be effective to target ADFs and thereby promote regeneration even in existing scars.

**Figure 6:**
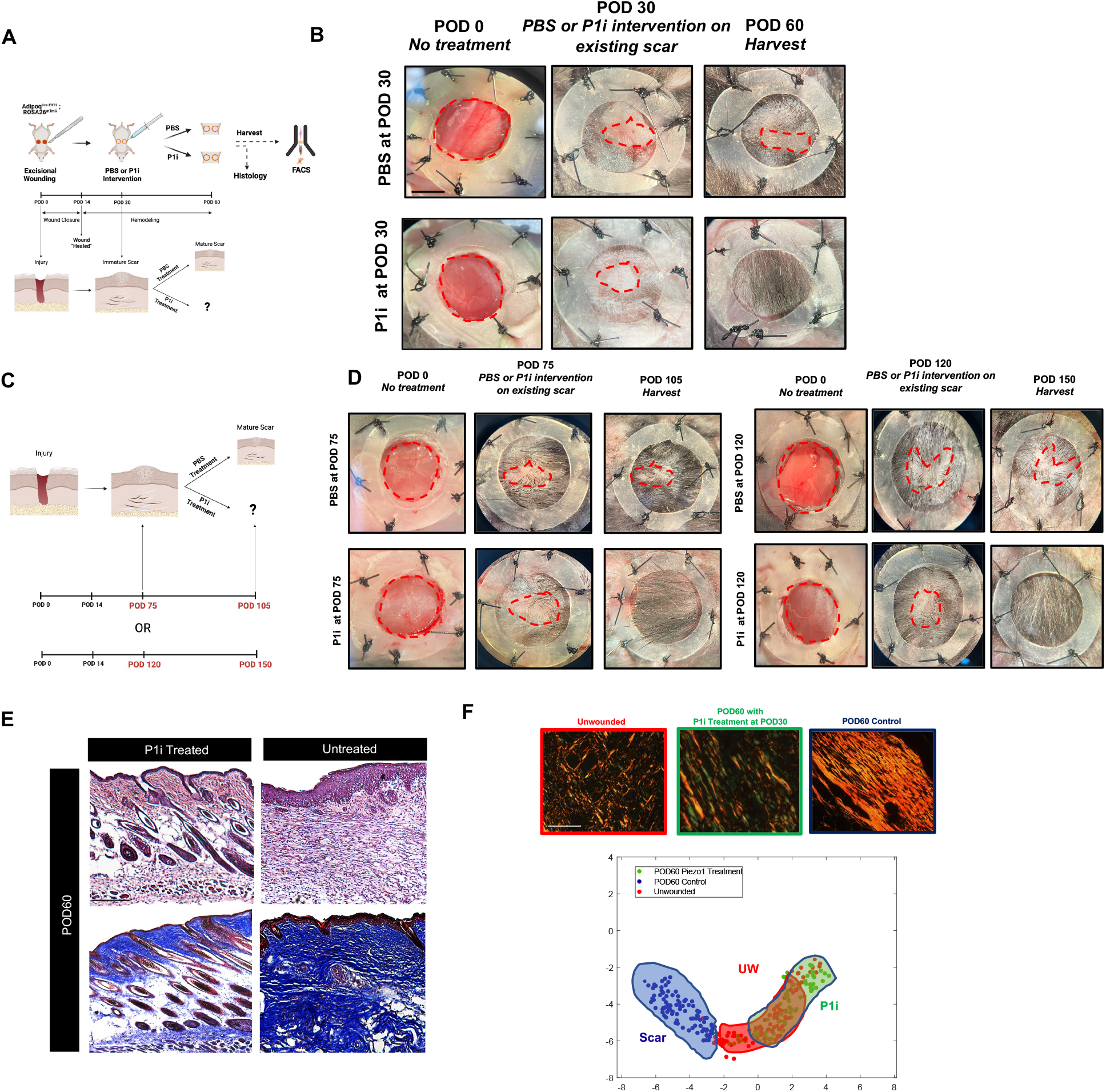
Piezo1 inhibition during wound remodeling reduces fibrosis in existing mouse scars. (**A**) Schematic, treatment of existing scars with Piezo1 inhibitor (P1i) during early remodeling phase at Post-operative day (POD) 30 with harvest at POD 60. (**B**) Gross photographs of wounds at POD 0, 30, and 60. (**C**) Schematic, treatment of existing scars with Piezo1 inhibitor (P1i) during mid remodeling phase at POD 75 with harvest at POD 105 and late remodeling at POD 120 with harvest at POD 150. (**D**) Gross photographs of wounds at POD 0, 75, and 105 (left) and POD 0, 120, and 150 (right) following PBS (top) or P1i treatment (bottom). (**E**) H&E (top) and trichrome (bottom) staining of POD 60 wounds treated with P1i (left) or control (untreated; right) at POD 30. (**F**) Top, picrosirius red staining of unwounded (UW) skin and POD 60 wounds treated at POD 30 with P1i or PBS (control). Bottom, UMAP of quantified extracellular matrix (ECM) ultrastructure parameters based on picrosirius red histology (each dot represents one histologic image). Data shown as mean ± S.D. *\*P* ≤ 0.05. Scale bars, 3mm (**A**) 150 µm (**F**), 20 µm.

### Spatial transcriptomics defines mechanically sensitive fibroblasts in wound healing

To further explore the significance of ADFs in wound repair, we applied the 10x Genomics Visium platform to analyze gene expression in the context of the spatial environment. We performed spatial transcriptomic analysis on histological sections from wounds at POD 14 (re-epithelized) with and without P1i/P2i, as well as unwounded skin (**Fig. 7A**). The epidermal, dermal, and hypodermal layers were clearly identified both histologically and according to their gene expression profiles (**Fig. 7B**). Furthermore, analysis of individual genes for well-known wound healing cell types, showed a clear distinction of keratinocytes in the epidermis (Krt14), fibroblasts and immune cells in the dermis (Col1a1 and Ptprc, respectively), and adipocytes in the hypodermis (Adipoq) as shown in wounds treated with PBS (**Fig. S18A**). POD 7 wounds were also easily histologically delineated according to the epidermis, dermis, and hypodermis (**Fig. S19A**).

**Figure 7:**
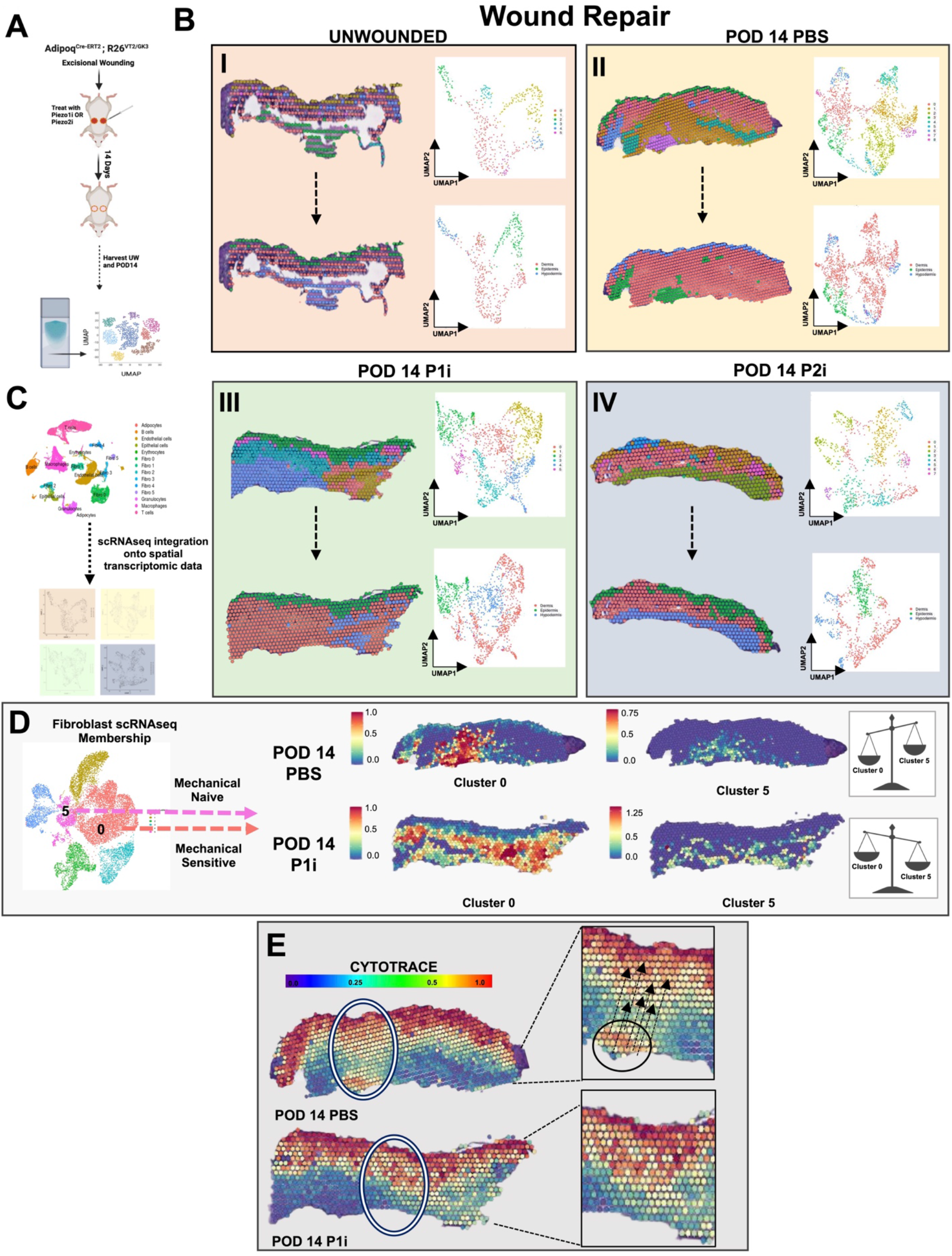
Visium analysis reveals that piezo inhibition during wound healing alters the spatial transcriptional. (**A**) Schematic for generating spatial transcriptomic data from splinted excisional wounds using the 10x Genomic protocol. (**B**) **[I-IV]** Delineation of scar layers based on Seurat Clusters (top left) with UMAP plot (top right) and underlying tissue histology (bottom left) with UMAP plot (bottom right) showing that the three scar layers can easily be distinguished by their transcriptional programs. Arrows the same tissue sample just colored by either Seurat Clusters or histological layer. (**C**) Schematic showing the anchor-based integration of scRNA-seq populations (defined in Fig.5) with Visium gene expression to project partial membership within each spot across all groups. (**D**) Spatial plots showing spatial expression of fibroblast cluster 0 (mechanically sensitive) and cluster 5 (mechanically naïve) in PBS (top) and P1i-treated wounds (bottom). (**E**) CytoTRACE analysis of PBS (top) and P1i- treated wounds (bottom).

To further elucidate the role of the mechanosensitive fibroblast population identified through scRNA-seq (**Fig. 5**), we evaluated the spatial expression of markers defining this population including Piezo1, Piezo2, Ptk2, and Yap1. In PBS treated wounds Piezo1, Piezo2, Ptk2, and Yap1 were found at the apical aspect of the wound, compared to P1i treated wounds where they appeared mostly at the basal region (**Fig. S18B**). A similar spatial pattern was observed at POD7 in PBS and P1i treated wounds (**Fig. S19B**). These data suggest that the genes defining the mechanosensitive fibroblast population (Piezo1, Piezo2, Ptk2, and Yap1) are spatially distinct in wounds treated with PBS versus those treated with P1i/P2i.

Analysis of spatially variable features across PBS and P1i/P2i treated wounds further illustrated the pro-fibrotic phenotype of PBS-treated wounds. The PBS treated wounds showed high expression of genes involved in scarring, including Cola1a1, Col1a2, and Acta1 (**Fig. S20A**). In comparison, P1i/P2i treated wounds revealed high expression of markers associated with the epidermis including Krt1 and Krt14 (**Fig. S20A**).

To ascertain the spatial pattern of scRNA-seq defined populations at POD 14 (**Fig. 5**), we performed anchor-based integration to assess membership of each of the six scRNA defined fibroblast clusters (**Fig. 7C**). We found that the predicted spatial distributions for our scRNA-seq clusters were largely congruent with our observed differences in the distribution of mechanically sensitive and naïve populations (**Fig. 7D, left**). Fibroblast scRNA-seq cluster 0, our mechanically sensitive population, was highly enriched in PBS- treated POD 14 wounds compared to P1i or P2i- treated wounds (**Fig. 7D, right top**). In comparison, fibroblast scRNA-seq cluster 5, our mechanically naive population was observed in P1i/P2i treated wounds (**Fig. 7D, right bottom**). Furthermore, adipocyte and epithelial cells were enriched in the spatial environment of P1i and P2i-treated wounds compared to PBS treated wounds at POD 14 (**Fig. S20B**). Lastly, to assess the relative differentiation states of the cells in the wound given a spatial context, we applied CytoTRACE to the four wound groups (**Fig. 7E**). We found greater transcriptional diversity from the apex compared to the basal dermis in a PBS treated scar, compared to wounds treated with P1i/P2i.

Collectively, these data suggest that mechanical inhibition through P1i or P2i can alter the spatial phenotype of adipocytes and fibroblasts during wound repair. Furthermore, the spatial pattern of ADFs and non-ADF subpopulations are distinct in wounds with regenerative and scarring phenotypes.

### Spatial phenotyping by CODEX demonstrates adipocyte-fibroblast cross-talk in wound healing

Building upon the spatial transcriptomic analyses described above, we further spatially phenotyped P1i-, P2i- and PBS-treated wounds at POD-14 (re-epithelized wounds) in a spatially-informed fashion at the protein level. We utilized CO-Detection by indEXing (CODEX), an assay in which a panel of 28+ individual protein markers are sequentially labeled with, and iteratively imaged via cyclic additions and washouts of dye-labeled oligonucleotide-conjugated antibodies (**Fig. 8A**; **markers shown in Table 1, left column**). Following denoising, image normalization, and automated cell segmentation, protein staining profiles were used to project a manifold of cell-representative clusters (**Fig. S21A and Fig. 8B**). Across all treatment groups, 19 cell clusters were identified on the basis of CODEX protein expression signatures, including subpopulations of six fibroblasts; three epithelial cells; five immune cells; two adipocytes, smooth muscle cells, endothelial cells, and pericytes (**Fig. 8B-C**). Broadly the expression of markers associated with mechanical signaling (PIEZO1, PIEZO2, YAP, and PTK2) was similar between P2i and P1i-treated wounds and unwounded skin, suggesting similar spatial protein expression (**Fig. 8D**). Notably, new hair follicle formation marked by Lef1 was highly expressed in P1i compared to PBS-treated wounds (**Fig. 8D**).

**Figure 8:**
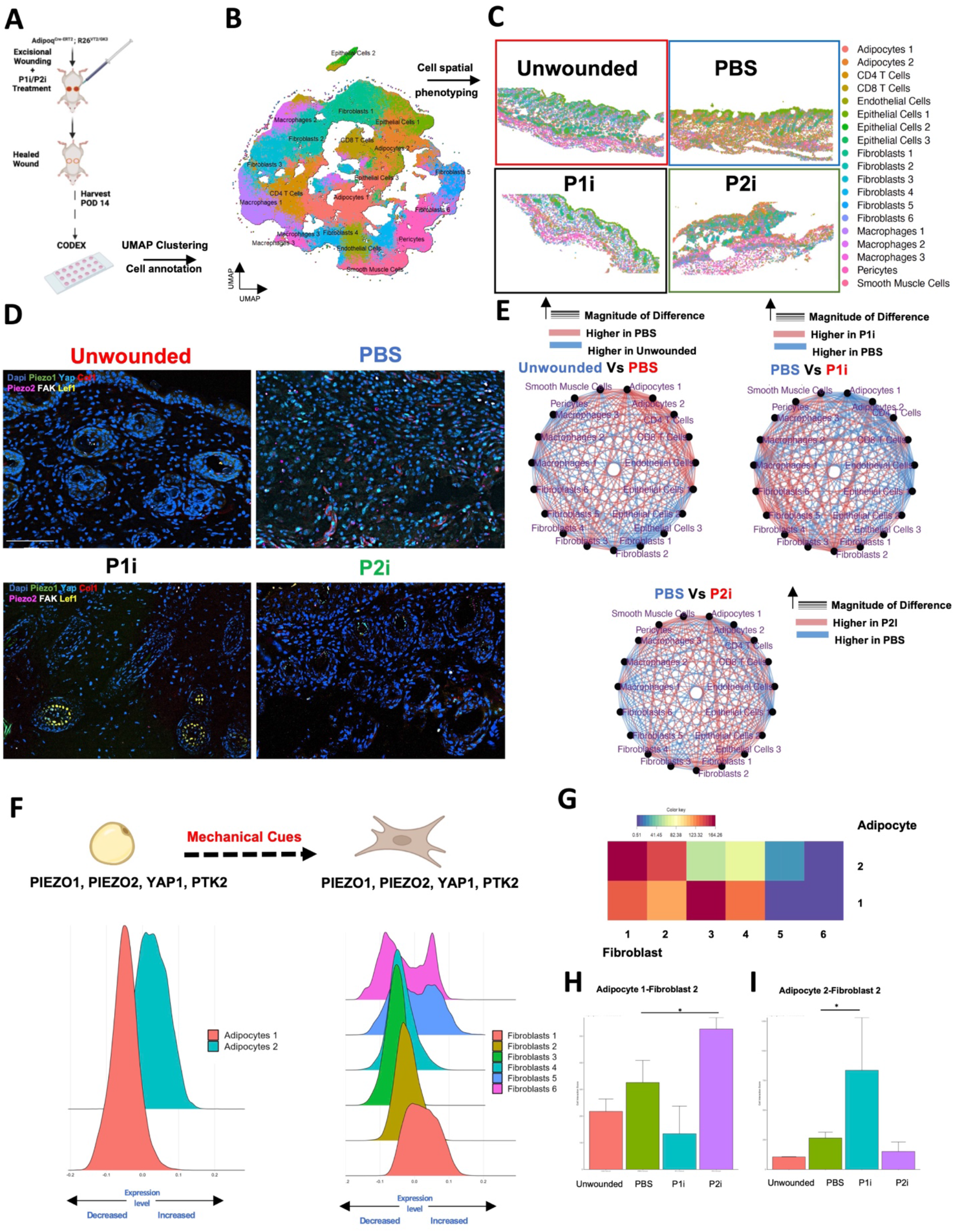
CODEX analysis reveals that Piezo inhibition during wound healing alters the spatial cross-talk between adipocytes and fibroblasts. (A) Schematic of the CODEX experiment strategy. (B) UMAP plot of CODEX data for all sequenced cells. (C) Representative images of identified UMAP CODEX clusters showing their spatial distribution in unwounded skin and PBS- and P1i/P2i-treated wounds. (D) Representative images of six selected CODEX markers in unwounded skin and PBS- and P1i/P2i-treated wounds. (E) Differential interaction maps in unwounded skin vs. PBS-treated wounds (top left), PBS-treated wounds vs. P1i treated wounds (top right), and PBS-treated vs P2i treated wounds (bottom). (F) Histograms of mechanosensitive protein module expression (PIEZO1, PIEZO2, YAP1, and PTK2 in adipocytes (left) and fibroblasts (right). (G) Heatmap of adipocyte-fibroblast interactions in all wounds. (H) Bar graph quantifying adipocyte cluster 1 - fibroblast cluster 2 interactions in unwounded, PBS, and P1i/P2i treated wounds. (I) Bar graph quantifying adipocyte cluster 2 - fibroblast cluster 2 interactions in unwounded, PBS, and P1i/P2i treated wounds. Scale bars, 50 um (D).

Differential interaction maps were then used to visualize the strength of spatial cell-cell interactions based on K-nearest-neighbor localization, which can be uniquely assessed using CODEX and other spatial phenotyping technologies. These spatial relationships were assessed in P1i-, P2i-, and PBS-treated wounds (**Fig. 8E and Fig. S21B**). PBS treated wounds showed strongly enriched interactions between adipocytes and fibroblasts compared to unwounded skin (**Fig. 8E, top left)**. In comparison, P1i-treated wounds and to a lesser extent, P2i-treated wounds, exhibited reduced adipocyte-fibroblast interactions compared to PBS-treated wounds (**Fig. 8E, top right and bottom)**.

To further assess the role of mechanosensitive gene expression in the transition from adipocyte to fibroblast phenotypes in wound healing, a protein expression module consisting of PIEZO1, PIEZO2, YAP, and PTK2 was computed for all CODEX annotated cell types. Two adipocyte clusters were identified by CODEX. Adipocyte cluster 2 had greater mechanosensitivity, as defined by the co-expression of these markers, compared to adipocyte cluster 1 (**Fig. 8F, left**). When analyzing PIEZO1 and PIEZO2 co-expression alone, adipocyte cluster 2 also showed greater expression than adipocyte cluster 1 (**Fig. S21C)**. The mechanosensitive signature of the six CODEX defined fibroblast subpopulations also varied, with clusters 1, 5, and 6 showing greater expression compared to clusters 2, 3, and 4 (**Fig. 8F, right**). Fibroblast clusters 1, 5, and 6 also displayed relatively higher co-expression of Perilipin and Adiponectin, suggesting these clusters to be ADFs (**Fig. S21D**). Furthermore, these putative ADFs were highly SCA1 positive, congruent with earlier findings (**Fig. S21E, Fig. 1E**).

Interestingly, the strength of cell-cell interactions between individual adipocyte and fibroblast subpopulations was also highly variable across the spatial microenvironment of wounds (**Fig. 8G and Fig. S21F**). The strongest adipocyte-fibroblast interactions were between fibroblast cluster 1 and adipocyte cluster 2, both of which are relatively mechanosensitive subpopulations (**Fig. 8F-G**). P1i and P2i treatment also modulated the strength of individual adipocyte-fibroblast subpopulation interactions compared to control, PBS-treated wounds (**Fig. 8H-I**). P2i treatment enhanced interactions between mechanically naïve subpopulations including adipocyte cluster 1 and fibroblast clusters 2 and 4, relative to PBS-treated wounds (**Fig. 8H and Fig. S21G**). In comparison, P1i-treated wounds shifted the interactions of mechanically sensitive adipocytes to co-localize more with mechanically naïve fibroblasts instead of mechanically sensitive fibroblasts, specifically enriching the interactions of adipocyte cluster 2 with fibroblast clusters 2 and 4 (**Fig. 8I and Fig. S21H**).

### Integration of RNA and protein spatial phenotyping modalities during wound healing

To understand if protein- and RNA-based spatial phenotyping revealed similar trends in cell spatial organization during wound repair, we compared cell-cell interactions using both techniques at POD 14 (**Fig. S22A**). Subclusters of cell types (e.g. Fibroblast cluster 1 and Fibroblast cluster 2) were combined into broad overlapping cell phenotypes to ensure that parallel cell populations and cell-cell networks could be compared using Visium and CODEX analysis. Interestingly, the cell-cell interactions in PBS versus P1i treated wounds at POD 14 were largely similar when evaluating between Visium and CODEX modalities (**Fig. S22B**). These data highlight the congruency in spatial analysis at an RNA and protein level during wound healing.

Collectively, these data suggest that P1i and P2i treatment differentially modify cell spatial organization in their mechanisms of supporting wound regeneration – in particular, by modulating interactions between mechanically defined adipocyte and fibroblast subpopulations.

### Spatial transcriptomics supports adipocyte to fibroblast transition in rescuing existing scars

To further analyze the spatial gene expression following scar rescue, the Visium platform was applied to scars treated with P1i or PBS at POD 30, 75, and 120 (**Fig. 9A**). Similar to analysis of acute wound healing, the epidermal, dermal, and hypodermal layers were easily elucidated from the spatial transcriptomic analysis in scars at POD 60 (**Fig. 9B**). Furthermore, the expected cell types including fibroblasts, immune, epithelial, and adipocytes were identified by spatial expression of representative genes (**Fig. S23A**).

**Figure 9:**
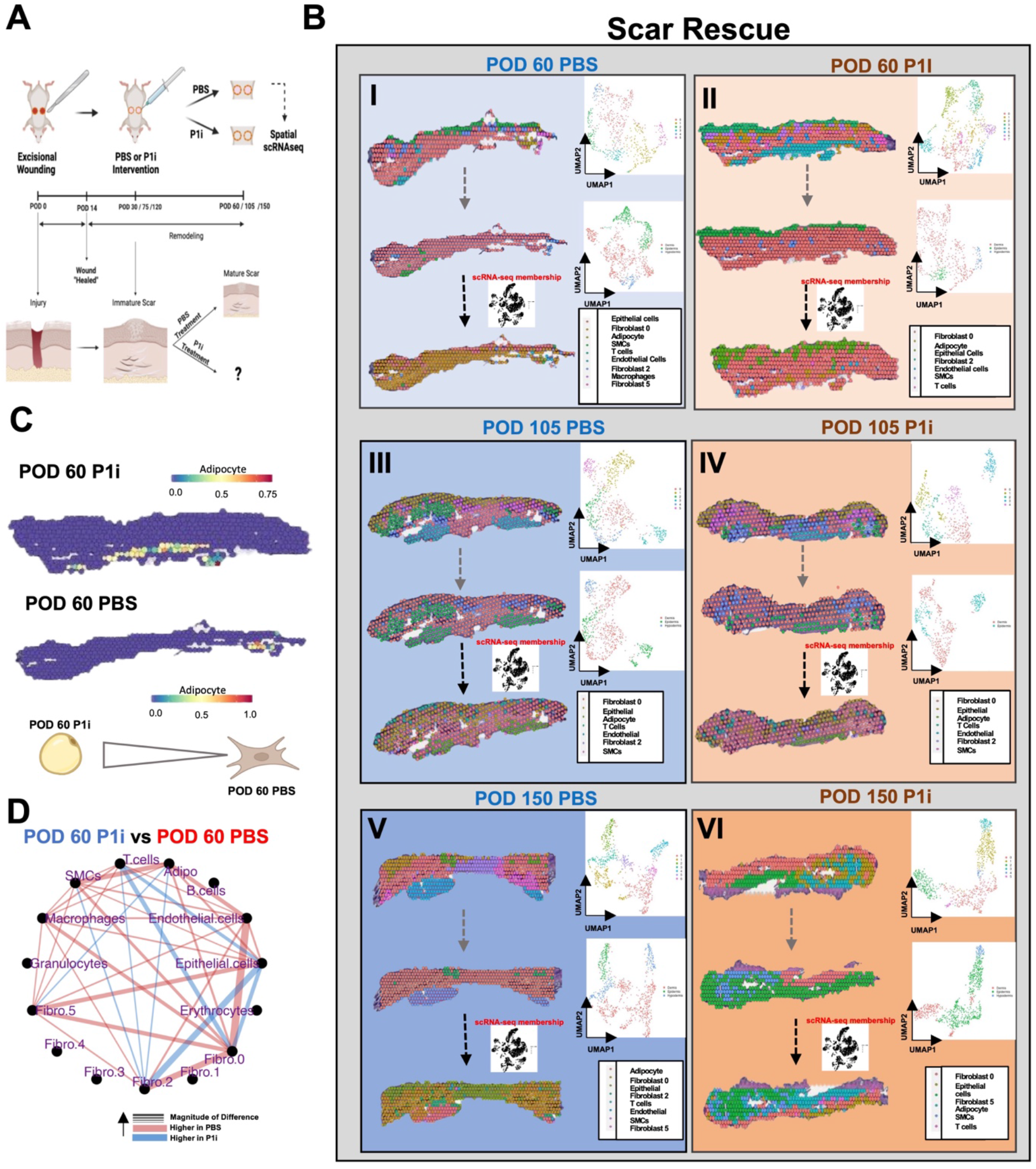
Visium analysis reveals that piezo inhibition during scar *rescue* alters the spatial transcriptional landscape. (**A**) Schematic for generating spatial transcriptomic data during scar rescue using the 10x Genomic protocol. (**B**) **[I-VI]** Top: Spatial plots (left) of scars colored by Seurat colors with UMAP (right). Middle: Spatial plots (left) of scars colored by tissue histology (left) with UMAP (right) showing that the three scar layers can easily be distinguished by their transcriptional programs. Bottom: Spatial plots (left) of scars colored by predicted cell types following anchor transfer from Fig.5 (left) with UMAP (right). Bottom: Schematic showing the anchor-based integration of scRNA-seq populations (defined in Fig.5) with Visium gene expression to project partial memberships within each spot across all groups. Arrows represent the same sample just colored by either UMAP or histological layer. (**C**) Spatial plot of adipocytes in P1i treated (top) and PBS treated (bottom) wounds at POD 60. (**D**) Differential interaction maps in P1i vs. PBS-treated wounds at POD 60.

Interestingly, the most differentially expressed genes during wound rescue following PBS or P1i treatment were similar to those defined during wound repair. P1i treated wounds showed high expression of genes associated with the epidermis, whereas PBS treated wounds revealed elevated expression of genes known to be associated with scarring (**Fig. S24**). To elucidate the spatial patterning of genes associated with the mechanically sensitive fibroblast populations, the spatial patterning of Piezo1, Piezo2, Yap1, and Ptk2 was evaluated across all timepoints (**Fig. S23B**). In P1i treated wounds the expression of Piezo1, Piezo2, Yap1, and Ptk2 were observed mostly at the basal aspects of the wounds compared to PBS treated wounds where they were found to be in the apical part of the wound.

To further clarify the role of ADFs in the spatial environment of established scars, the scRNA-seq defined cell populations were again anchor transferred onto the spatial transcriptomic analysis of histological sections (**Fig. 9B**). Over early (POD 60), mid remodeling (POD 105), and late remodeling (POD 150) the spatial proportions of adipocytes and fibroblasts varied with PBS and P1i treatment (**Fig. 9B**). The spatially identified adipocytes were greater in number in the P1i treated scars compared to PBS treated scars at POD 60, 105, and 150 (**Fig. 9C**). These data suggest that the adipocyte-fibroblast transition is not only transcriptionally evident at the single cell level but also distinct in the context of the spatial organization of the tissue.

To assess spatial communication of cell types in the scars, differential interaction maps were compared between the seven groups (**Fig. 9D**). Interestingly, P1i-treated wounds showed a larger number of interactions between adipocytes and epithelial cells, suggestive of regenerative communications (**Fig. 9D**). In comparison, PBS-treated wounds revealed a relatively higher number of adipocyte-fibroblasts communications, suggestive of possible fibrotic communications (**Fig. 9D**). Lastly, CytoTRACE further revealed, as observed during wound healing (**Fig. 7**), greater transcriptional diversity from the apical dermis compared to the basal dermis in a PBS-treated scar compared to a P1i-treated scar (**Fig. S25**).

Collectively, these data suggest that the adipocyte-fibroblast crosstalk, may play an important role in the spatial organization of pre-existing scars. Furthermore, mechanotransduction modulation may alter the spatial organization of the adipocyte to fibroblast transition in a way that facilitates the regeneration of established scars.

### Spatial phenotyping by CODEX demonstrates adipocyte-fibroblast cross-talk in rescuing existing scars

To phenotype the spatial environment in the context of rescuing existing scars, CODEX was further employed to spatially define protein expression, cell populations, and associated cell-cell interactions (**Fig. 10A**; **markers shown in Table 1 middle column, S26A**). We examined the spatial microenvironment of scars treated with P1i or PBS at POD 30 (early remodeling period) and harvested at POD 60, treated with P1i or PBS at POD 75 (mid- remodeling) and harvested at POD 105, and treated with P1i or PBS at POD 120 and harvested at POD 150 (late remodeling) (**Fig. 6**). Across all treatment groups, a manifold of 16 clusters was identified based on protein expression, including multiple subpopulations of adipocytes, fibroblasts, epithelial cells, smooth muscle cells, endothelial cells, and macrophages (**Fig. 10B-C**). PBS treated wounds showed higher expression of mechanical markers at POD 60, 105, and 150 compared to those treated with P1i (**Fig. 10D**). Differential interaction maps were then used to visualize spatially defined cell-cell interactions in P1i- and PBS-treated scars (**Fig. 10E**). Most notably, the interactions of adipocyte and fibroblast subpopulations varied between PBS and P1i treated scars at POD 30 (harvested at POD 60), 75 (harvested at POD 105), and 120 (harvested at POD 150) (**Fig 10E, S26B-D**). P1i treated scars showed enriched interactions mediated by adipocyte cluster 2 compared to PBS treated scars (**Fig. 10E**). On the other hand, PBS treated wounds showed enriched interactions mediated by adipocyte clusters 1 and 3 (**Fig. 10E**). Following P1i treatment, adipocyte cluster 2 interacted more strongly with epithelial cells, implicating regenerative communication, at POD 60, 105, and 150 (**Fig. 10F, left**). In comparison, following PBS treatment, adipocyte cluster 1 interacted more strongly with smooth muscle cells, suggestive of inflammatory and scar-associated communication (**Fig. 10F, right**).

**Figure 10:**
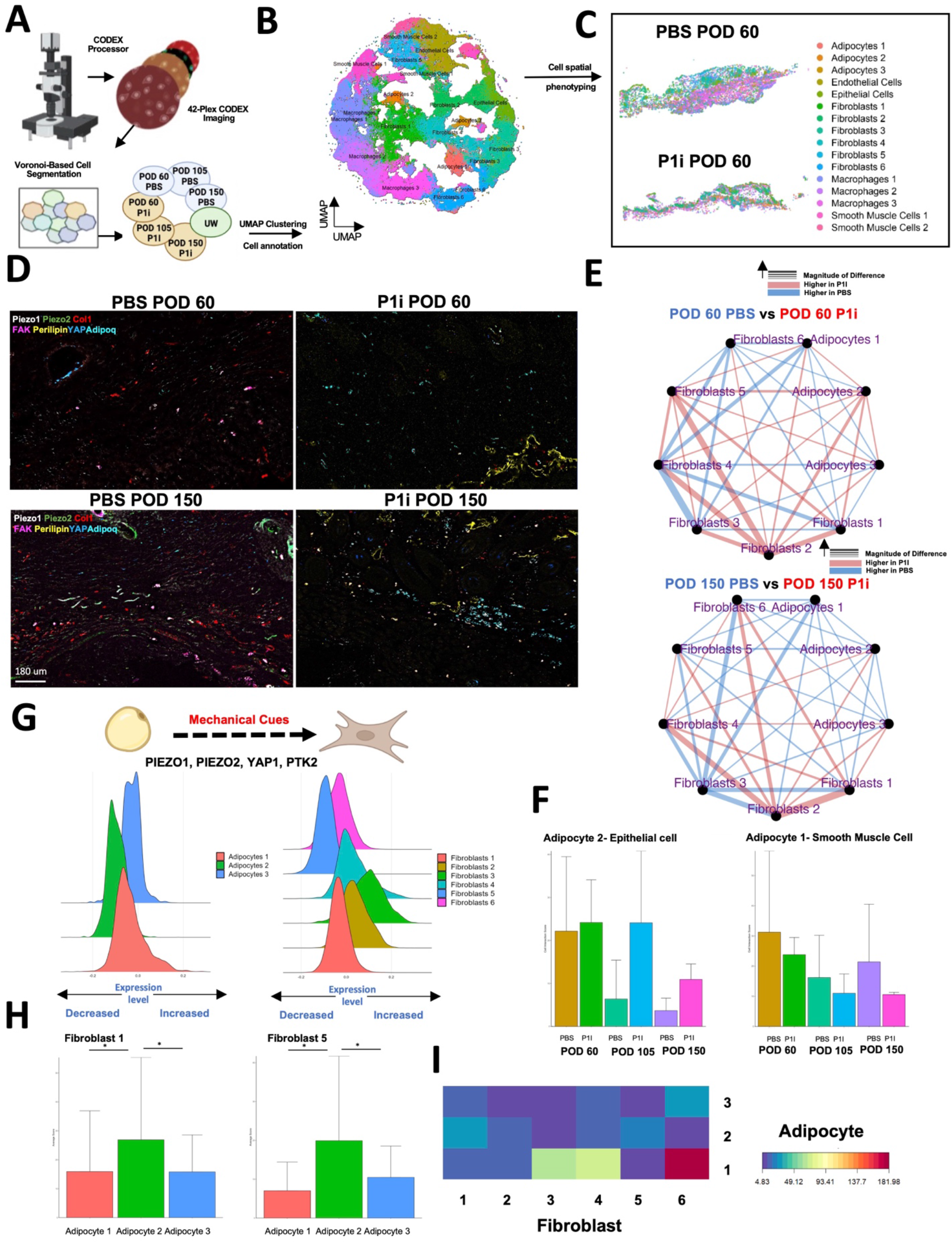
CODEX analysis reveals that piezo inhibition following scar rescue alters the spatial cross-talk between adipocytes and fibroblasts. (A) Schematic of the CODEX experiment. (B) UMAP plot of CODEX data for all sequenced cells. (C) Representative images of identified UMAP CODEX clusters showing their spatial distribution in PBS- and P1i-treated wounds. (D) Representative image of six selected CODEX markers in PBS (left)- and P1i- (right) treated wounds at POD 60 (top) and POD 150 (bottom). (E) Differential interaction maps in PBS- treated vs. P1i-treated wounds at POD 60 (top) and POD 150 (bottom). (F) Bar graph quantifying adipocyte cluster 2 - epithelial cell interactions (left) and adipocyte cluster 1 - smooth muscle cell interactions (right) in PBS and P1i treated wounds. (G) Histograms of mechanosensitive protein module expression (PIEZO 1, PIEZO2, YAP1, and PTK2) in adipocytes (left) and fibroblasts (right). (H) Bar graphs quantifying fibroblast 1 - adipocyte interactions (left) and fibroblast 5- adipocyte interactions (right) in all wounds. (I) Heatmap of adipocyte-fibroblast interactions in all wounds. Data shown as mean ± S.D. *\*P* ≤ 0.05, *\**\*\**P* ≤ 0.001. Scale bars, 180 um (D).

To further assess the role of mechanosensitive protein expression in adipocyte-fibroblast transition during scar rescue, a protein module of PIEZO1, PIEZO2, YAP1, and PTK2 was again analyzed in all CODEX annotated cell types (**Fig. S26E, left**). Adipocyte 1 and 3 had the greatest mechanical sensitivity, as defined by the co-expression of these markers compared to adipocyte cluster 2 (**Fig. 10G, left**). CODEX defined adipocyte cluster 3 also demonstrated the greatest collagen type I expression, indicative of a fibrotic phenotype (**Fig. S26F**). CODEX further revealed variability in the mechanical signature of the fibroblast clusters during scar rescue. Fibroblast clusters 2, 3, 4, and 6 showed greater mechanical sensitivity compared to clusters 1 and 5, indicative of a more fibrotic phenotype (**Fig. 10G, right**). Most interestingly, across the timepoints, the pro-regenerative mechanically naïve populations (Fibroblast 1 or 5 and Adipocyte 2) and pro-fibrotic mechanically sensitive populations (Fibroblast 2, 3, 4, 6 and Adipocyte 1 or 3) showed high levels of intra-group interactions (**Fig. 10H-I, S26G**). Collectively, CODEX spatial protein analysis suggested that signaling between adipocytes and fibroblasts markedly occurs during the rescue of existing scars. The extensive cross-talk observed between profibrotic, mechanically sensitive adipocytes and fibroblasts suggests that targeted mechanical modulation, which can be achieved by P1i, underlies wound regeneration during scar rescue.

### Piezo inhibition reduces scarring in a human foreskin xenograft wound model

Finally, while our results in mice were consistent with Piezo1 or Piezo2 blockade preventing adipocyte-to-fibroblast transition and reducing scarring, it is ultimately critical to determine whether similar anti-scarring effects are seen in human wounds. We developed a xenograft model to study healing by human cells, wherein human foreskin samples were engrafted onto CD-1 nude recipient mice then underwent full-thickness wounding (**Fig. 11A**). Unwounded grafted skin had similar histologic appearance, ECM ultrastructure, and Col1 expression compared to non-grafted skin (**Fig. S27A-C**), supporting that engraftment itself did not cause a fibrotic reaction within foreskin tissue. Following wounding, gross and histologic examination confirmed that distinct fibrotic scars formed within healed wounded xenografts (**Fig. 11B, D and S27D**). IF for human- and mouse-specific Col1 confirmed that xenografted human versus adjacent mouse skin were distinguishable on the basis of collagen and that the xenografts healed with human collagen (**Fig. 11G**, **top left panel, and S27D**), consistent with wound repair by human cells within xenografts (as opposed to wound repair by mouse cells from surrounding tissue). We next treated xenograft wounds with P1i or P2i. As in mice, Piezo inhibition grossly and histologically reduced scarring (**Fig. 11B-E**). ECM ultrastructure of P1i wounds was indistinguishable from that of unwounded xenograft skin (**Fig. 11F**) and Piezo inhibitor-treated scars had reduced collagen and increased adiponectin content (**Fig. 11G-H**). Piezo-inhibited wounds also had increased staining for CD31 (suggesting increased vascularization) and CK19 (consistent with regenerated secondary elements, e.g., Fordyce/oil glands; **Fig. 11I-J**). RNAscope *in situ* hybridization confirmed fewer Piezo1^+^ cells when Piezo1 was inhibited (**Fig. S27E**). Collectively, these results suggest that Piezo1 inhibition in human skin wounds yielded wound regeneration with minimized scarring.

**Figure 11:**
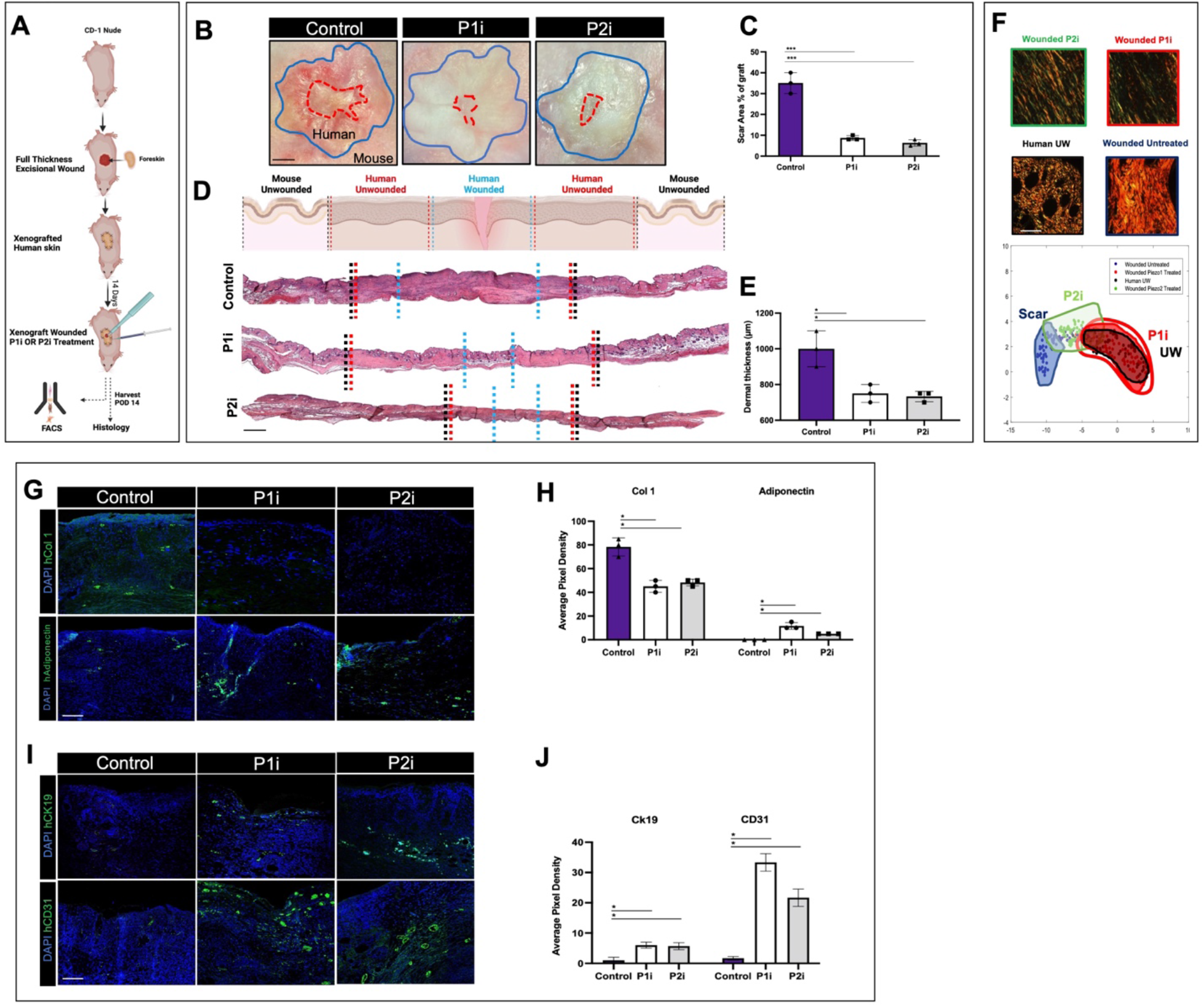
Piezo inhibition reduces scarring in a human skin xenograft wound model. (**A**) Schematic of human foreskin xenografting and wounding experiments. (**B**) Gross photos of healed postoperative day (POD) 14 xenograft wounds following treatment with Piezo1 inhibitor (P1i), Piezo2 inhibitor (P2i), or control (PBS). Blue outlined regions, xenograft; red dotted region, wound within xenograft. (**C**) Quantification of scar area as percentage of total graft area from gross images. (**D**) Schematic depiction (top row) and H&E histology (bottom three rows) xenograft wounds. Dotted lines on histology images indicate boundaries of regions identified in schematic (e.g., mouse unwounded [UW], human UW) in corresponding color font. (**E**) Quantification of xenograft wound dermal thickness from histology. (**F**) Top, picrosirius red staining of xenograft wounds. Bottom, UMAP of quantified extracellular matrix (ECM) ultrastructure parameters for indicated xenograft conditions. (**G**) IF staining for human-specific Col 1 (hCol1, top) or adiponectin (bottom; both, green signal) in indicated xenograft wound conditions. (**H**) Quantification of hCol1 and adiponectin expression in xenograft wounds from IF. (**I**) IF staining for human CK19 (hCK19, top) or human CD31 (hCD31, bottom; both, green signal) in indicated xenograft wound conditions. (**J**) Quantification of hCK19 and hCD31 expression in xenograft wounds from IF. Data shown as mean ± S.D. *\*P* ≤ 0.05, *\**\*\**P* ≤ 0.001. Scale bars, 500 µm (**B, D**), 25 µm (**F**), 50 µm (**G, I**).

### Mechanically naïve human fibroblasts are identified in P1i treated xenografts using scRNA-seq

To assess if P1i acts on similar fibroblast clusters to cause regeneration, scRNA-seq analysis was performed on xenografts harvested at POD 14 post injury using a hybrid human-mouse xenotranscriptome **(Fig. 12A**). Mouse cells were removed informatically and subsequent analysis focused exclusively on human cells (**Fig. S28A**). Partitional analysis identified 8 transcriptionally-distinct clusters **(Fig. 12B-C**), corresponding to fibroblast and keratinocyte populations (**Fig. S28A**). Each cluster was present in all three treatment conditions but Cluster 3 was elevated in P1i treated wounds (**Fig. 12D, Fig S28B**). Interestingly, cluster 3 was characterized by elevated expression of pro-regenerative genes such as TWIST2, WNT5A, CALU, and SFRP2 (**Fig**. **12E-F**) which were found to be highly expressed in P1i-treated wounds (**Fig**. **12F and Fig. S29A**). This cluster was also enriched for pathways such as “skin morphogenesis”, as well as “neural crest differentiation” (**Fig. 12G**). By contrast, Cluster 0 was enriched for pathways related to adipocyte differentiation (**Fig. S28C**) and elevated expression of genes related to adipocyte-fibroblast transition (**Fig. S28C**) including expression of JUN a transcription factor associated with fibrosis (**Fig. S28D**).

**Figure 12:**
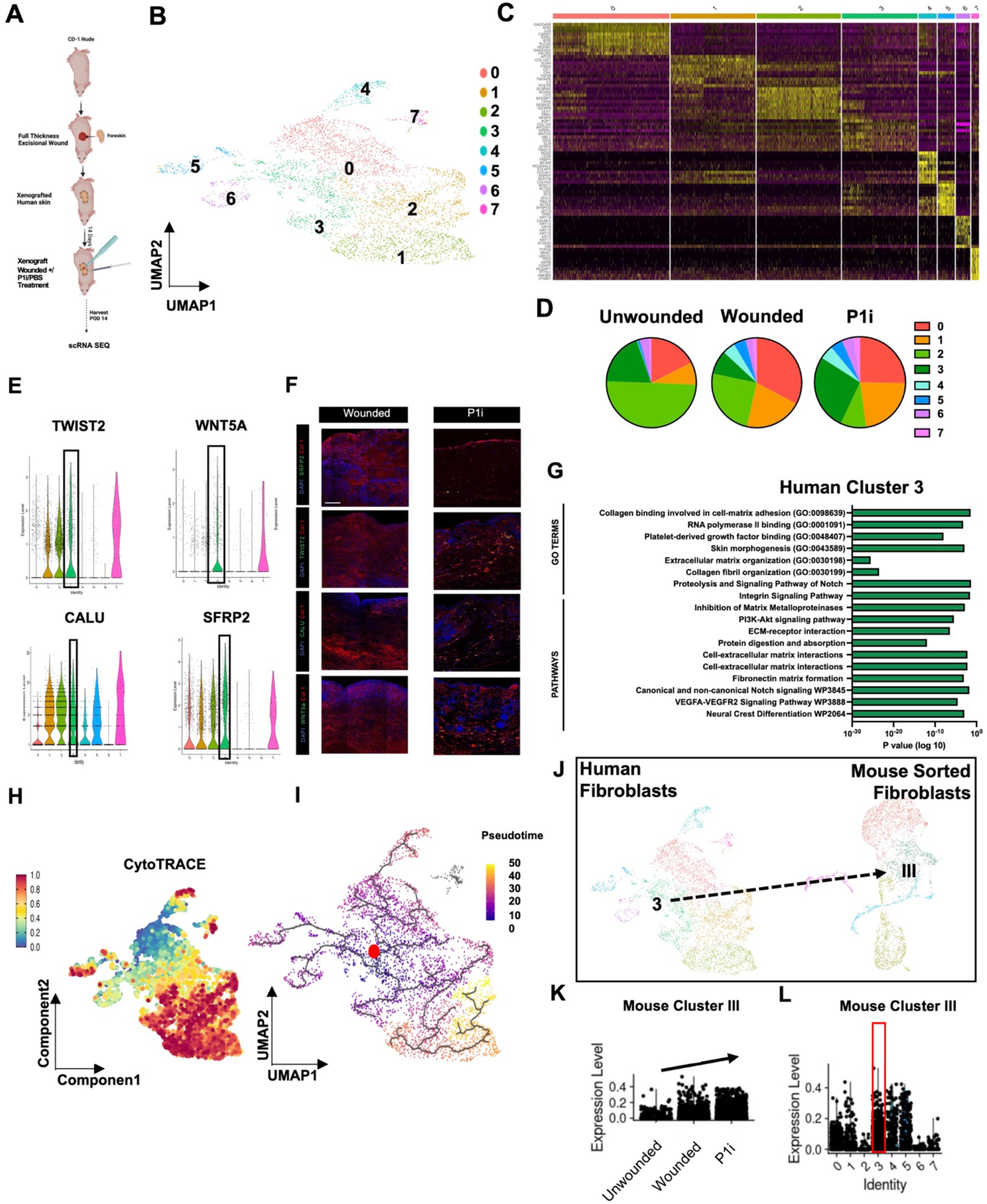
Piezo inhibition reduces scarring in a human skin xenograft wound model through altering the transcriptional profile of resident cells. (A) Schematic of the xenograft wounding model. (B) UMAP of scRNA-seq data from all wound cells, colored by Seurat cluster. (C) Heatmap showing top differentially expressed genes for each Seurat fibroblast cluster. (D) Relative representation of fibroblasts belonging to each Seurat cluster from each experimental condition. (E) Violin plots showing gene expression of Twist2, Wnt5a, Calu, and Sfrp2 among the Seurat clusters. (F) Immunostaining of Sfrp2, Twist2, Calu, and Wnt5a (green) with co expression of collagen type I (red) at Post-operative day 14 (POD 14) in wounded (left) and P1i treated wounds (right). (G) Gene Ontology (GO) pathway analysis for human cluster 3. (H) CytoTRACE analysis of the fibroblast clusters. (I) Pseudotime analysis of the fibroblast clusters. (Red dot shows the root point) (J) Schematic of the anchor transfer of human fibroblasts onto the mouse sorted fibroblasts (from Fig.2) with arrows demonstrating the clusters that were transcriptionally similar. (K) Bar graph depicting the similarity of the mouse cluster 3 onto the human xenograft unwounded, wounded and P1i treated derived fibroblasts. (L) Bar graph depicting the similarity of the mouse cluster 3 onto the human xenograft fibroblasts derived Seurat clusters. Data shown as mean ± S.D. *\*P* ≤ 0.05, *\**\*\**P* ≤ 0.001. Scale bars, 250 µm (F).

CytoTRACE analysis identified significant differences in the differentiation state of cluster 3 cells compared to other fibroblast clusters (**Fig. 12H**), allowing us to construct pseudotime trajectories from this cluster (**Fig. 12I**). This demonstrated significant changes in the expression of key genes such as THY1, ROBO2, TRPS1, and TWIST2, throughout the putative transition of cells from cluster 3 to other subgroups (**Fig. S28E**). Pseudotime analysis suggested a trajectory from cluster 3, to cluster 0 (associated with JUN expression), and then to cluster 1 (associated with VCAM1 expression) (**Fig. 12I and Fig. S28C-D**). These candidate trajectories were also supported by RNA velocity analysis (**Fig. S28F**). Furthermore, mechanosensitive genes (PIEZO1, PIEZO2, YAP1, and PTK2) were highly expressed over these pseudotime trajectories (**Fig S28G**).

CellChat also demonstrated a downregulation of cluster 0 activity with other clusters following P1i compared to control wounds (**Fig. S28H-I**).

Interestingly, to evaluate these clusters in the context of our previously-defined fibroblast populations, we performed cross-species mapping using a label transfer-based approach (**Fig. 12J- L**), which suggested that cluster 3 cells were of highest similarity to our mechanically-naive mouse fibroblast III cells. These data suggest that mechanically naïve fibroblast populations may exist during human wound healing.

### Mechanical cues may convert human regenerative ADFs towards fibrotic ADFs

To further confirm that human cluster 3 represented a pro-regenerative ADF population we utilized our *in vitro* collagen gel system. We first, FACS sorted human fibroblasts based on expression of cluster 3 markers including ROBO2, CD29, and SFRP2 and seeded them into 3D collagen gels (**Fig**. **S29B)**. Upon stretching for 48 hours, these fibroblasts demonstrated high expression of typical fibroblast markers including Collagen Type I and low expression of typical adipocyte markers including PPAR*γ* using IF analysis (**Fig. S29B**). In contrast, stretching with P1 inhibition kept the fibroblasts in a more adipocyte-like phenotype, with high expression of PPAR*γ* and low expression of Collagen Type I, similar to fibroblasts in an unstretched state (**Fig. S29B**). These data suggest that mechanotransduction may induce the phenotype of the regenerative human cluster 3 towards a more fibrotic phenotype.

The scRNA analysis also demonstrated high expression of Wnt5A and Calu in human fibroblast cluster 3, both of which are known to play a role in the Wnt signaling pathway. To determine if P1i may activate canonical Wnt pathways to maintain cluster 3 in an adipocyte-like phenotype, we applied Wnt5a recombinant protein to the *in vitro* collagen gel system (**Fig. S29C**).

As expected upon stretching of human fibroblasts displayed low expression of regenerative cluster 3 marker, CD29 but high expression of Cluster 0 (CJUN) and 1 (VCAM1) markers (**Fig. S29C**). In contrast, in the presence of either P1i or Wnt5a recombinant protein human fibroblasts demonstrate a more adipocyte like phenotype with high expression of PPAR*γ* and low expression of Collagen Type 1 (**Fig. S29C**). In summary, P1i treatment may revert fibroblasts towards the mechanically naïve fibroblast pro-regenerative fibroblast population through the activation of canonical Wnt pathways and thereby preventing the transition to pro-scarring phenotypes defined by CJUN and VCAM1 expression.

### Spatial proteomic phenotyping by CODEX reveals adipocyte plasticity in wound repair

While scRNA-seq of human xenografts investigates human fibroblast subpopulations at a transcriptomic level, we further utilized CODEX to analyze spatially defined protein expression and cellular organization (**Fig. 13A; markers shown in Table 1 right column**). Specifically, we compared the spatial microenvironment of PBS, P1i, and unwounded skin at POD 14. Across all treatment groups, a manifold of 20 clusters was identified based on protein expression, including multiple subpopulations of adipocytes, fibroblasts, epithelial cells, smooth muscle cells, helper T cells, cytotoxic T cells, endothelial cells, B cells, and macrophages (**Fig. 13B, S30A**). Broadly, the spatial pattern of the protein markers was similar between P1i treated wounds and unwounded skin (**Fig. 13B-C**). Interestingly, CODEX analysis identified four adipocyte subpopulations based on protein expression (**Fig. 13B**). We thus compared the representation of adipocyte subtypes within P1i- versus PBS-treated wounds (**Fig. 13E, S30B**). The relative proportions of CODEX-defined adipocyte clusters significantly differed based on wound treatment, with fewer of the adipocyte 1 subtype and more of the adipocyte 4 subtype in P1i-treated wounds compared to PBS-treated wounds (\**p* < 0.05; **Fig. 13E, S30B**).

**Figure 13:**
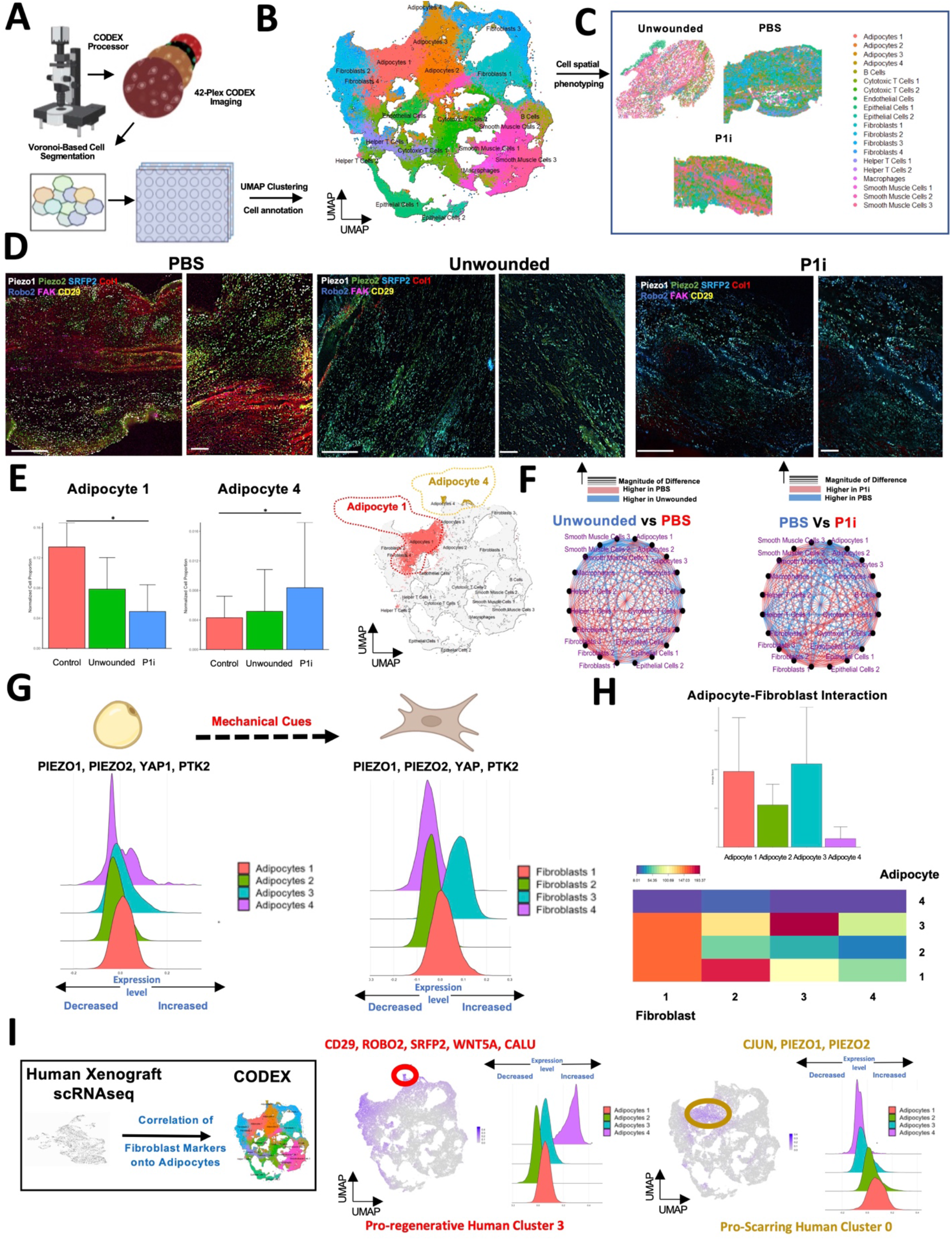
CODEX analysis reveals that piezo inhibition in human xenografts alters spatial cell cross-talk. (**A**) Schematic of the CODEX experiment. (**B**) UMAP plot of CODEX data for all sequenced cells. (**C**) Representative images of identified UMAP CODEX clusters showing their spatial distribution in unwounded skin and PBS- and P1i**-**treated wounds. (**D**) Representative images of six selected CODEX markers in unwounded skin and PBS- and P1i **-**treated wounds. (**E**) Bar graphs quantifying adipocyte cluster 1 (left) and adipocyte cluster 4 (middle) proportions in unwounded skin and PBS- and P1i-treated wounds, with representative feature plot of adipocyte cluster 1 and cluster 4 (right). (**F**) Differential interaction maps in unwounded skin vs. PBS-treated wounds (left) and PBS-treated wounds vs. P1i-treated wounds (right). (**G**) Histograms of mechanosensitive protein module expression (PIEZO 1, PIEZO2, YAP, and FAK) in adipocytes (left) and fibroblasts (right). (**H**) Bar graph of adipocyte subpopulation interactions with all fibroblasts (top) and heatmap of individual adipocyte-fibroblast interactions (bottom) in all wounds. (**I**) Correlation between fibroblast clusters identified by scRNA-seq and CODEX analyses (left). Feature plots showing that scRNA-seq cluster 3 fibroblasts mapped onto CODEX Type 4 adipocytes (middle) and scRNA-seq cluster 0 fibroblasts mapped onto CODEX Type 1 adipocytes (right). Data shown as mean ± S.D. *\*P* ≤ 0.05, *\**\*\**P* ≤ 0.001. Scale bars, 150 um left 50 um right

Differential interaction maps were then used to visualize spatially defined cell-cell interactions in P1i- and PBS-treated wounds (**Fig. 13F**), particularly for the adipocyte 1 and 4 subpopulations that were differentially represented across treatments. PBS-treated wounds showed broadly enriched interactions between fibroblasts and adipocytes compared to P1i-treated wounds (**Fig. 13F and Fig. S30C-E**). Interactions by individual adipocyte clusters also varied among the three treatment groups. PBS-treated wounds showed highly enriched interactions between adipocytes 1 and various immune cells (e.g. cytotoxic T cells, helper T cells, B cells), suggestive of inflammatory and scar-associated communication (**Fig. S30F**). In contrast, P1i-treated wounds showed strong crosstalk between adipocytes 4 and epithelial cells, suggestive of regenerative interactions (**Fig. S30F**).

To further assess the role of mechanosensitive expression in the transition from adipocyte to fibroblast phenotypes, a protein module of PIEZO1, PIEZO2, YAP, and PTK2 was analyzed in all CODEX annotated cell types (**Fig. S31A**). Adipocyte and fibroblast subpopulations had varying levels of mechanical sensitivity, as defined by the co-expression of these markers (**Fig. 13G)**. Interestingly, adipocyte clusters 1 and 3 exhibited strong protein expression of PIEZO1, PIEZO2, YAP, and PTK2 suggesting that these clusters have a more mechanically activated phenotype compared to adipocyte clusters 2 and 4 (**Fig. 13G**). On the other hand, fibroblast clusters 1 and 3 had greater mechanosensitive expression compared to fibroblast clusters 2 and 4 (**Fig. 13G**).

We also analyzed a canonical fibroblast-like (COL1, COL4) protein expression module in each of the adipocyte subpopulations to infer relative transitional status. Adipocyte cluster 4 had the lowest expression of fibroblast-like markers (COL1, COL4) compared to adipocyte clusters 1, 2, and 3, indicating that adipocyte cluster 4 may represent a “mechanically naïve”, less fibroblast-like state compared to adipocytes 1, 2, and 3 (**Fig. S31B**). Interestingly, adipocyte cluster 4 demonstrated a bimodal expression profile for both mechanical sensitivity (PIEZO1, PIEZO2, YAP1, and PTK2; **Fig. 13G**) and fibroblast-like proteins (COL1, COL4; **Fig. S31B**), in addition to high Ki67 expression (**Fig. S31C**), suggesting that a subset of this cluster may be transitioning towards a more fibroblast-like phenotype and further underscoring the observed plasticity of adipocytes. Adipocyte cluster 1 and 3 also interacted more extensively with fibroblasts in the spatial microenvironment compared to adipocytes 2 and 4 (*p < 0.05) (**Fig. 13H, top panel**). Taken together, these data suggest that mechanically sensitive adipocytes have higher expression of fibroblast-like markers and enriched spatial interactions with fibroblasts.

We similarly analyzed a canonical adipocyte-like (ADIPOQ, PERILIPIN) protein expression module in each of the fibroblast subpopulations to analyze relative transitional status. Fibroblast clusters 1 and 3 had lower expression of adipocyte-like markers compared to fibroblast clusters 2 and 4 (**Fig. S31D**), as well as relatively higher mechanosensitive protein expression (PIEZO1, PIEZO2, YAP1, and PTK2; **Fig. 13G**). Fibroblasts cluster 1, the subtype with the greatest mechanosensitive expression, had the greatest degree of interaction with adipocyte clusters 1, 2, and 3 but not adipocyte cluster 4 (**Fig. 13H, bottom panel**). Thus, mechanically sensitive fibroblasts demonstrated lower expression of adipocyte-like markers, but a higher degree of spatial interaction with adipocytes. Overall, our analyses of CODEX-defined fibroblasts and adipocytes established a strong association between mechanically activated protein expression, enhanced adipocyte-fibroblast spatial interactions, and further transition from adipocytes to fibroblasts.

Lastly, we mapped the pro-scarring (CJUN/PIEZO1/PIEZO2+) and pro-regenerative (ROBO2^/^SFRP2/WNT5A/CALU+) ADF clusters from our human xenograft scRNA-seq analysis to the CODEX dataset (**Fig. 13I left and S31A middle and right**) . Specifically, CODEX-defined adipocytes (PERILIPIN^+^ADIPOQ^+^) were interrogated using expression modules of the aforementioned markers for pro-regenerative scRNA-seq cluster 3 and pro-scarring scRNA-seq cluster 0 (**Fig. 13I left and S31A middle and right**). Interestingly, CODEX adipocyte cluster 4, which demonstrated strong epithelial cell spatial interactions, mapped closely to the putatively pro-regenerative scRNA-seq fibroblast cluster 3 (**Fig. 13I middle**). CODEX adipocyte cluster 1, which demonstrated strong immune-adjacent and inflammatory interactions, mapped closely to the putatively pro-scarring scRNA-seq fibroblast cluster 0 (**Fig. 13I right**). This further suggests that human ADF scRNA-seq clusters 3 and 0, which display regenerative and fibrotic expression, respectively, may be derived from adipocytes with similar protein expression profiles. In summary, CODEX spatial protein analysis suggested that adipocytes display heterogeneity in their expression of mechanosensitive proteins, which produces distinct effects on their spatial interaction niche and their expression of markers along the adipocyte-to-fibroblast transition. Furthermore, fibrotic and regenerative markers identified by scRNA-seq strongly correlate with distinct protein-defined adipocyte subtypes. Collectively, these results suggest that mechanical cues are critical to facilitating the adipocyte to fibroblast transition in human wound repair and fibrosis.

## Discussion

Overall, we have shown via adipocyte engraftment and genetic lineage-tracing studies that mature adipocytes lose adipogenic markers and transition to fibroblasts in wounds. Using both *in vitro* and *in vivo* models to modulate adipocyte mechanical environment, as well as single-cell transcriptomic analysis in Rainbow mice, we demonstrated that transition of both mouse and human adipocytes into adipocyte-derived fibroblasts is a mechanically-driven process, differentiating via a “mechanically naïve” fibroblast intermediate and involving signaling by mechanosensitive ion channel (Piezo) genes (**Fig. S32**). These results elucidate a molecular signature and role of adipocytes and adipocyte-fibroblast dynamics in contributing to dermal fibrosis during wound healing, and highlight a novel mechanism through which wound mechanics drive fibrosis and govern adipocyte fate. Finally, we found that blocking adipocyte mechanotransduction via Piezo1 inhibition prevents adipocyte-to-fibroblast transition and yields significantly reduced scarring with features of wound regeneration, a result which suggests that adipocyte-fibroblast differentiation is a meaningful contributor to scarring and can be targeted to prevent fibrosis.

Our investigations into the effects of Piezo inhibition during the remodeling phase of healing has particularly exciting translational implications, as they suggest that Piezo inhibition could even rescue existing scars, which would have important therapeutic implications for the hundreds of millions of patients who suffer from existing scars. We have demonstrated the importance of the adipocyte to fibroblast transition in skin fibrosis due to mechanical forces at both a gene and protein level using novel Visium and CODEX integration. Furthermore, by combining scRNA-seq with Visium and CODEX, we provide extensive support for the relevance of adipocyte mechanosensing in both wound repair and established scars.

While results in our novel foreskin xenograft model are promising for ability to translate these findings to wound healing mediated by human cells, a key limitation is that foreskin inherently differs from skin at other body sites; for instance, it lacks HF and, importantly, has relatively minimal associated fat. However, this model allowed us to examine the global effect of P1i/P2i on healing by human fibroblasts; further studies of other types of human wounds and the cellular impact of P1i and P2i in human cell-mediated wound healing will be necessary to determine the mechanism(s) of P1i/P2i-induced regeneration in human wounds.

While the roles of canonical mechanotransduction (e.g., FAK, YAP) in driving pro-fibrotic fibroblast activity have been broadly studied, the present findings suggest an entirely new pathway and mechanism by which *adipocyte* mechanosensing can influence fibrosis. Although the data provided are encouraging, further work is need to fully characterize the adipocyte-fibroblast transition observed following murine skin injury. Firstly, to complement the scRNAseq analysis of murine skin wounds over time, single nuclei RNA-sequencing (sNuc-Seq) will be performed to identify and characterize the adipocyte subpopulation/s following skin injury and those that contribute to the adipocyte-fibroblast transition. Secondly, the Visium and CODEX spatial analysis in acute wounds and established scars will be further analyzed to understand how the location differs between the mechanical naïve and mechanical sensitive fibroblast subpopulations in the wound bed and how this may influence the transition of one cell state to another to reduce skin scarring. Lastly, further lineage tracing experiments will be performed to understand how Piezo inhibition rescues murine skin scars including how inhibition from adipocytes to fibroblasts could rescue skin scars. Further analysis using immunostaining of the hair follicle formation following Piezo inhibition in established scars will be performed. These data will provide further understanding into the role of the adipocyte-fibroblast transition in skin scarring.

We have found that Piezo proteins are highly expressed on adipocytes and not fibroblasts; as contributions of adipocytes to scarring remain relatively unknown, and contributions of mechanosignaling to adipocyte biology even more so, these findings could open a new path of inquiry through which a distinct cell lineage could be a functionally important and targetable driver of fibrosis. Our findings also raise the interesting possibility that distinct mechanical signaling pathways may be involved in driving fibrosis from the perspective of different cell types (e.g., adipocytes versus fibroblasts, or even different fibroblast subsets). Future work will be of interest in more robustly elucidating which of these pathways are most relevant to therapeutically targeting different human fibroses.

## Acknowledgments

We thank the Stanford Functional Genomics Facility, Stanford Cell Sciences Imaging Facility, and Stanford Shared FACS Facility Cores.

## Author contributions

Conceptualization: MFG, HET, NJG

Formal analysis: MFG, HET, NJG, AFS, SM, DH, NQ, MJ Funding acquisition: HPL, GCG, DCW, MTL

Investigation: MFG, HET, NJG, AFS, KC, JLG, SM, MK, KEBR, DBA, NMDD, DS, EJF, JBLP, ACC, JC, MD, DA, CB, NL

Methodology: MFG, HET, NJG, AFS, KC, JLG, SM, KEBR Software: MFG, NJG, SM, DH, MJ

Supervision: NQ, OK, HPL, GCG, DCW, MTL Visualization: MFG, HET, NJG, AFS, DH, MJ Writing – original draft: HET, MFG, MJ

Writing – review & editing: NQ, OK, HPL, GCG, MJ, DCW, MTL

## Data and materials availability

The scRNA-seq data generated during this study will be deposited in a GEO repository and made publicly available prior to publication. Original scripts for the ECM ultrastructure algorithm used in this study have been deposited in a Github repository (https://github.com/shamikmascharak/Mascharak-et-al-ENF) and are publicly available. Further information and requests for resources and reagents should be directed to and will be fulfilled by the lead contact, Michael T. Longaker.

