## Supplementary Figures for "Piezo inhibition prevents *and* rescues scarring by targeting the adipocyte to fibroblast transition"

### Legends and Methods

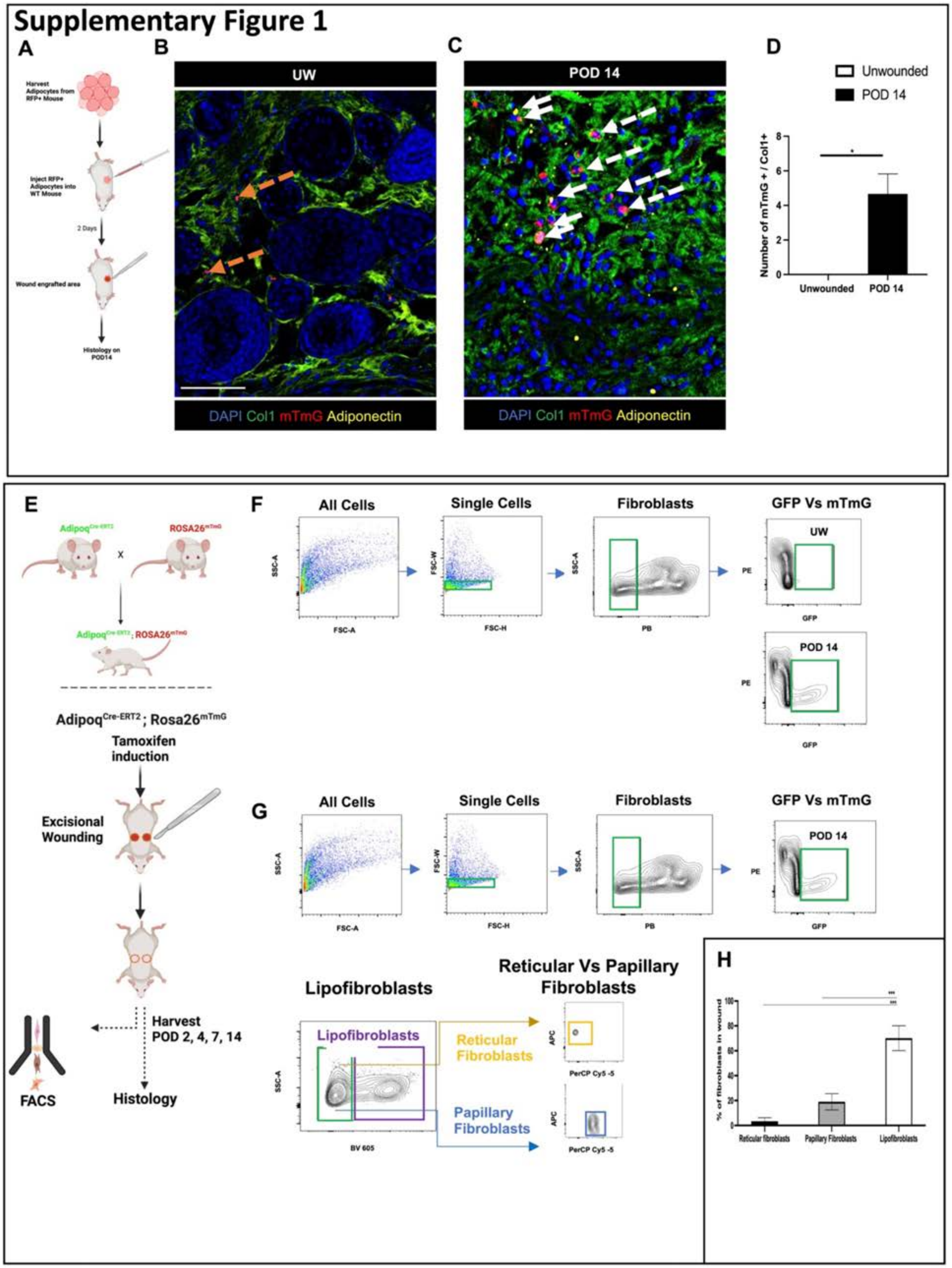

Supplementary Figure 1
A
B
C
D
UW
POD 14
DAPI Col1 mTmG Adiponectin
DAPI Col1 mTmG Adiponectin
Unwounded
POD 14
Number of mTmG + / Col1+
Unwounded
POD 14
E
Adipoq
ROSA26
Adipoq[Cre-ERT2]; Rosa26[mTmG]
Tamoxifen induction
Excisional Wounding
Harvest POD 2, 4, 7, 14
FACS
Histology
F
All Cells
Single Cells
Fibroblasts
GFP Vs mTmG
SSC-A
FSC-A
FSC-W
FSC-H
PB
PE
GFP
UW
POD 14
G
All Cells
Single Cells
Fibroblasts
GFP Vs mTmG
POD 14
Lipofibroblasts
Reticular Vs Papillary Fibroblasts
Lipofibroblasts
Reticular Fibroblasts
Papillary Fibroblasts
BV 605
APC
PerCP Cy5 -5
H
% of fibroblasts in wound
Reticular fibroblasts
Papillary Fibroblasts
Lipofibroblasts

**Figure S1: Adipocyte transplantation and wounding experiments. A.** Schematic of fluorescent (*mTmG*, Tomato red fluorescent protein [RFP]$^+$) adipocyte transplantation and wounding experiments. **B-C.** Immunofluorescent (IF) histology of transplanted *mTmG* adipocytes in unwounded skin (**B**) and postoperative day (POD 14) wounds (**C**) with IF staining for fibroblast and adipocyte markers. mTmG, Tomato fluorescence from transplanted adipocytes (red fluorescent signal); IF staining for adiponectin (yellow signal) and Col1 (green signal). Dotted orange arrows, Tomato$^+$ cells; solid white arrows, Adipoq$^+$Col1$^+$Tomato$^+$ cells; dotted white arrows, Adipoq$^-$Col1$^+$Tomato$^+$ cells. Scale bar, 50 µm. **D.** Quantification of transplanted *mTmG* cells also expressing Col1 per high-powered field (HPF). **E.** Top: Breeding scheme of $Adipoq^{Cre\text{-}ERT};R26^{mTmG}$ mice. Bottom: Schematic of $Adipoq^{Cre\text{-}ERT};R26^{mTmG}$ wounding experiments for lineage tracing of adipocyte-derived cells. **F.** Fluorescence-activated cell sorting (FACS) strategy for sorting GFP$^+$ (adipocyte-derived; GFP$^+$ gate) versus Tomato$^+$ (mTmG, non-adipocyte-derived; PE$^+$ gate) fibroblasts (Lin$^-$; PB$^-$ gate) with representative plots for unwounded (UW) and POD 14 wound sorts. **G.** FACS strategy for sorting lipo- (Sca1$^+$; BV 605$^+$ gate), reticular (Dlk1$^+$; APC$^+$ gate), and papillary dermal (CD26$^+$; PerCP Cy5.5$^+$ gate) ADF (Lin$^-$GFP$^+$) subtypes. **H.** Quantified percentage of wound ADFs expressing markers of each fibroblast subtype by FACS.

**D, H.** Data shown as mean ± S.D. *$P \leq 0.05$, ***$P \leq 0.001$.

### Supplementary Figure 2

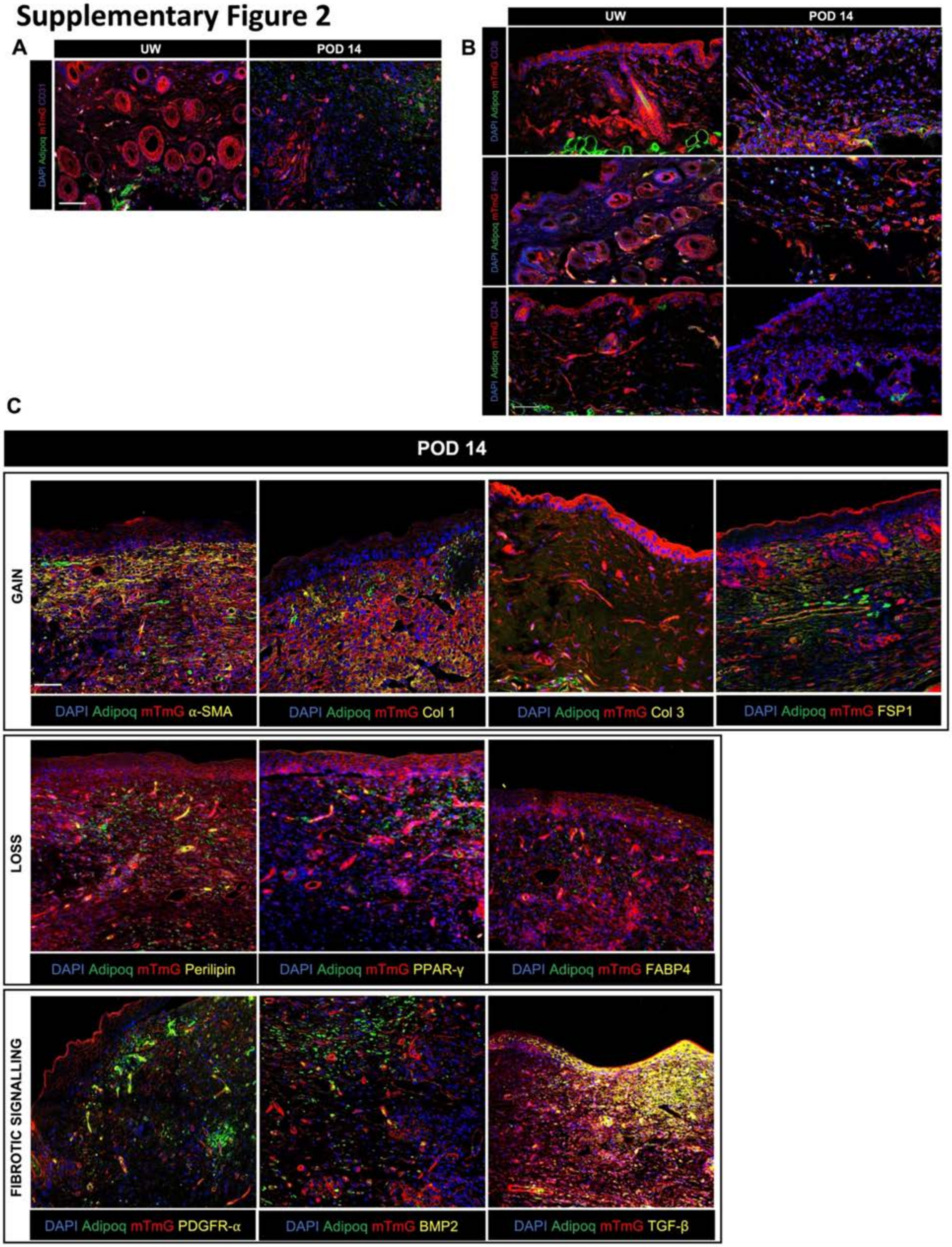

**Figure S2**: **Immunofluorescent analysis of ADF marker expression in *Adipoq*$^{Cre\text{-}ERT}$*;R26*$^{mTmG}$ skin and wounds.** **A.** Immunofluorescent (IF) staining of unwounded (UW) skin and postoperative day (POD) 14 wounds for CD31 (endothelial cell marker; purple fluorescent signal). **B.** IF staining of UW skin and POD 14 wounds for CD8 and CD4 (T cell markers) and F480 (macrophage marker; all, purple signal). **C.** IF staining of POD 14 wounds showing that ADFs gain expression of indicated fibroblast markers (top row), lose adipocyte markers (second row), and localize with known fibrotic signaling markers (bottom row; all IF, yellow signal). **A-C.** DAPI, nuclear counterstain (blue signal); mTmG, Tomato$^{+}$ (*Adipoq* lineage-negative) cells (red signal); Adiponectin, GFP$^{+}$ (*Adipoq* lineage-positive) cells (green signal). Scale bars, 25 µm.

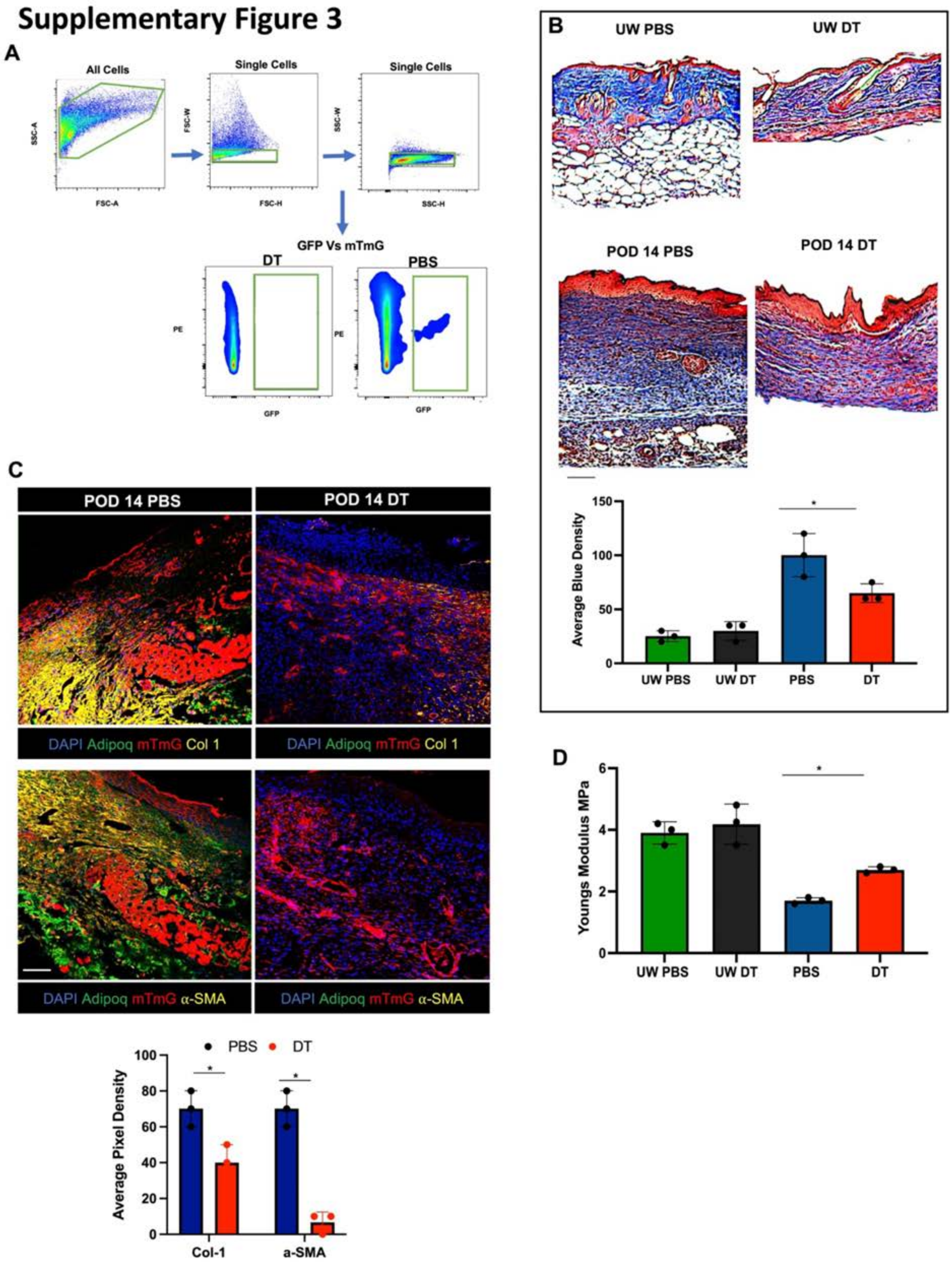

Supplementary Figure 3
A
All Cells
SSC-A
FSC-A
Single Cells
FSC-W
FSC-H
Single Cells
SSC-W
SSC-H
GFP Vs mTmG
DT
PBS
PE
GFP
B
UW PBS
UW DT
POD 14 PBS
POD 14 DT
Average Blue Density
UW PBS
UW DT
PBS
DT
C
POD 14 PBS
POD 14 DT
DAPI Adipoq mTmG Col 1
DAPI Adipoq mTmG α-SMA
PBS
DT
Average Pixel Density
Col-1
a-SMA
D
Youngs Modulus MPa
UW PBS
UW DT
PBS
DT

**Figure S3: Analysis of adipocyte ablation in *Adipoq*$^{Cre\text{-}ERT}$*;R26*$^{mTmG}$*;R26*$^{Awai}$ wounds. A.** Fluorescence-activated cell sorting (FACS) strategy for quantifying GFP$^{+}$ (adipocyte-derived) wound cells with representative plots for diphtheria toxin (DT)- and PBS-treated wounds showing that GFP$^{+}$ cells are ablated with DT treatment. **B.** Masson's trichrome staining (top) and quantification of average blue (connective tissue) staining density (bottom) for PBS- and DT-treated unwounded (UW) skin and POD 14 wounds. **C.** Top, fluorescent histology of POD 14 PBS- and DT-treated wounds with IF staining for Col 1 or α-smooth muscle actin (α-SMA) (yellow). DAPI, nuclear counterstain (blue signal); mTmG, Tomato$^{+}$ (*Adipoq* lineage-negative) cells (red signal); Adipoq, GFP$^{+}$ (*Adipoq* lineage-positive) cells (green signal). Bottom, quantification of Col 1 and α-SMA expression from IF. **D.** Young's modulus calculated from tensile strength testing of PBS- and DT-treated wounds or UW skin.

**B-D.** Data shown as mean ± S.D. $^{*}P \leq 0.05$. Scale bars, 25 μm.

### Supplementary Figure 4

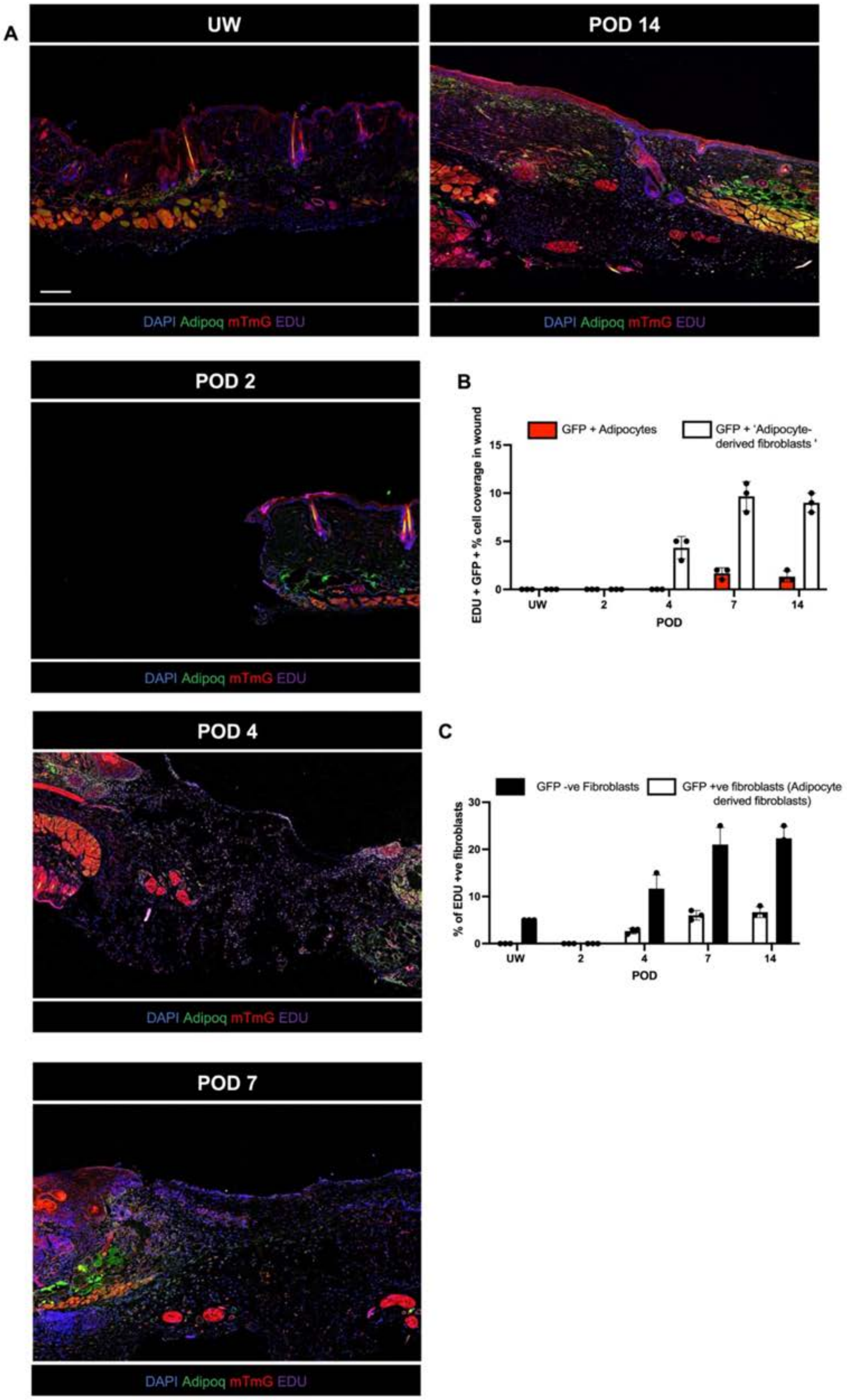

**Figure S4: EdU analysis of cell proliferation in *Adipoq*$^{\textit{Cre-ERT}}$*;R26*$^{\textit{mTmG}}$ wounds. A.** Labeling of proliferating cells in unwounded (UW) skin and wounds at indicated timepoints using EdU (purple signal); DAPI, nuclear counterstain (blue signal); mTmG, Tomato$^+$ (*Adipoq* lineage-negative) cells (red signal); Adipoq, GFP$^+$ (*Adipoq* lineage-positive) cells (green signal). Scale bars, 100 μm. **B.** Quantification of EdU signal colocalization with adipocytes (GFP$^+$Col1$^-$) and ADFs (GFP$^+$Col1$^+$) from histology. **C.** Quantification of percentage of non-adipocyte-derived (GFP$^-$) and adipocyte-derived (GFP$^+$) fibroblasts (Lin$^-$) positive for EdU incorporation by fluorescence-activated cell sorting (FACS).

**B-C.** Data shown as mean ± S.D.

### Supplementary Figure 5

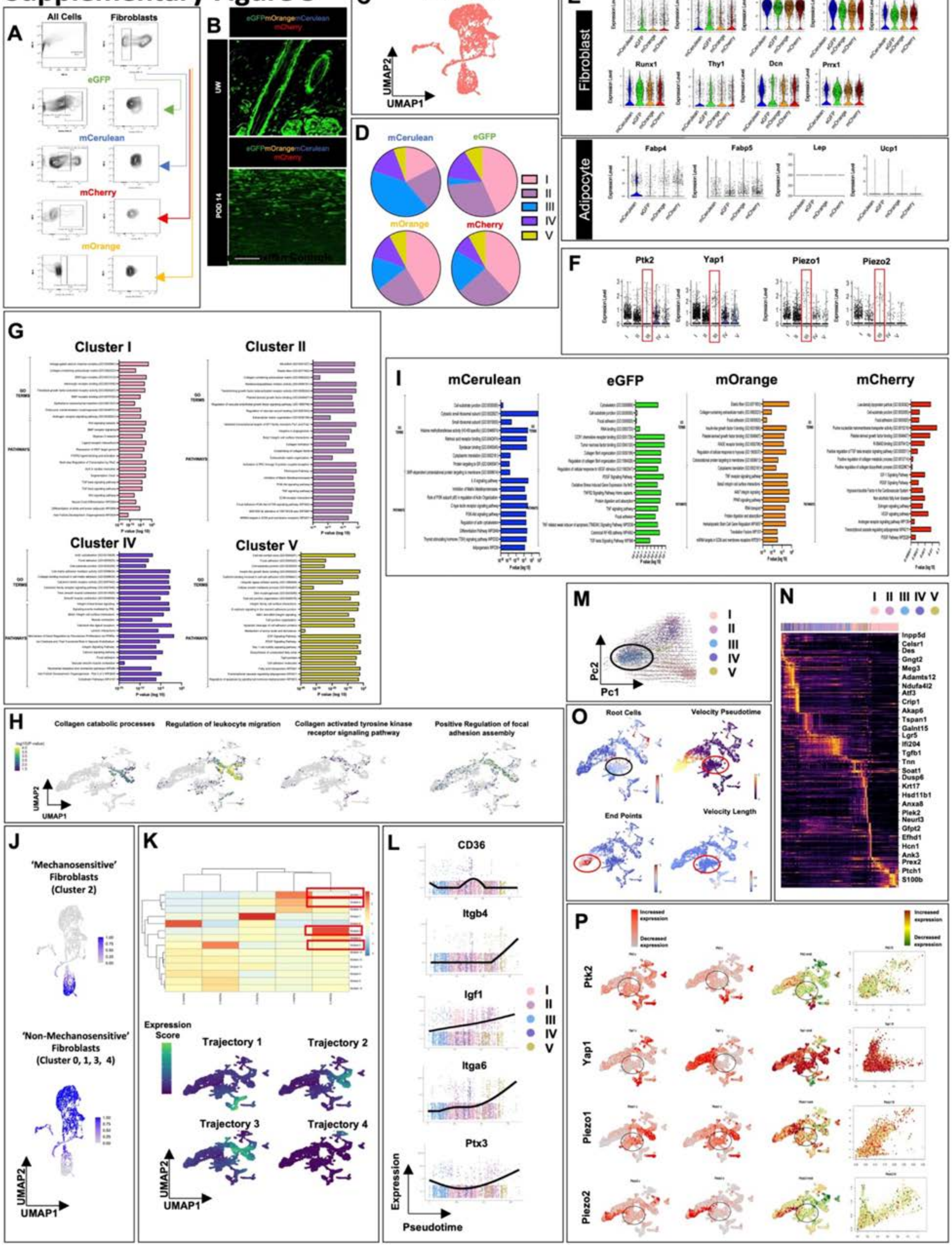

**Figure S5: Analysis of Rainbow ADF scRNA-seq data. A.** Fluorescence-activated cell sorting (FACS) strategy for sorting fibroblasts (Lin$^{-}$) of each Rainbow reporter color (eGFP, mCerulean, mCherry, or mOrange). **B.** Fluorescent histology of non-tamoxifen-induced Rainbow control specimens showing only eGFP expression (green signal) and absence of other Rainbow clone colors (mOrange, mCerulean, or mCherry; orange, blue, or red signal, respectively) that would indicate nonspecific recombination in the absence of tamoxifen induction. Scale bar, 25 µm. **C.** UMAP of Rainbow scRNA-seq fibroblasts colored by cell type (SingleR) demonstrating that all cells had a fibroblast transcriptomic identity. **D.** Quantification of relative representation of cells belonging to each Seurat fibroblast cluster (I-V) by Rainbow clone color. **E.** Violin plots showing expression of known fibroblast (top) and adipocyte (bottom) genes for cells of each Rainbow color (eGFP, non-adipocyte lineage-derived; mCerulean, mOrange, and mCherry, adipocyte-derived). **F.** Expression of mechanical signaling genes by Seurat cluster identity. **G.** GO pathway analysis for Seurat fibroblast clusters I, II, IV, and V. **H.** GeneTrail analysis for Seurat fibroblast clusters I-V showing comparable enrichment for fibrosis-related processes in clusters I, II, IV, and V. **I.** Gene Ontology (GO) pathway analysis for cells based on Rainbow color. **J.** Rainbow fibroblast UMAP showing anchor-based label transfer mapping of embedded cells from "mechanosensitive" and "non-mechanosensitive" fibroblast populations onto those previously defined by Foster et al. (Foster et al., 2021) **K.** Top, clustergram of pseudotime gene modules, with gene modules 1-4 indicated in red boxes in top panel. Bottom, expression scores of pseudotime modules 1-4 representing trajectories starting from cluster III. **L.** Expression of genes from pseudotime gene modules 1-4 across pseudotime (indicated in red boxes in top panel **K**). Overlaid lines represent regression fit to gene expression of cells over pseudotime; colors indicate Seurat cluster identity of each cell. **M.** scVelo analysis of cells along the first two principal components, demonstrating differentiation trajectory stemming from Seurat cluster III (black circle). **N.** scVelo heatmap highlighting genes with high correlation with velocity pseudotime, indexed by Seurat clusters. **O.** Top and bottom left, scVelo analysis of root and end point cells (black and red circles highlight root and end point cells, respectively). Top right, scVelo analysis of velocity pseudotime showing trajectory starting from cluster III (red circle). Bottom right, scVelo analysis of velocity length showing trajectory starting from cluster III (red circle). **P.** Gene-level scVelo analysis of specific genes of interest, depicting differences between spliced and unspliced RNA counts.

#### Supplementary Figure 6

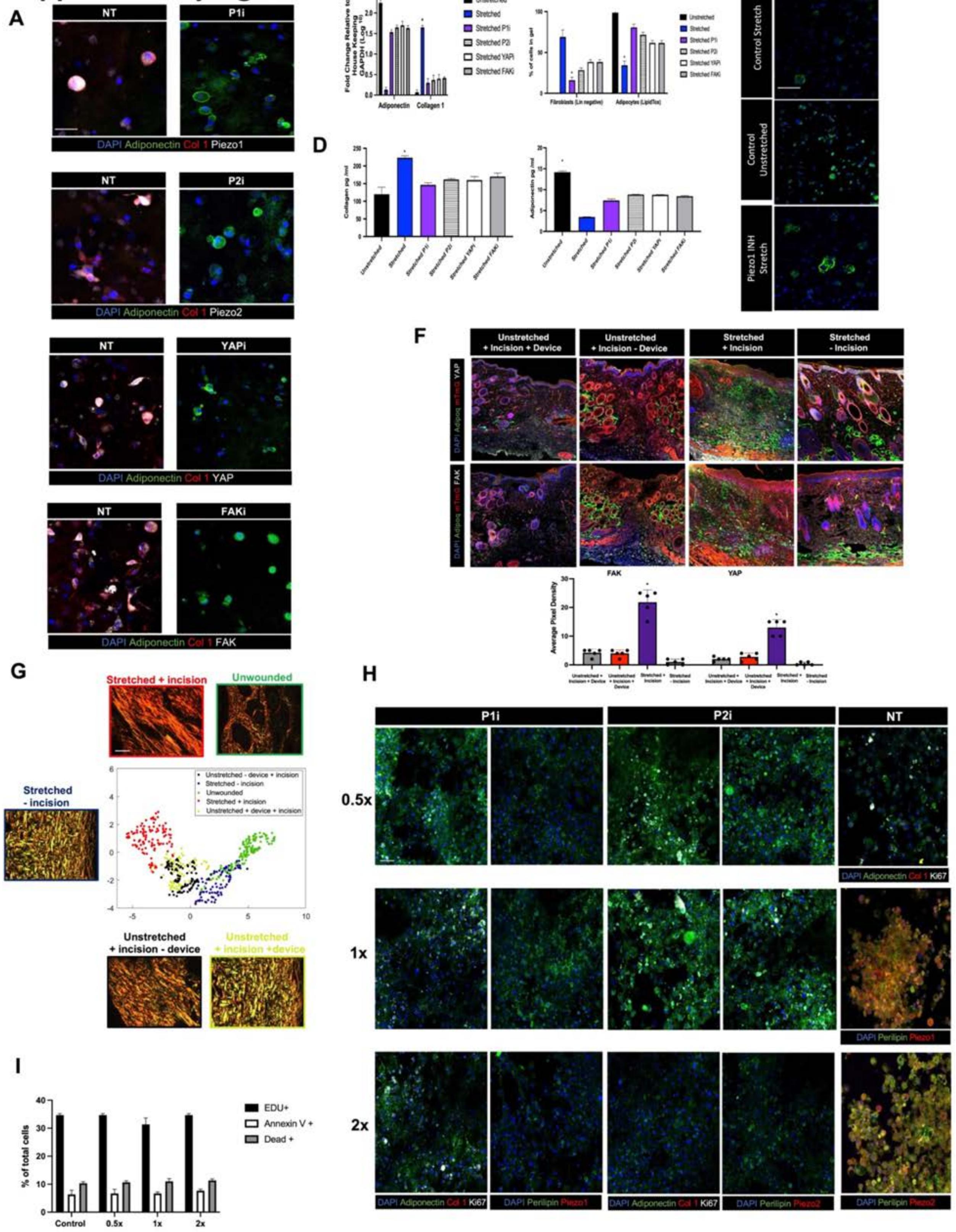

**Figure S6: Analysis of mechanosignaling activity and inhibition in mouse adipocytes *in vitro* and *in vivo.*** **A.** Immunofluorescence (IF) staining of mouse adipocytes cultured with mechanical stretching, with or without (no treatment, NT) indicated mechanosignaling inhibitors. IF staining for adiponectin (green fluorescent signal), Col 1 (red signal), and indicated mechanosignaling markers (white signal). **B.** Quantification of adiponectin and Col 1 expression by RT-qPCR. **C.** Quantification of percent of cultured cells that are fibroblasts (Lin$^{-}$) or adipocytes (LipidTox$^{+}$) by FACS. **D.** Enzyme-linked immunosorbent assay (ELISA) quantification of collagen (left) or adiponectin (right) protein expression in cultured cells. **E.** Lipid Tox staining of mouse adipocytes with stretch (top), unstretched (middle), and with P1i inhibitor (bottom). **F**. Top, fluorescent histology of mouse hypertrophic scarring (HTS) model wounds and skin, indicated conditions; mTmG, Tomato$^{+}$ (*Adipoq* lineage-negative) cells (red signal); Adiponectin, GFP$^{+}$ (*Adipoq* lineage-positive) cells (green signal); IF staining for FAK (top row) or YAP (bottom row; both, white signal). Bottom, quantification of FAK (left) and YAP (right) expression from IF staining. **G.** Picrosirius red staining of skin and wounds from mouse HTS model experimental conditions (surrounding panels) and UMAP of quantified extracellular matrix (ECM) ultrastructure parameters based on picrosirius red histology (central panel; each dot represents one histologic image). **H.** IF staining of mouse adipocytes cultured under varying concentrations of mechanosignaling inhibitors (see Methods for specific concentrations) or NT control, with IF staining for adiponectin (green signal), Col 1 (red signal), Ki67 (gray signal), perilipin (green signal), Piezo1 (red signal), and Piezo2 (red signal) as indicated. **I.** Percentage of all cells positive for EdU, annexin V (apoptosis marker), or DAPI (dead cells) by FACS of mouse adipocytes treated with indicated varying concentrations of Piezo1 inhibitor (P1i) including 0.5X, 1X, and 2X. None of the markers' expression was significantly different between any of the inhibitor concentrations or compared to control ($P > 0.05$).

**B-E, H.** Data shown as mean ± S.D. **B-E.** *$P \leq 0.05$ (vs. all other conditions). Scale bars, 10 μm (**A**), 100 μm (**E**), 150 μm (**F**), 100 μm (**G**).

### Supplementary Figure 7

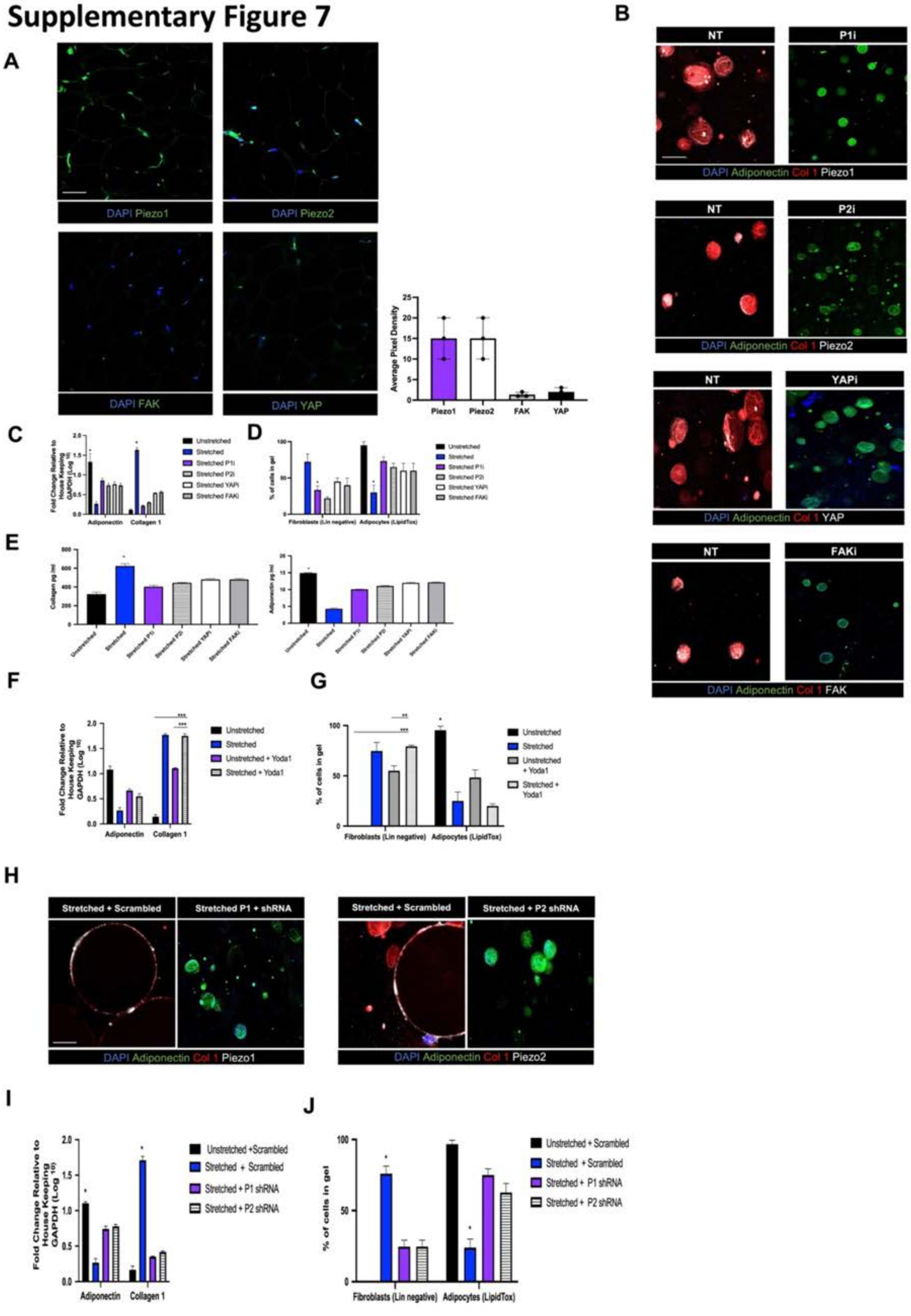

**Figure S7: Analysis of mechanosignaling activity and inhibition in human adipocytes *in vitro*.**

**A.** Left, immunofluorescence (IF) staining of human adipose tissue with IF staining for indicated mechanosignaling markers (green signal). Right, quantification of expression of each marker from IF. **B-E.** As in **S6A-D**, but with human adipocytes. **F, G.** As in **C, D**, but with or without Yoda1 (Piezo1 agonist) treatment. **H.** IF staining of cultured human adipocytes treated with indicated shRNA with IF staining for adiponectin (green signal), Col 1 (red signal), and Piezo1 or Piezo2 as indicated (white signal). **I, J.** As in **C, D**, but with indicated shRNA treatments.

**A, C-G, I-J.** Data shown as mean ± S.D. **C-G, I-J.** $^{*}P \leq 0.05$, $^{**}P \leq 0.01$, $^{***}P \leq 0.001$. P-value reflects pairwise comparison between indicated conditions, or comparison of one condition vs. all other conditions when specific pairwise comparison not indicated. Scale bars, 100 µm (**A, B, H**).

#### Supplementary Figure 8

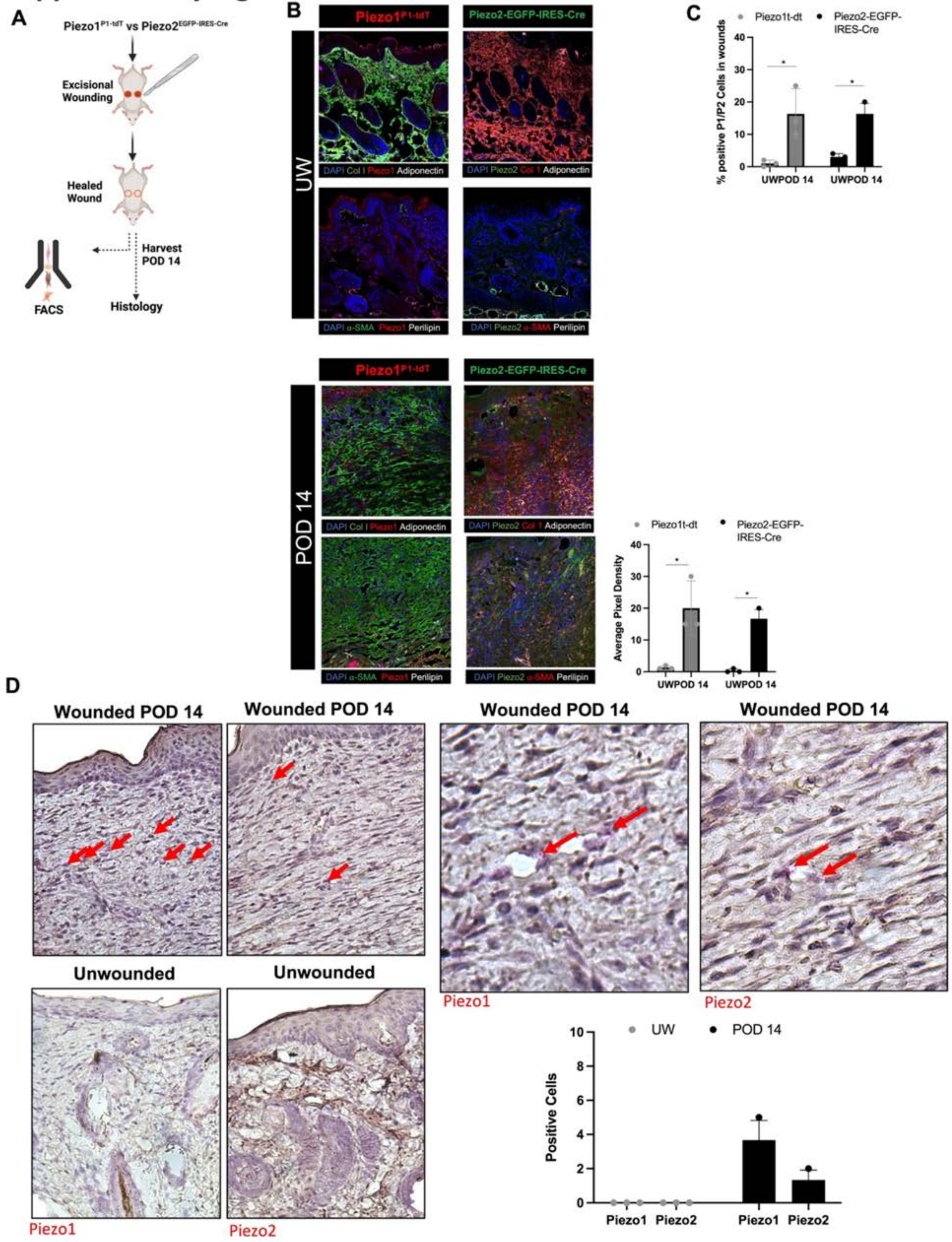

**Figure S8: *Piezo1* and *Piezo2* lineage tracing in wounds. A.** Schematic of *Piezo1*$^{t\text{-}dT}$ and *Piezo2*$^{EGFP\text{-}IRES\text{-}Cre}$ wounding experiments to genetically trace *Piezo1* and *Piezo2* lineage-positive cells in wounds. **B.** Fluorescent histology of unwounded (UW) skin (top) and POD 14 wounds (bottom left) with immunofluorescence (IF) staining for fibroblast and adipocyte markers. First column: Piezo1, tdTomato signal labeling *Piezo1* lineage-derived cells (red signal); IF for Col 1 or α-smooth muscle actin (α-SMA) (green signal), adiponectin or perilipin (white signal). Second column: Piezo2, EGFP signal labeling *Piezo2* lineage-derived cells (green signal); IF for Col 1 or α-SMA (red signal), adiponectin or perilipin (white signal). Bottom right, quantification of *Piezo1* (tdTomato$^{+}$) and *Piezo2* (EGFP$^{+}$) lineage-derived cells in respective lineage-tracing models by fluorescent signal on histology, UW skin vs. POD 14 wounds. **C.** Fluorescence-activated cell sorting (FACS) quantification of percentage of cells of *Piezo1* or *Piezo2* lineage origin based on tdTomato or EGFP expression, respectively. **D.** Left panels, RNAscope *in situ* hybridization for Piezo1 (first column) and Piezo2 (second column; both, pink signal). Top right, higher-power zoom of Piezo1 (left) and Piezo2 (right) RNAscope. Red arrows indicate positively stained cells. Right bottom, quantification of Piezo1$^{+}$ and Piezo2$^{+}$ cells per HPF from RNAscope.

**B-D.** Data shown as mean ± S.D. **B, C.** $*P \leq 0.05$. Scale bars, 50 μm (**B**), 100 μm (**D**, left panels), and 20 μm (**D**, right panels).

Supplementary Figure 9

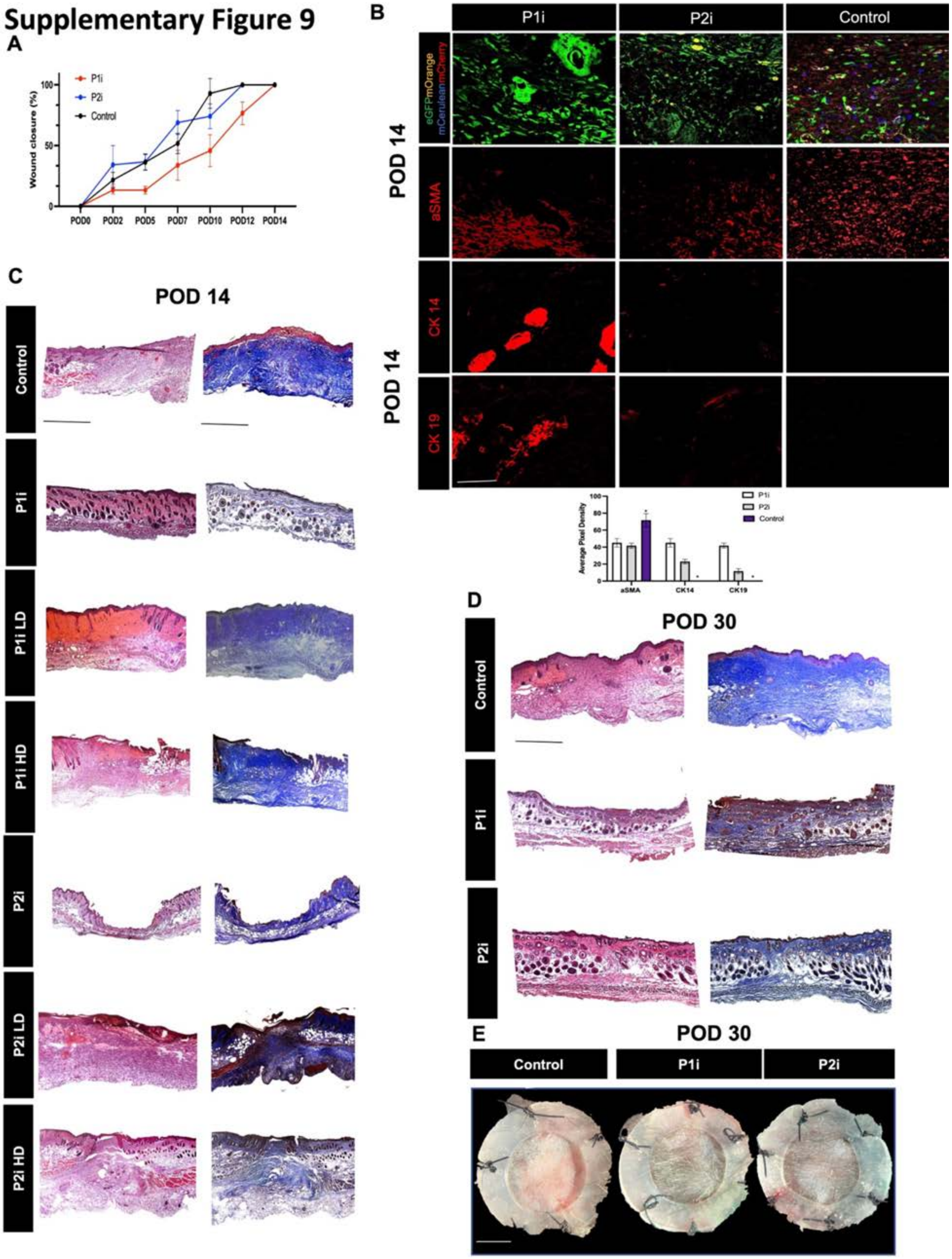

**Figure S9: Analysis of wounds treated with small molecule inhibitors of Piezo1 and Piezo2.** **A.** Wound curve showing percent of original wound area re-epithelialized at indicated timepoints in wounds treated with Piezo1 inhibitor (P1i), Piezo2 inhibitor (P2i), or PBS (control). **B.** Top, fluorescent histology of Rainbow wounds showing Rainbow clone colors (first row; eGFP, green signal; mOrange, orange signal; mCerulean, blue signal; mCherry, red signal) or immunofluorescence (IF) staining for α-smooth muscle actin (α-SMA) (second row), CK14 (third row), or CK19 (fourth row; all IF, red signal, not consecutive sections). Bottom, quantification of expression of indicated markers from IF. $^*P \leq 0.05$ vs. all other conditions. **C, D.** H&E staining (left) and Masson's trichrome staining (right) of POD 14 (**C**) and POD 30 (**D**) wounds with indicated treatments (P2iLD; P2i Low dose, P2iHD; P2i high dose, P1iLD; P1iLow dose, P1iHD; P2iLow dose). (**E**). Gross photos of wounds at POD 30 treated with PBS, P1I or P2I.

**A, B.** Data shown as mean ± S.D. Scale bars, 25 μm (**B**), 500 μm (**C, D**) 6mm (**E**).

#### Supplementary Figure 10

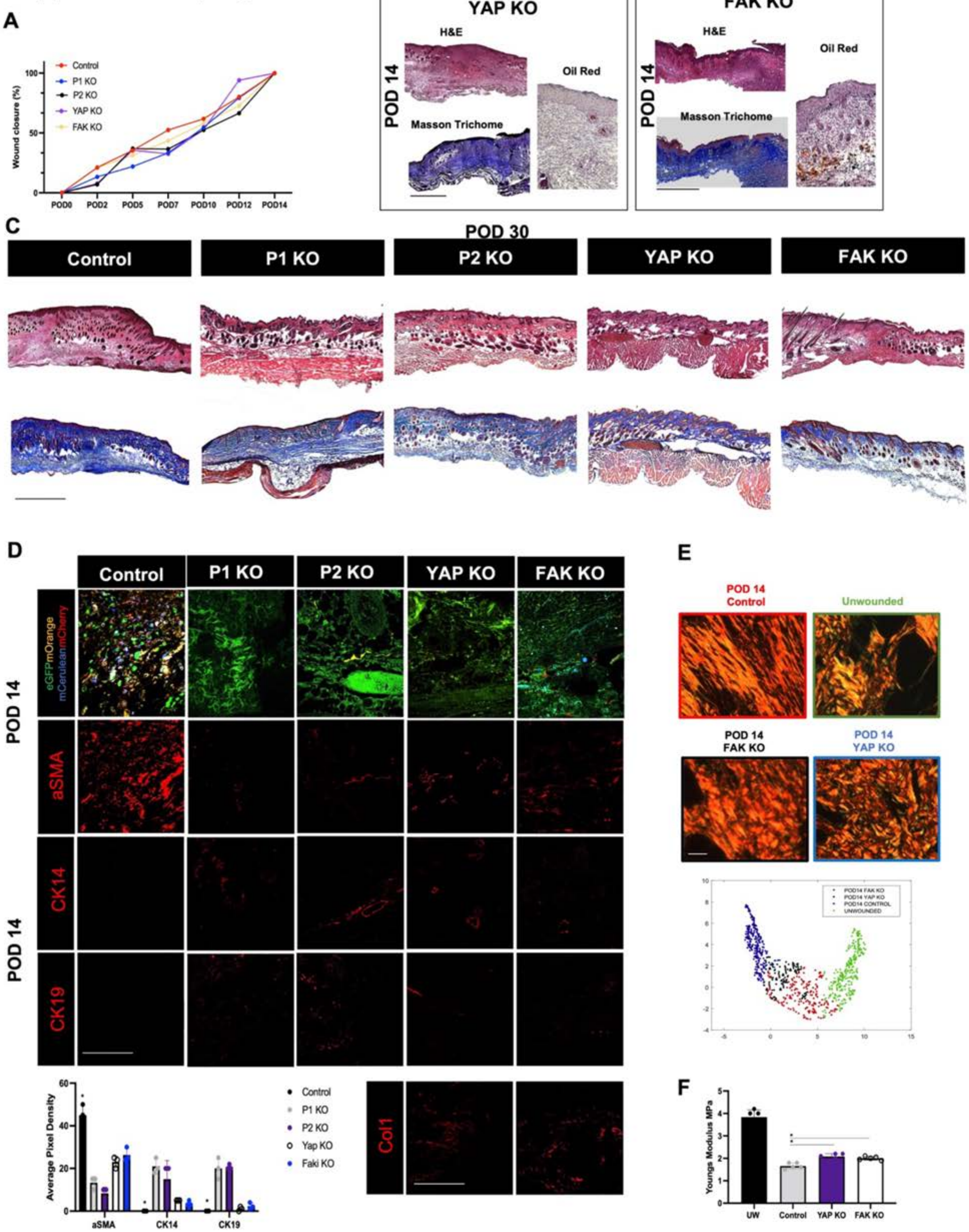

**Figure S10: Analysis of wounds treated with adipocyte-targeted genetic knockout of mechanosignaling genes. A.** Wound curve showing percent of original wound area re-epithelialized at indicated timepoints in wounds with adipocyte-targeted Piezo1 (P1), Piezo2 (P2), YAP, or FAK knockout (KO). **B.** Histology of YAP (left) and FAK (right) KO wounds, each showing H&E (top left), Masson's trichrome (bottom left), and Oil Red O (bottom right) staining. **C.** H&E (top) and Masson's trichrome (bottom) staining of indicated wound conditions at POD 30. **D.** Fluorescent histology of Rainbow wounds with P1, P2, YAP, or FAK KO or control at POD 14, showing Rainbow clone colors (first row; eGFP, green signal; mOrange, orange signal; mCerulean, blue signal; mCherry, red signal) or IF staining for α-SMA (second row), CK14 (third row), CK19 (fourth row, or Col1 (bottom row right; all IF, red signal, not consecutive sections). Bottom left, quantification of expression of indicated markers by IF. **E.** Top, picrosirius red staining of unwounded (UW) skin and control, FAK KO, and YAP KO wounds at POD 14; bottom, UMAP of quantified extracellular matrix (ECM) ultrastructure parameters based on picrosirius red histology (each dot represents one histologic image). **F.** Young's modulus, calculated from tensile strength testing, by wound condition.

**A, D, F.** Data shown as mean ± S.D. **D, F.** $^{*}P \leq 0.05$. P-value reflects pairwise comparison between indicated conditions, or comparison of one condition vs. all other conditions when specific pairwise comparison is not indicated. Scale bars, 500 μm (**B**, H&E and trichrome), 25 μm (**B**, Oil Red O), 500 μm (**C**), 25 μm (**D**), 20 μm (**E**).

#### Supplementary Figure 11

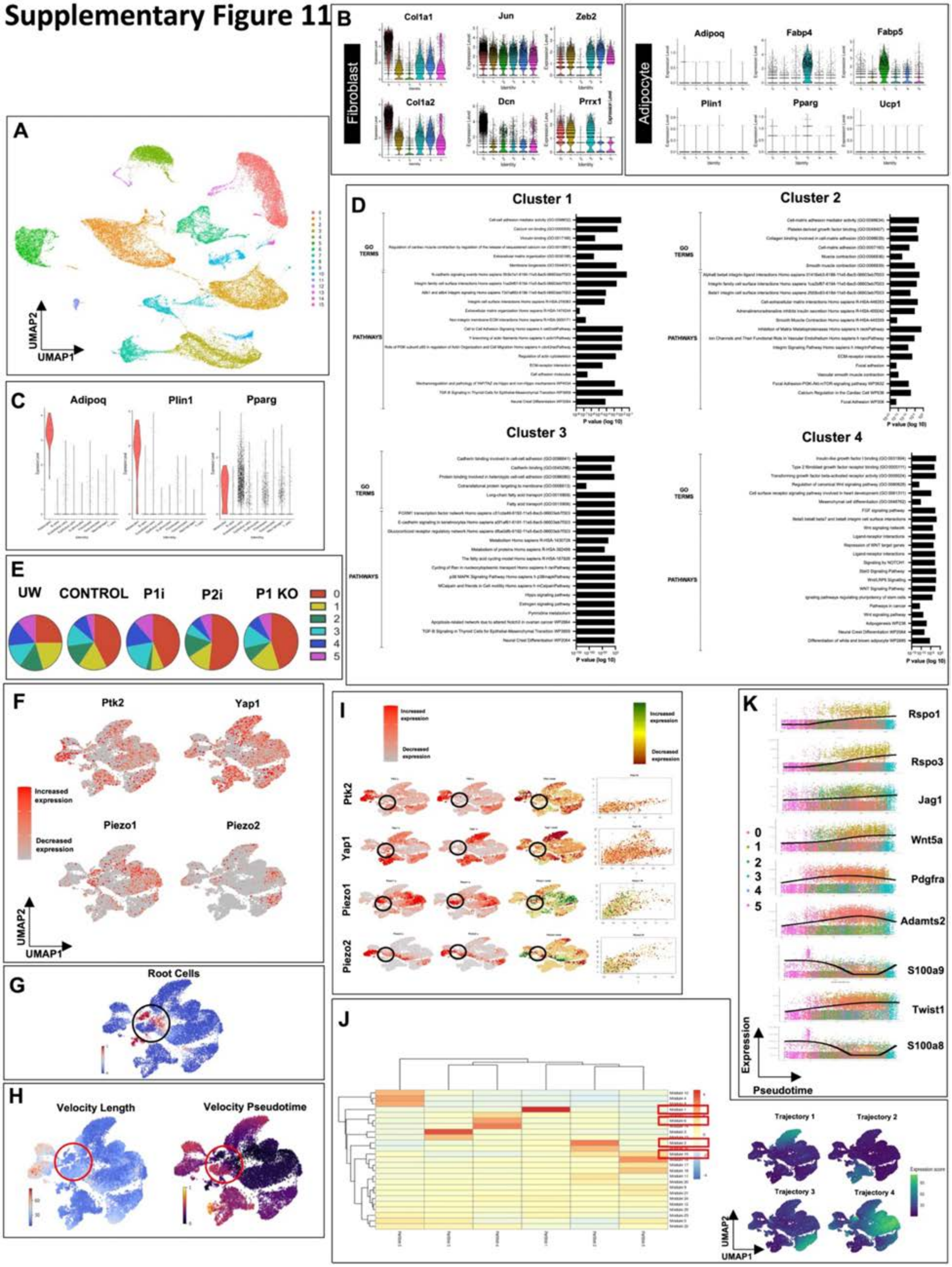

**Figure S11: scRNA-seq analysis of wounds with and without Piezo inhibition. A.** UMAP of all wound cells colored by Seurat cluster (dataset from Fig. 5). **B.** Violin plots showing expression of known fibroblast (left) and adipocyte (right) genes for cells of each fibroblast Seurat cluster (0-5). **C.** Violin plots showing expression of indicated characteristic adipocyte genes by cell type within scRNA-seq dataset. **D.** Gene ontology (GO) pathway analysis for Fig. 5 Seurat fibroblast clusters 1, 2, 3, and 4. **E.** Quantification of relative representation of cells belonging to each Seurat fibroblast cluster (0-5) by experimental condition. **F.** Fibroblast UMAP colored by expression of select mechanosignaling genes. **G.** scVelo analysis of root cells showing trajectory starting at cluster 5 (black circle). **H**. Left, scVelo velocity length analysis showing the trajectory starting at cluster 5 (red circle). Right, scVelo velocity pseudotime anlaysis showing the trajectory starting at cluster 5 (red circle). **I.** Gene-level scVelo analysis of specific genes of interest, depicting differences between spliced and unspliced RNA transcripts. **J.** Left, clustergram of pseudotime gene modules, with gene modules 1-4 indicated in red boxes. Right, expression scores of pseudotime modules 1-4 representing trajectories starting from cluster 5. **K**. Expression of genes from pseudotime gene modules 1-4 (indicated in red boxes in left panel of **J**) across pseudotime. Overlaid lines represent regression fit to gene expression of cells over pseudotime; colors indicate Seurat cluster identity of each cell.

#### Supplementary Figure 12

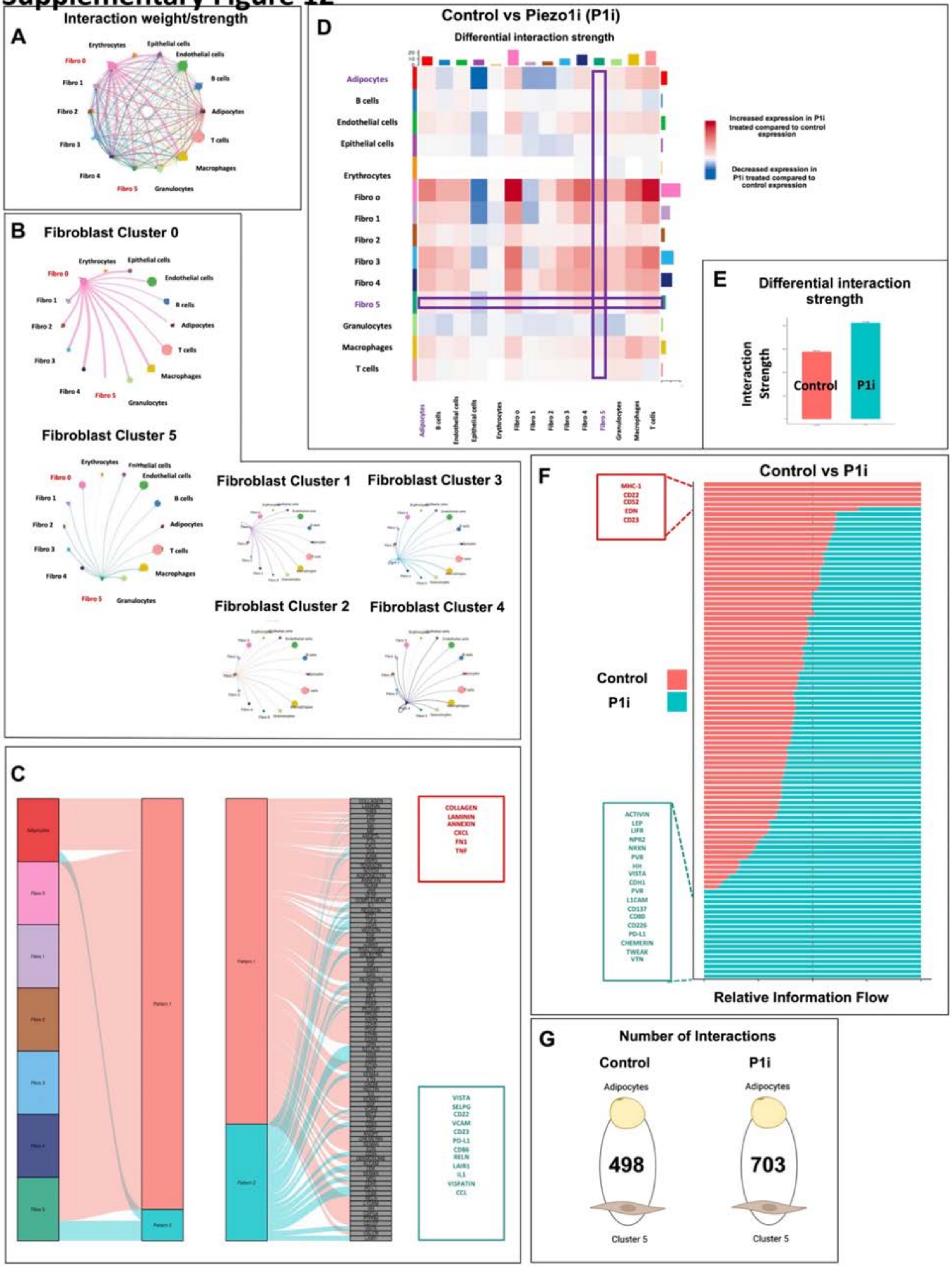

**Figure S12: CellChat analysis of cell-cell interactions by wound condition from scRNA-seq dataset. A.** Interaction weight/strength of cell-cell interactions between all sequenced cells in unwounded skin and wounds at POD 14 (control, Piezo1 inhibitor [P1i], Piezo2 inhibitor [P2i], and P1 knockout [KO] wounds). **B.** Circle plot of cell-cell interactions for each fibroblast cluster. **C.** River plot of outgoing communication from cells in unwounded (UW) and wounded skin at POD 14. **D.** Heatmap illustrating cell-cell interactions in P1i compared to control wound cells (blue shades, decreased cell-cell signaling in P1i compared to control wounds; red, increased cell-cell signaling in P1i compared to control wounds). Purple boxes highlight increased overall interactions between cluster 5 fibroblasts and other cell types. **E.** Bar chart showing differential interaction strength between control vs. P1i wound cells. **F.** Relative (control vs. P1i, normalized to add to 1) calculated information flow for indicated genes for all cell interactions following control vs. P1i treatment. **G.** Number of interactions between adipocytes and cluster 5 fibroblasts in control and P1i treated wounds.

### Supplementary Figure 13

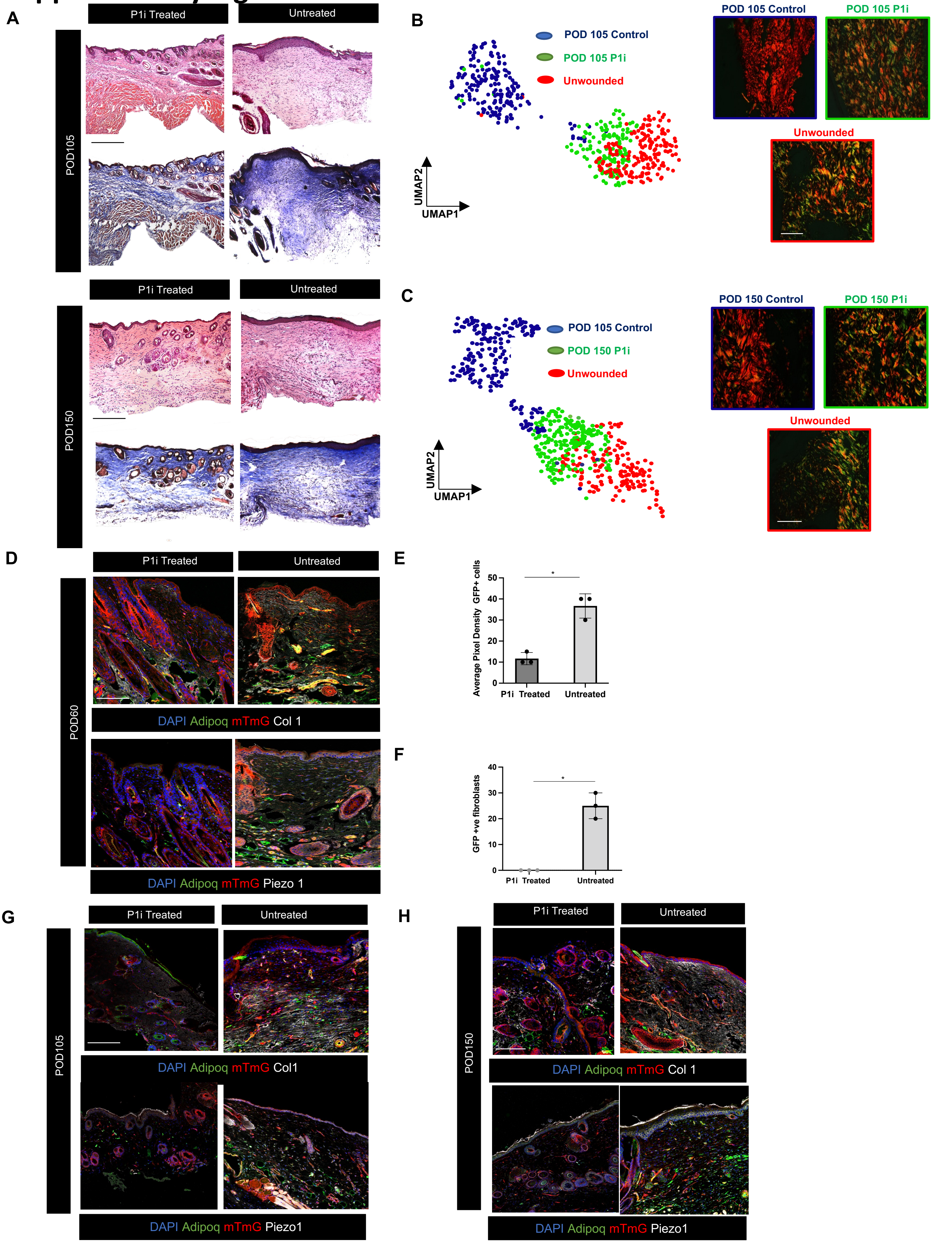

**Figure S13: Analysis of Piezo1 inhibition during scar rescue** **A**. H&E (top row) and Masson Trichome staining (bottom row) of P1i and PBS treated wounds at POD 105 (top) and POD 150 (bottom). **B.** Representative Picrosirius red staining image (right) and associated UMAP (left) following PBS or P1i treatment at P0D 105. **C**. Representative Picrosirius red staining image (right) and associated UMAP (left) following PBS or P1i treatment at P0D 150. **D.** Fluorescent histology of scars at POD 60 following PBS or P1i treatment showing immunofluorescence (IF) staining for Collagen type I (Col1) (top row) and Piezo1 (bottom row). **E**. Quantification of expression of indicated markers from IF. *$P \leq 0.05$ vs. all other conditions. **F.** Quantification of GFP cells at POD 60 scars treated with either P1i or PBS by flow cytometry. **G**. Fluorescent histology of scars at POD 105 following PBS or P1i treatment showing immunofluorescence (IF) staining for Collagen type I (Col1) (top row) and Piezo1 (bottom row) at POD 105. **H**. Fluorescent histology of scars at POD 150 following PBS or P1i treatment showing immunofluorescence (IF) staining for Collagen type I (Col1) (top row) and Piezo1 (bottom row) at POD 150. Scale bars, 250 μm (**A**, **D**, **G, H**), 25 μm (**C**).

#### Supplementary Figure 14

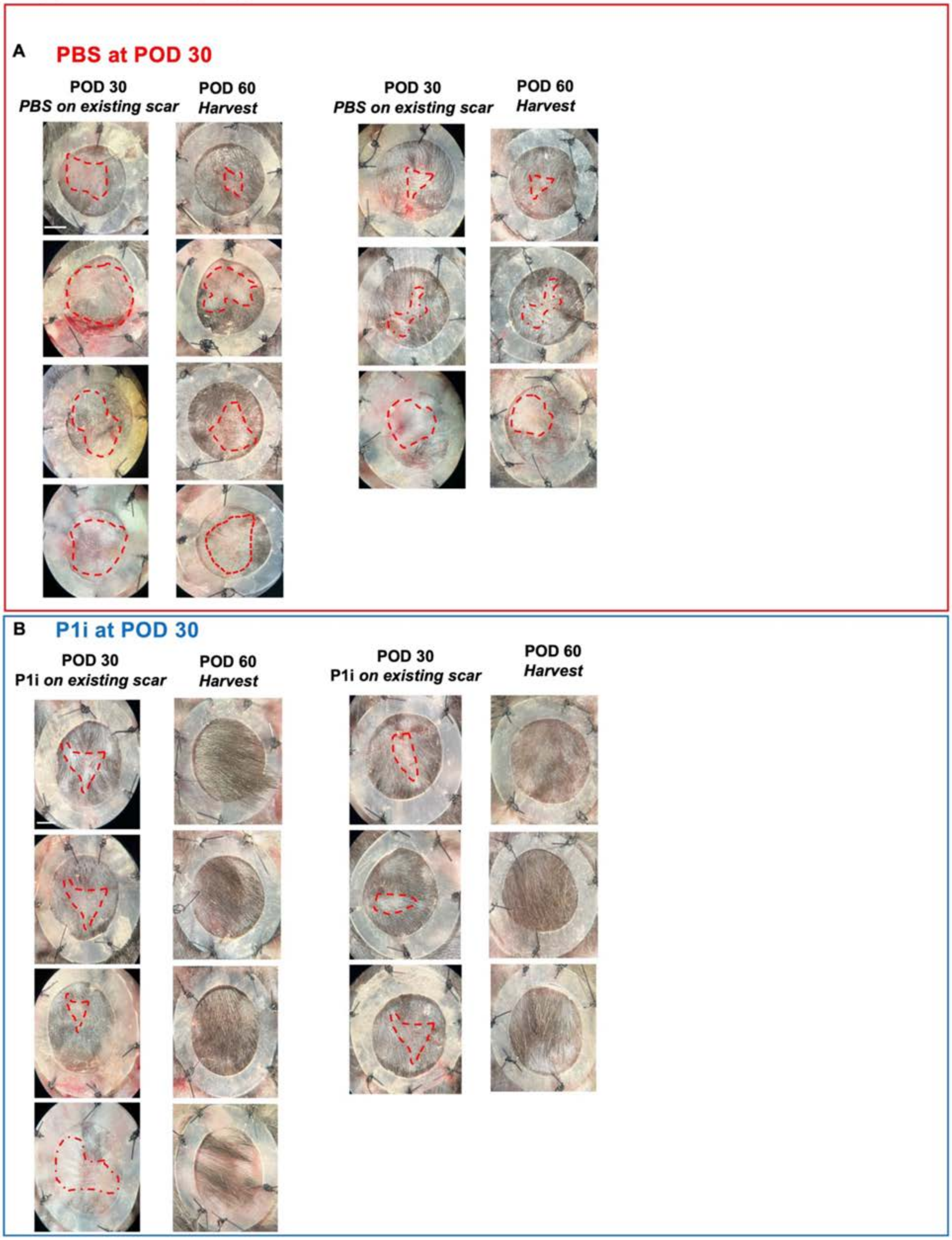

**Figure S14: Analysis of Piezo1 inhibition during scar rescue A.** Gross histology scars at POD 60 following PBS or **B.** P1i treatment (red dotted arrows illustrating the scars). All gross histology of wounds at POD 60 are shown. Scale bar = 3mm.

### Supplementary Figure 15

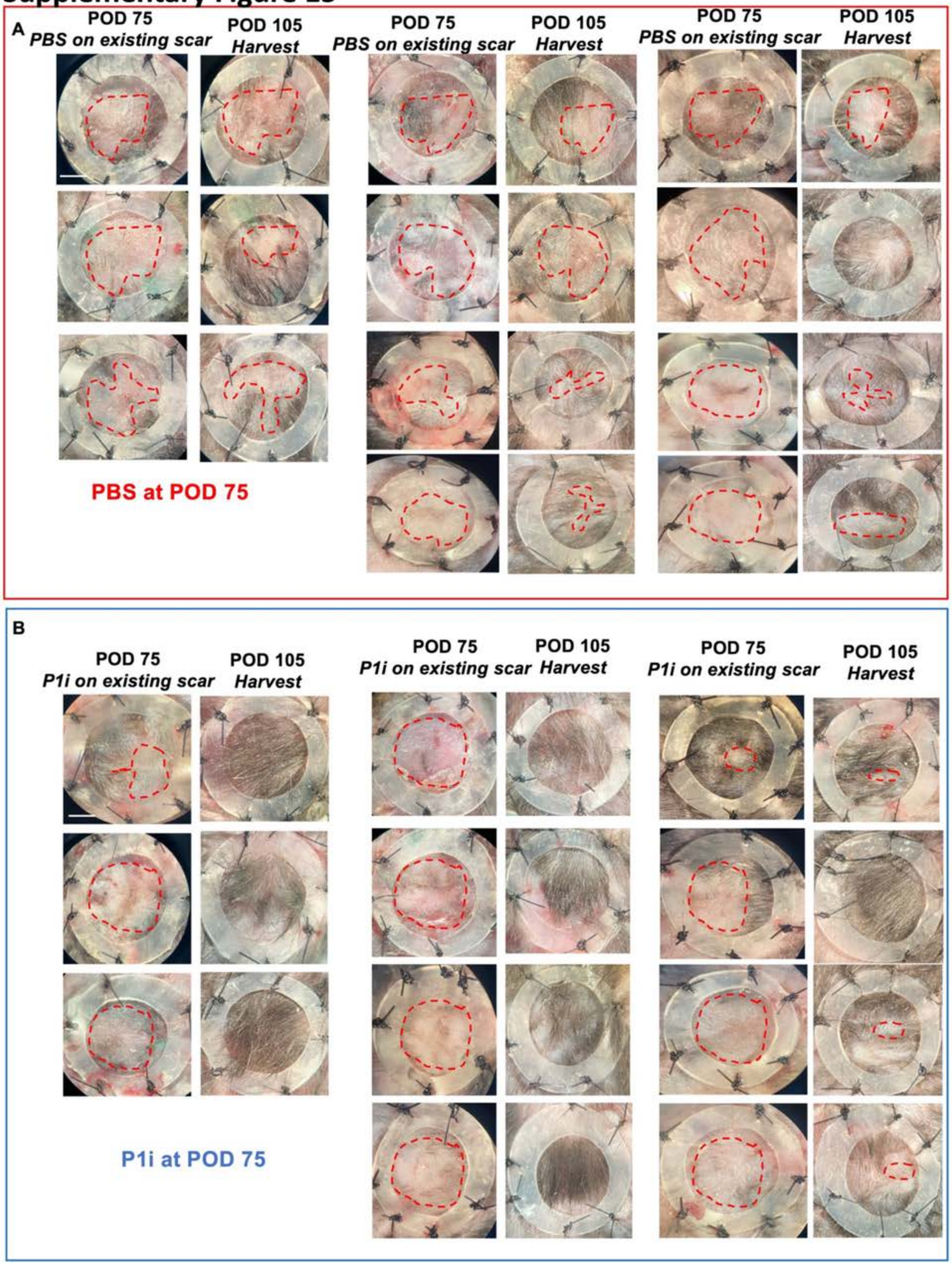

**Figure S15: Analysis of Piezo1 inhibition during scar rescue A.** Gross histology scars at POD 105 following PBS or **B.** P1i treatment (red dotted arrows illustrating the scars). All gross histology of wounds at POD 105 are shown. Scale bar = 3mm.

#### Supplementary Figure 16

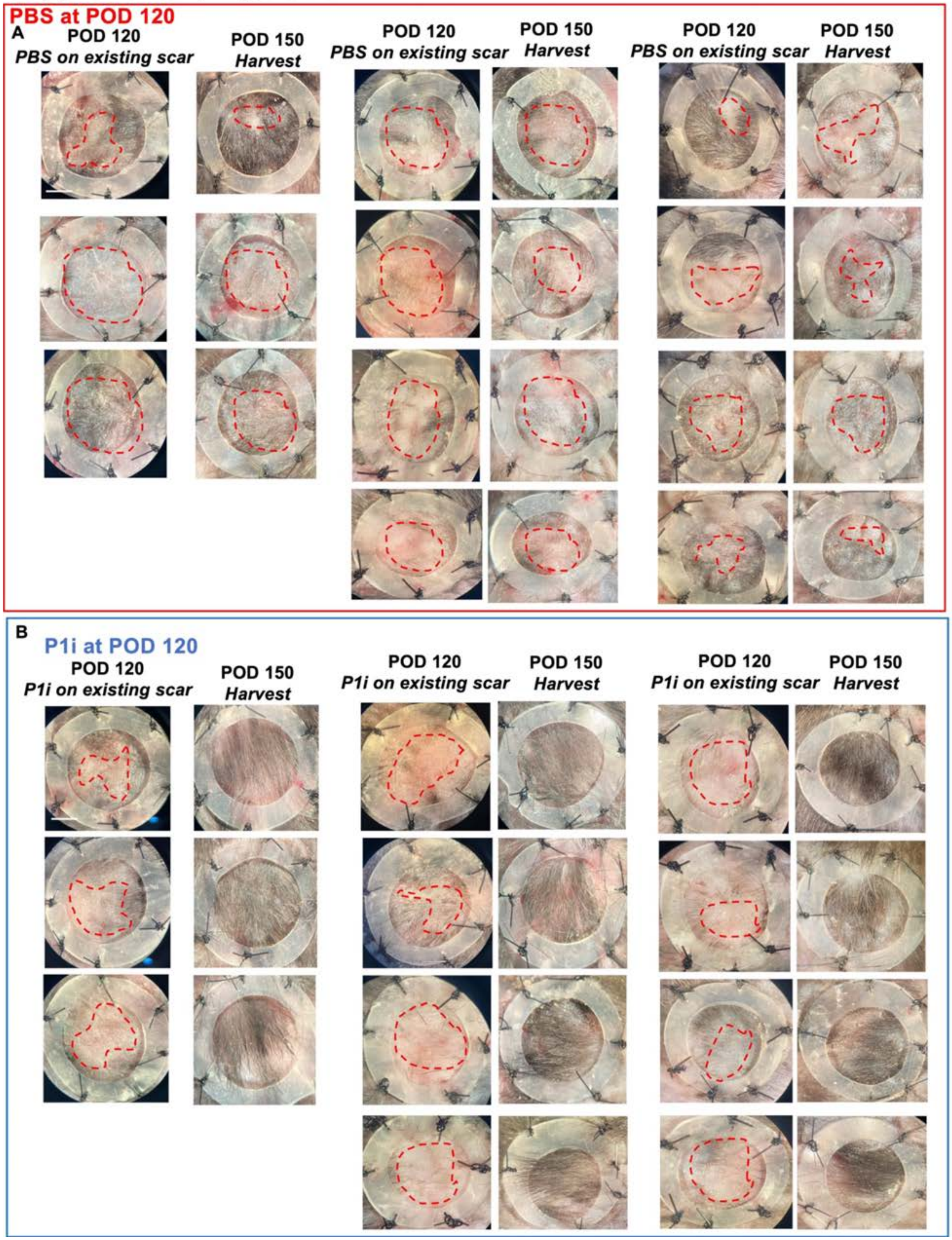

**Figure S16: Analysis of Piezo1 inhibition during scar rescue A.** Gross histology scars at POD 150 following PBS or **B.** P1i treatment (red dotted arrows illustrating the scars). All gross histology of wounds at POD 150 are shown. Scale bar = 3mm.

#### Supplementary Figure 17

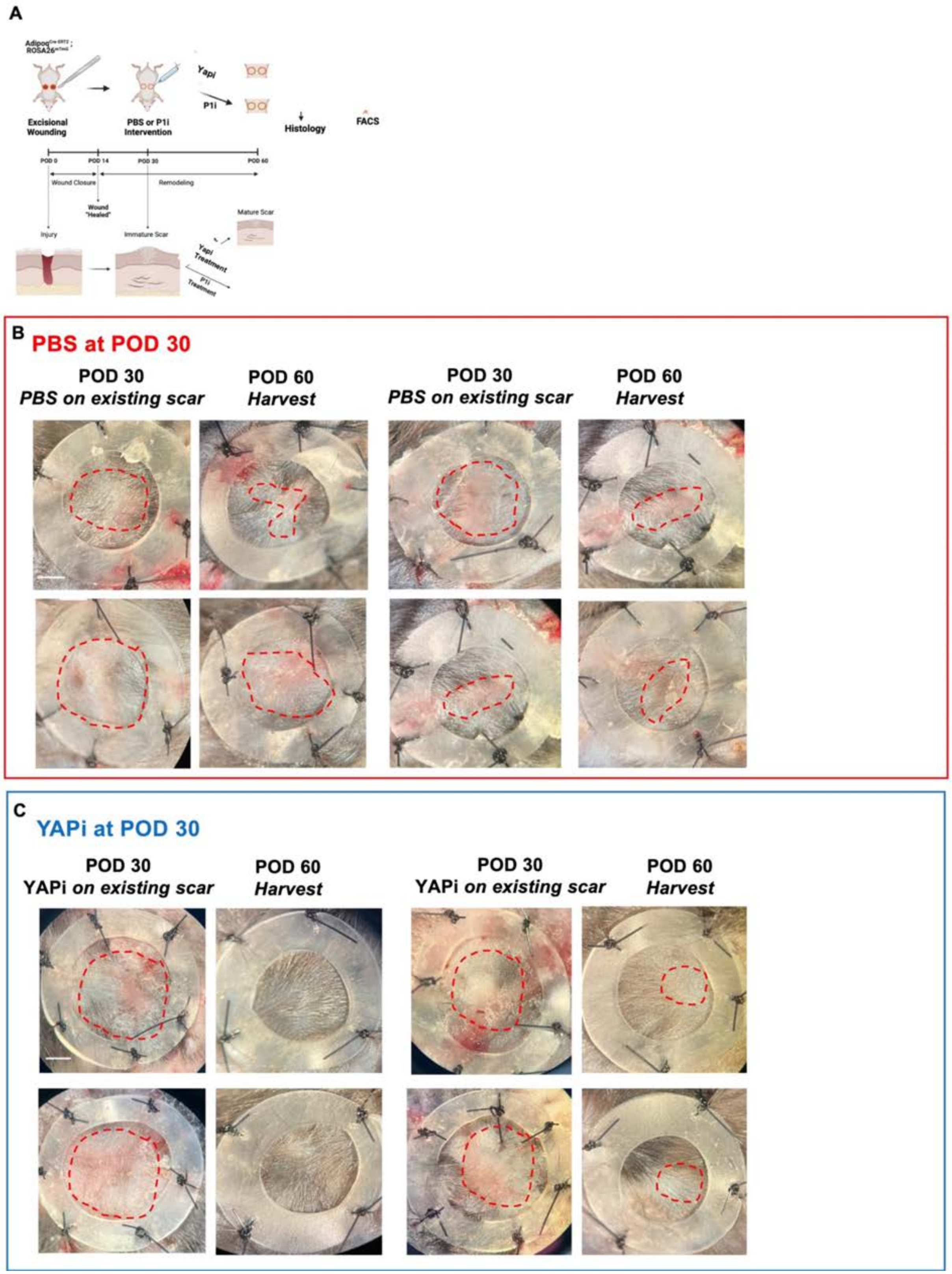

**Figure S17: Analysis of YAPi inhibition during scar rescue**. **A.** Schematic showing YAPi or PBS of scars at POD 30 and harvested at POD 60. **B.** Gross histology scars at POD 60 following PBS or **B.** YAPi treatment (red dotted arrows illustrating the scars). Scale bar = 3mm.

### Supplementary Figure 18

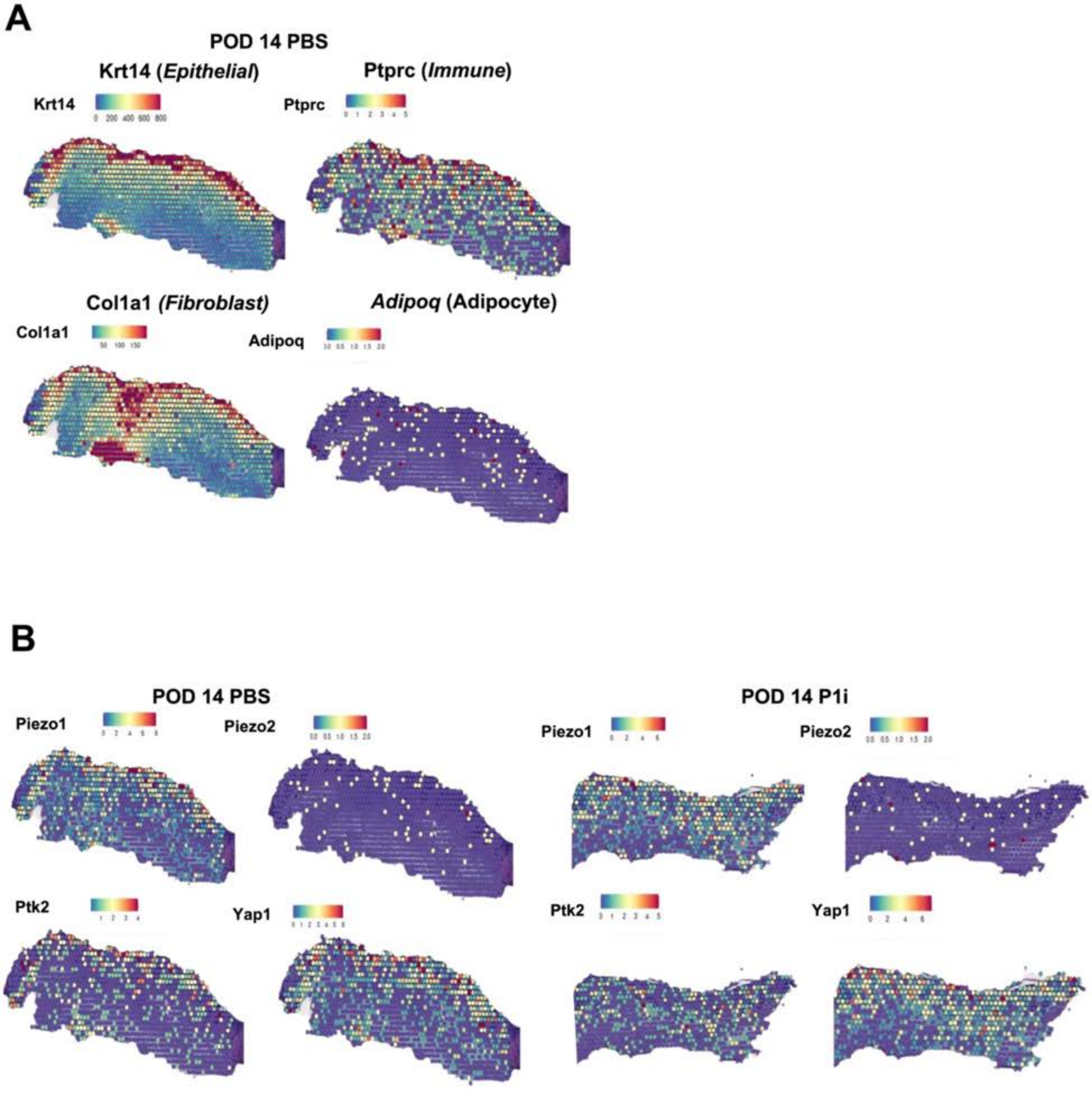

**Figure S18: VISIUM analysis at POD 14**. **A.** Spatial plots of epithelial (Krt14), immune (Ptprc), fibroblast (Col1a1) and adipocyte (Adipoq) cell markers at POD 14 following PBS treatment. **B.** Spatial plots of mechanical markers (Piezo1, Piezo2, Ptk2, and Yapi) at POD14 following PBS treatment (left) and P1i treatment (right).

#### Supplementary Figure 19

A

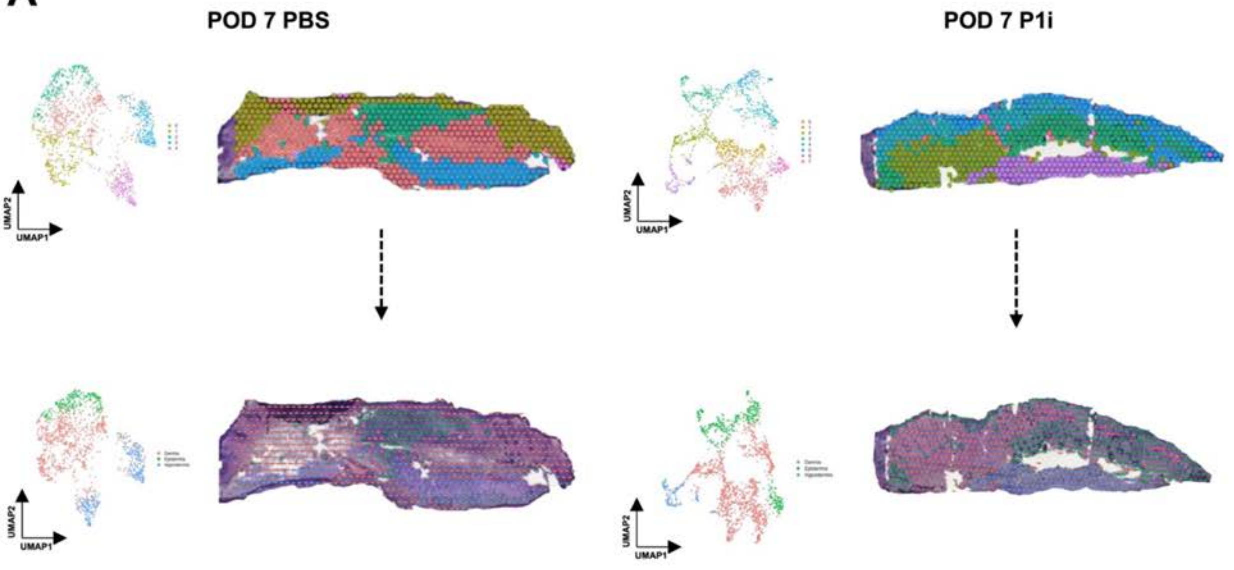


B

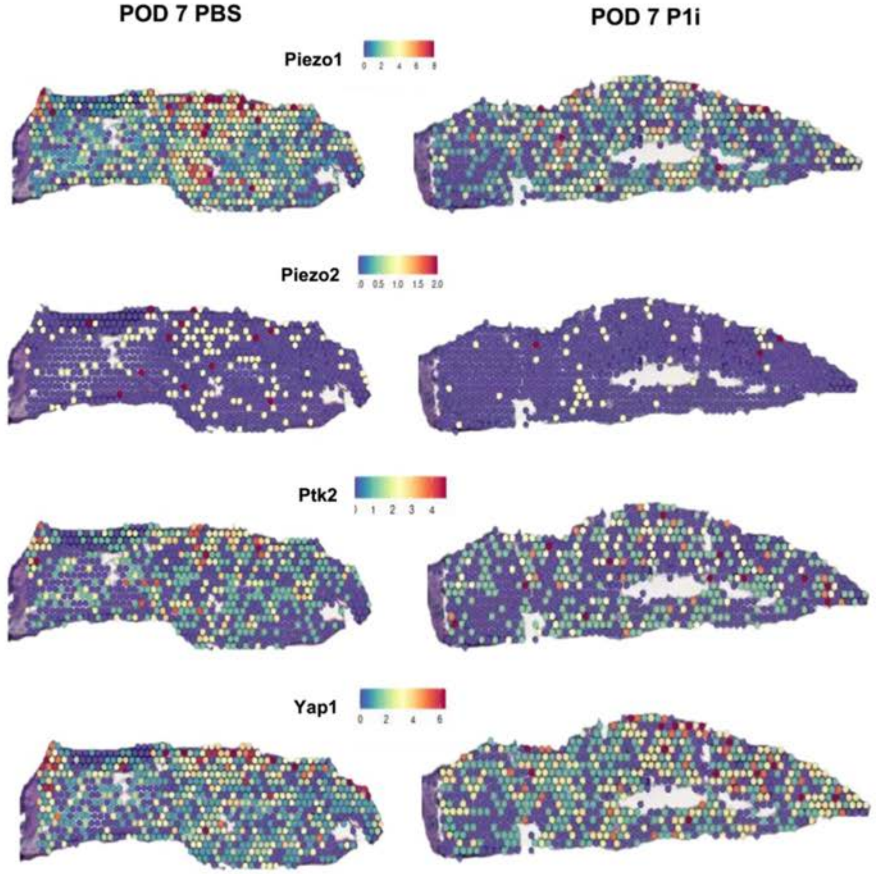

**Figure S19: VISIUM analysis at POD 7**. **A.** Spatial plots of POD 7 PBS (left) or P1i treated (right) wounds colored by Seurat clusters (top row) with associated UMAPs (left). Spatial plots of POD 7 PBS (left) or P1i treated (right) wounds colored by epithelial, dermal and hypodermis category (right) with associated UMAP (left). **B.** Spatial plots of mechanical markers at POD 7 (Piezo1, Piezo2, Ptk2, and Yapi) following PBS treatment (left column) and P1i treatment (right column).

#### Supplementary Figure 20

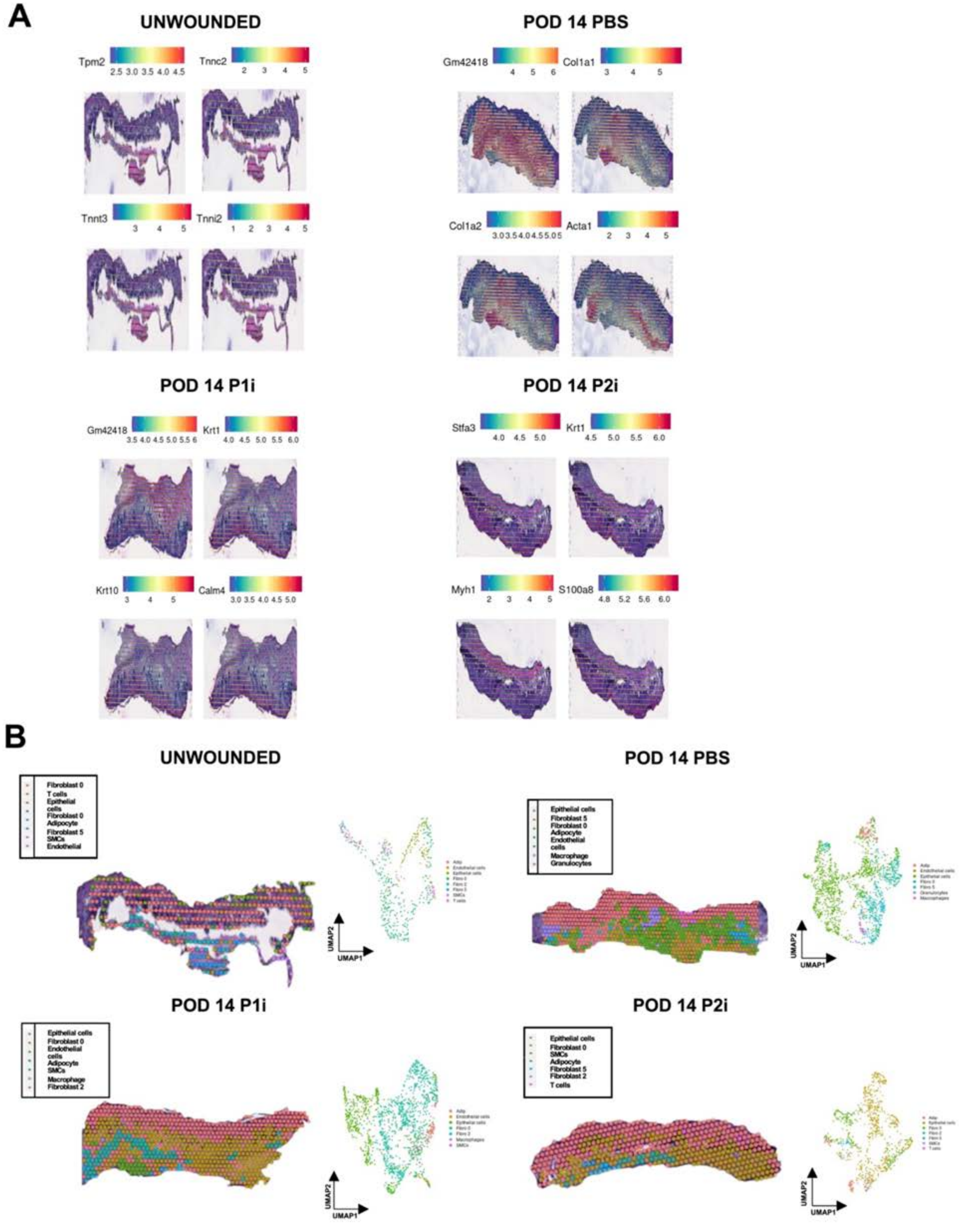

**Figure S20: VISIUM analysis at POD 14**. **A.** Spatial plots of spatially variable features at POD 14 following PBS treatment (top right), unwounded (top left), following P1i treatment (bottom left) and P2i treatment (bottom right). **B.** Spatial plots of cell types maximally predicted from scRNAseq *Fig.5* data.

#### Supplementary Figure 21

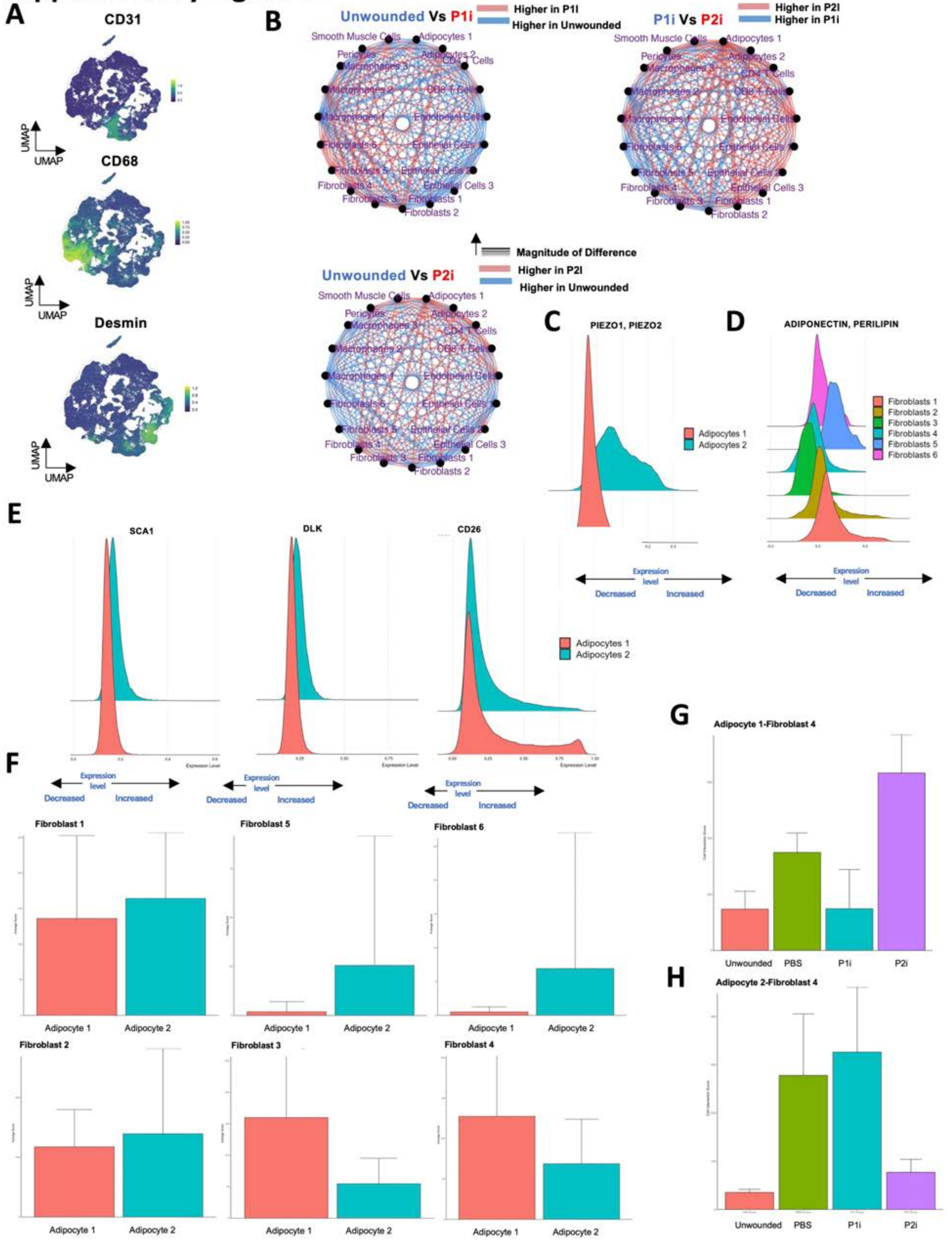

**Figure S21: CODEX analysis at POD 14. A.** Representative UMAP images of CD31 (top), CD68 (middle) and Desmin (bottom) of protein expression. **B.** Differential interaction maps in unwounded skin vs. P1i-treated wounds (top left), P1i-treated wounds vs. P2i treated wounds (top right), and unwounded vs P2i treated wounds (bottom). **C.** Histogram of protein expression of PIEZO 1 and PIEZO 2 in adipocytes. **D.** Histogram of protein expression of Adiponectin and Perilipin in fibroblasts. **E.** Histogram of protein expression of Sca1 (right), Dlk (middle), and CD26 (right) in adipocytes. **F**. Bar graph quantifying adipocyte-fibroblast intra interactions. **G**. Bar graph quantifying Adipocyte 1 - Fibroblast 4 interactions. **H**. Bar graph quantifying Adipocyte 2 - Fibroblast 4 interactions.

#### Supplementary Figure 22

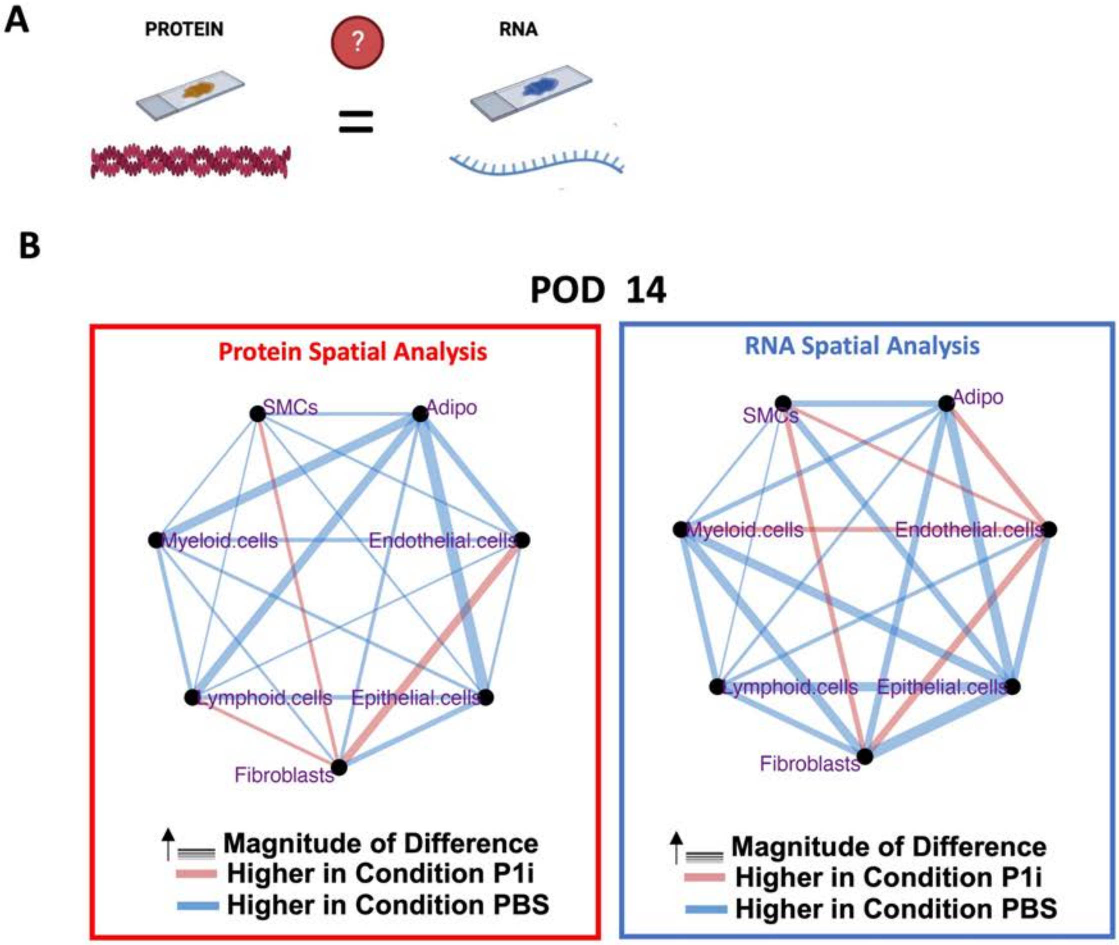

**Figure S22: Comparison of cell-cell interactions in CODEX and Visium analysis at POD 14.** **A.** Schematic to show comparison of spatial analysis at RNA and protein level. **B.** Differential interaction maps in PBS vs. P1i-treated wounds using CODEX analysis (left) and Visium analysis (right).

#### Supplementary Figure 23

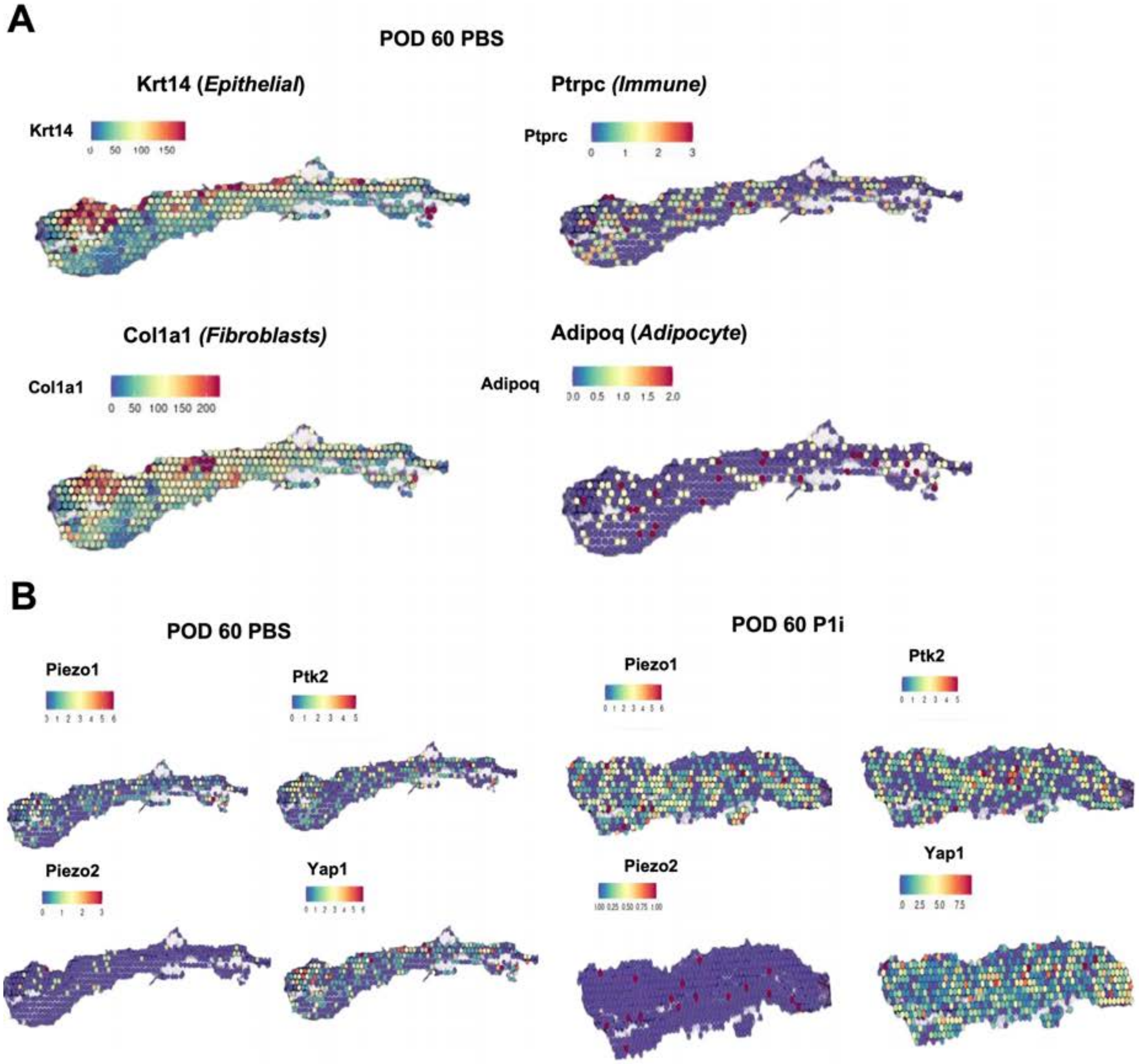

**Figure S23: VISIUM analysis at POD 14**. **A**. Spatial plots of epithelial (Krt14), immune (Ptprc), fibroblast (Col1a1), and adipocyte (Adipoq) markers at POD 60 scars following PBS treatment. **B**. Spatial plots of mechanical markers (Piezo1, Piezo2, Yap1, and Ptk2) at POD 14 following PBS treatment (left) and P1i treatment (right).

#### Supplementary Figure 24

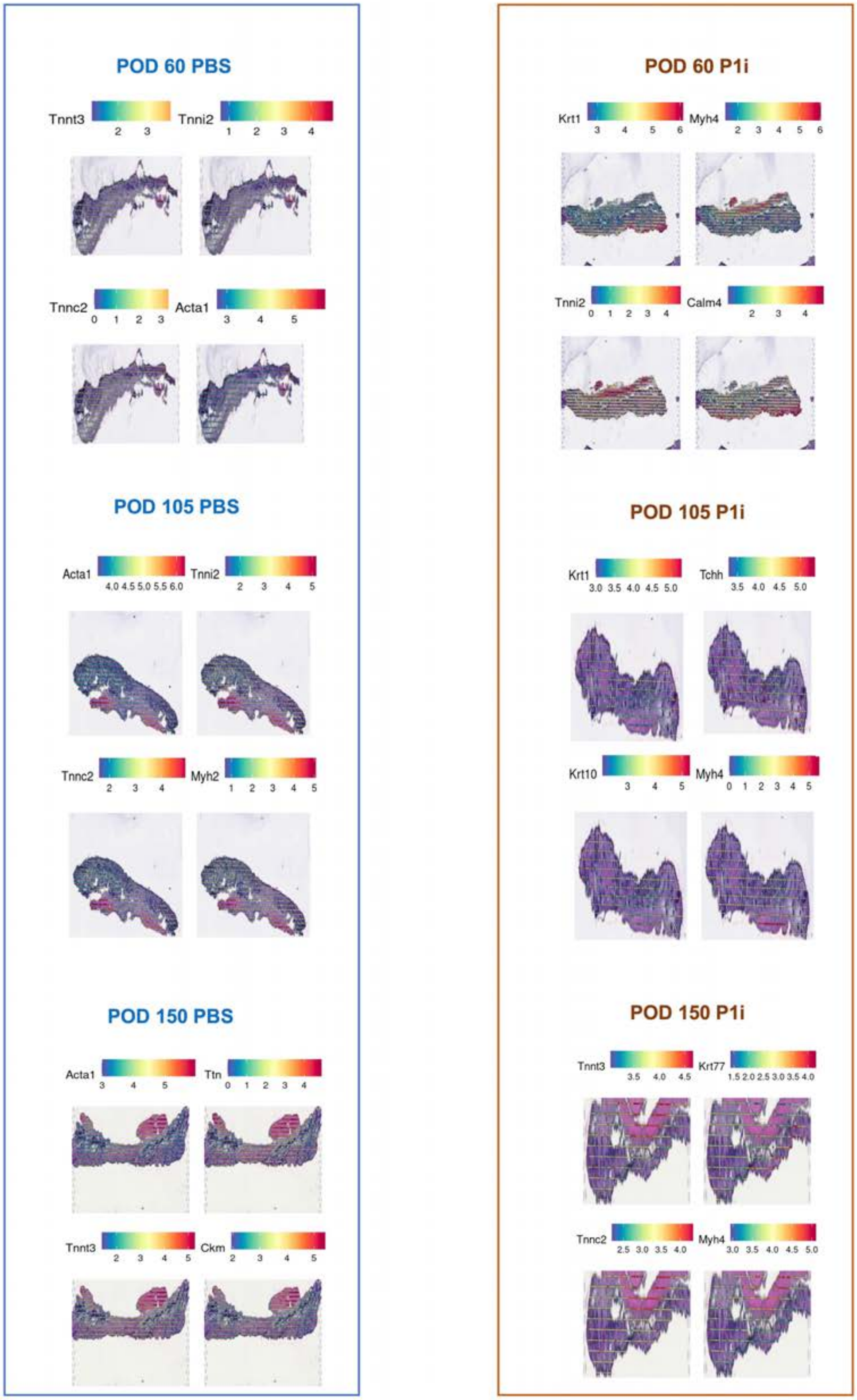

**Figure S24: VISIUM analysis at POD 60, 105, and 150**. Spatial plots of spatially variable features at POD 60 following PBS treatment (left) and P1i treatment (right).

#### Supplementary Figure 25

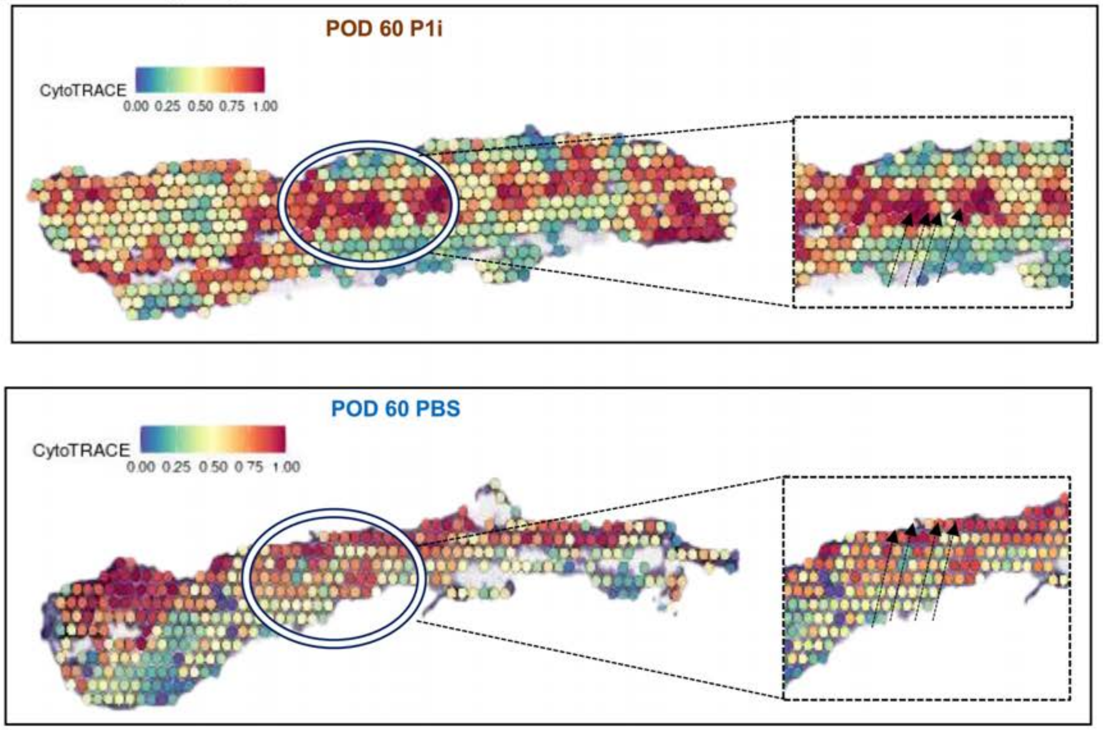

**Figure S25: CyoTRACE analysis of scars at POD 60**. Spatial plot of scars at POD 60 treated with P1i or PBS colored by CytoTRACE.

#### Supplementary Figure 26

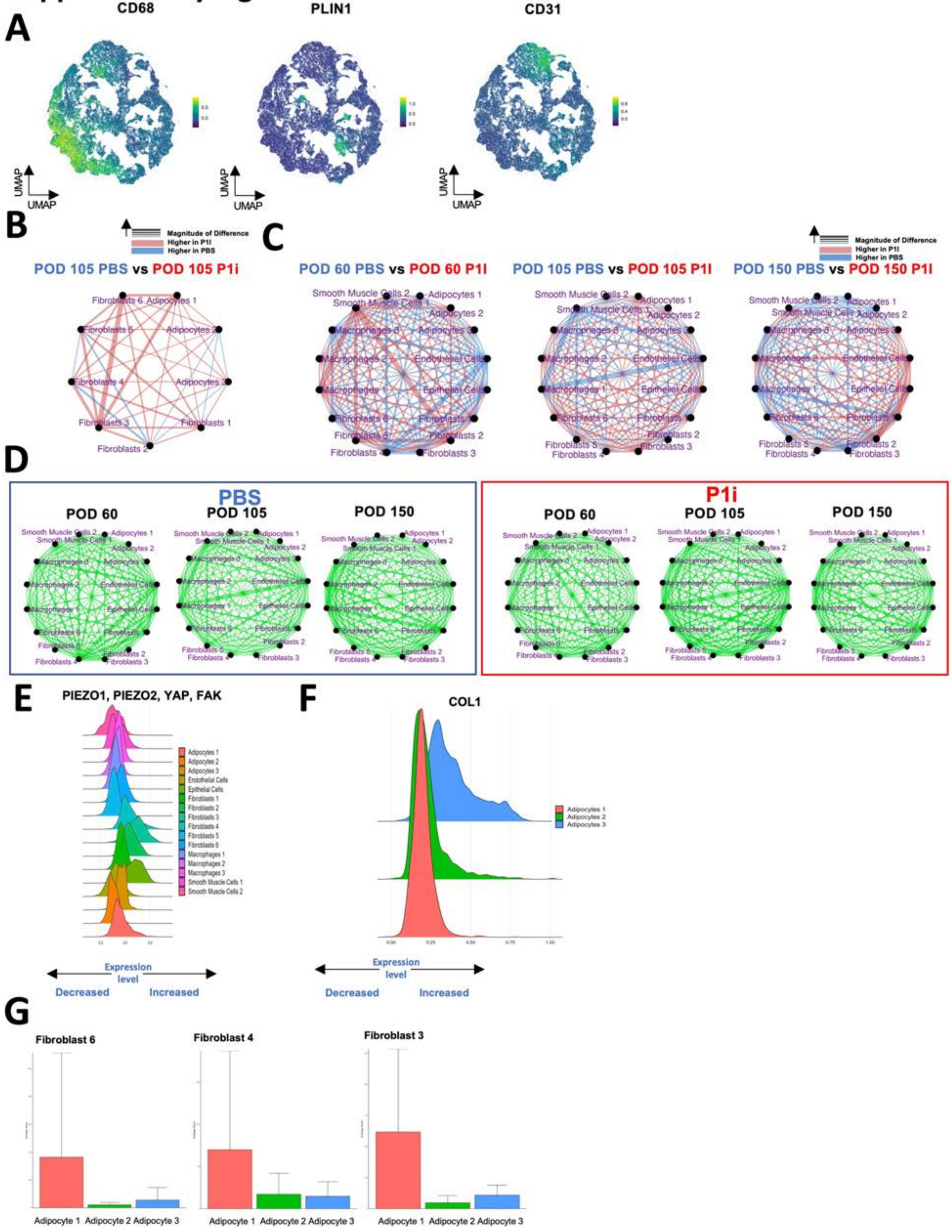

**Figure S26: CODEX analysis at POD 60, 105, and 150. A.** Representative UMAPs of CD68 (left), PLIN1 (middle), and CD31 protein expression (right). **B**. Differential interaction maps of adipocytes and fibroblasts in PBS-treated wounds vs. P1i-treated wounds at POD 105. **C.** Differential interaction maps in PBS-treated wounds vs. P1i-treated wounds (left) at POD 60, POD 105 (middle), and POD 150 (right). **D**. Differential interaction maps in PBS-treated wounds (left) at POD 60, 105 and 150 and in P1i treated wounds (right) at POD 60, 105 and 150. **E.** Histogram of protein expression of PIEZO 1, PIEZO2, YAP, and FAK in adipocytes. **F.** Histogram of protein expression of COL1 in adipocytes. **G** Bar graph quantifying Adipocyte-Fibroblast inter interactions.

### Supplementary Figure 27

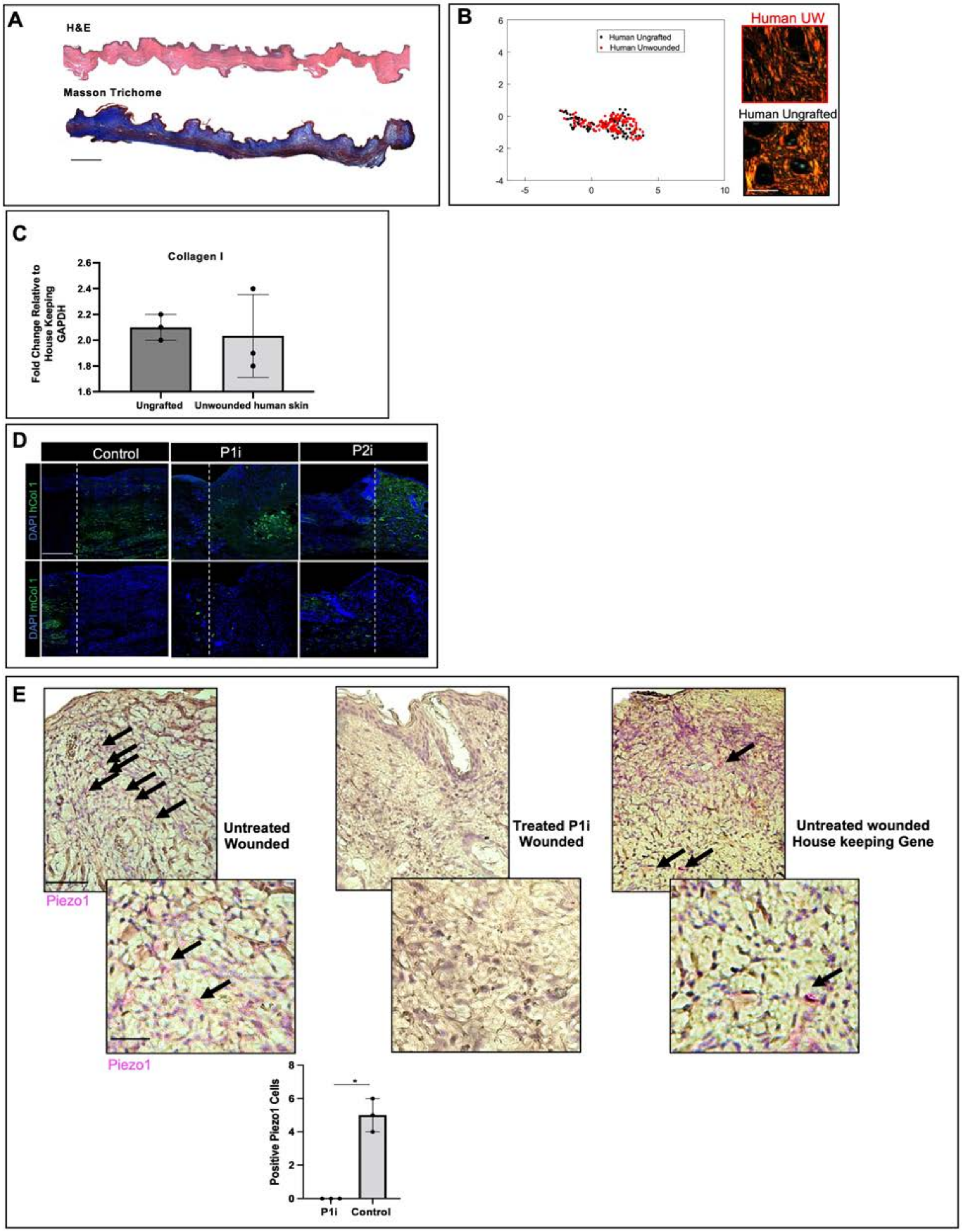

**Figure S27: Analysis of human foreskin tissue xenografts. A.** H&E (top) and Masson's trichrome (bottom) histology of ungrafted human foreskin samples. **B.** Left, UMAP of quantified extracellular matrix (ECM) ultrastructure parameters (each dot represents one histology image); right, picrosirius red histology of unwounded xenografted (UW) vs. native, ungrafted human foreskin. **C.** RT-qPCR quantification of Col1 expression by ungrafted versus unwounded grafted foreskin. **D.** Immunofluorescence (IF) staining for human-specific Col1 (hCol1, top) and mouse-specific Col 1 (mCol1, bottom; both, green signal) in consecutive sections of xenograft samples. White dotted lines indicate boundary between mouse skin (left side of each image) and xenografted human skin (right side). **E.** Top, RNAscope *in situ* hybridization of control (untreated) wounds (far left, far right) and P1i xenograft wounds (middle) targeting Piezo1 (left and middle panels) or positive control housekeeping gene (right panels; all, pink signal); black arrows indicate positively stained cells. Bottom, quantification of Piezo1$^{+}$ cells per HPF in each wound condition from RNAscope.

Data shown as mean ± S.D. *$P \leq 0.05$. Scale bars, 1 mm (**A**), 1 mm (**B**), 20 μm (**D**), 100 μm (**E**, first row), and 20 μm (**E**, second row).

#### Supplementary Figure 28

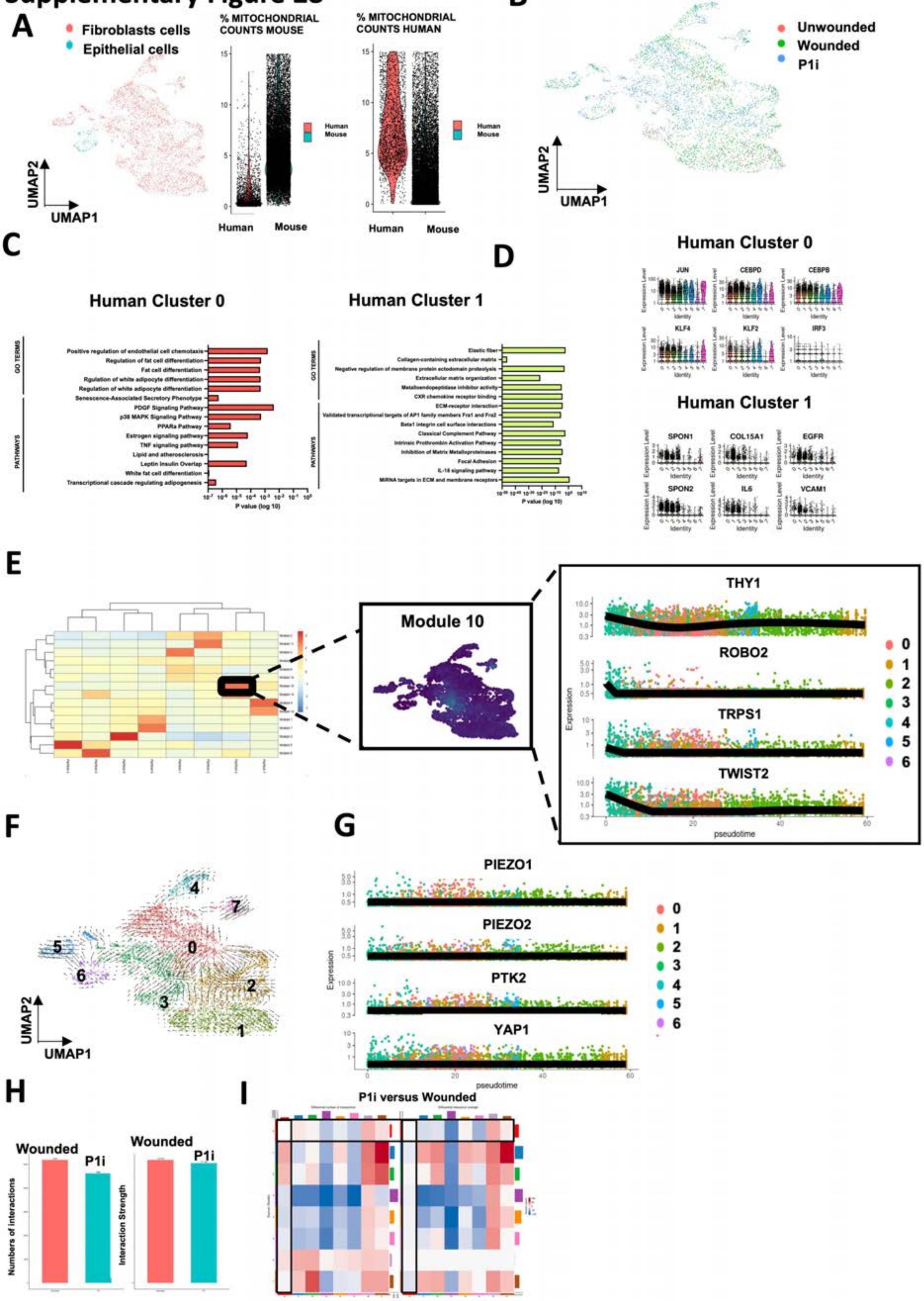

**Figure S28: scRNA-seq analysis of human xenograft at POD 14. A.** UMAP of all wound cells colored by cell type (left) and human/mouse origin (right). **B.** UMAP of all wound cells colored by wound treatment including unwounded, wounded, and P1i treated. **C**. Gene ontology pathways (GO) pathways of human cluster 0 (left) and human cluster 1 (right). **D**. Violin plots showing expression of indicated characteristic genes in cluster 0 (top) and cluster 1 (bottom). **E**. Expression of genes by pseudotime analysis with gene module 10 highlighted (indicated in black box) across pseudotime with selected genes including Thy1, Robo2, Trsp1, and Twist2. **F**. scVelo analysis of UMAP of all fibroblast cells. **G**. Expression of mechanical genes through pseudotime analysis coloured by Seurat cluster. **H**. Cell chat analysis of cell-cell interactions in wounded (red) and P1i-treated wounds (green) at POD 14. **I.** Heatmap of cell communications in P1i treated wounds versus untreated wounds at POD 14. in PBS and P1i treated wounds at POD 14 (black box showing interactions between human cluster 0 cells).

#### Supplementary Figure 29

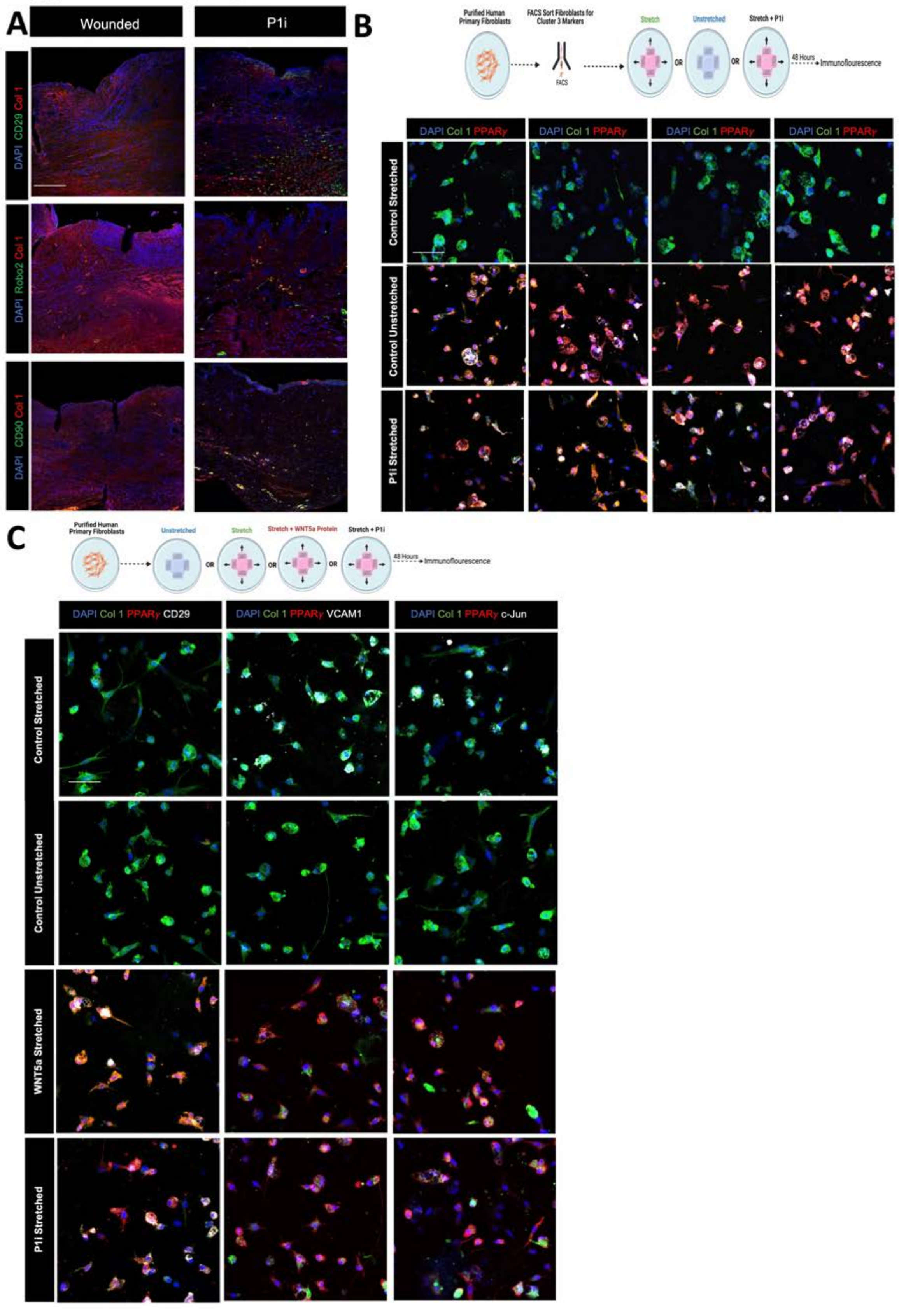

**Figure S29: Xenograft analysis following at POD 14. A.** Immunostaining of xenograft wounds for CD29, Robo2, and CD90 in wounded and P1i-treated wounds at POD 14. **B**. Top: Schematic showing experimental plan. Bottom: Immunostaining of human fibroblasts sorted by cluster 3 markers for PPary and Collagen type 1 following stretching (top row), no stretch (middle row), and stretching with P1i inhibitor (P1i) (bottom row). **C**. Top: Schematic showing experimental plan. Bottom: Immunostaining of human fibroblasts for CD29, VCAM1, CJUN following stretching (top row), no stretch (second row), stretching with Wnt5A recombinant protein (third row), and stretching with P1i (bottom row).

Data shown as mean ± S.D. $*P \leq 0.05$. Scale bars, 100 μm (**A**), 200 μm (**B**).

#### Supplementary Figure 30

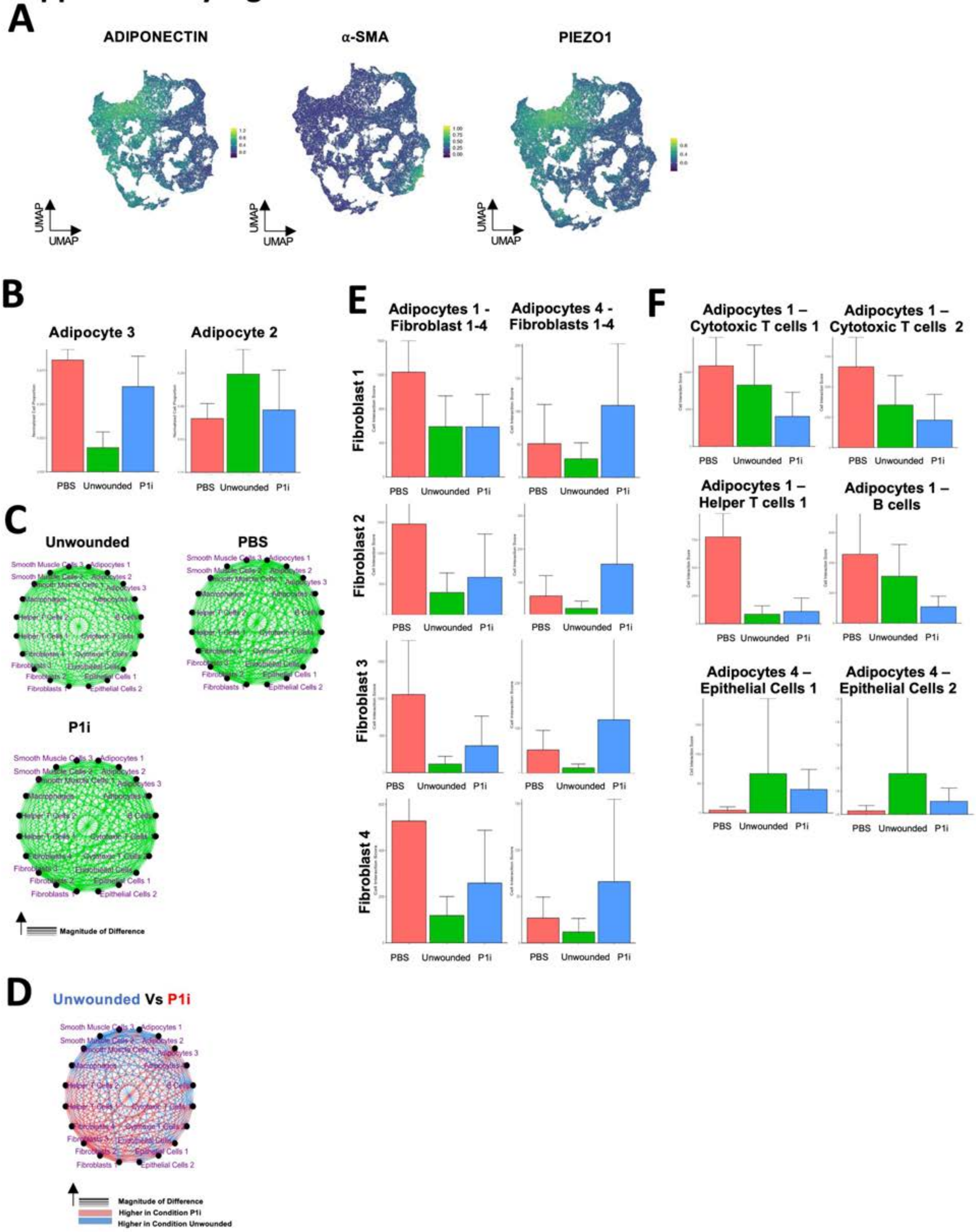

**Figure S30: CODEX analysis of human xenografts. A.** Representative UMAPs of Adiponectin (left), a-SMA (middle), and Piezo1 (right). **B**. Bar graphs showing cell proportion of adipocyte 3 and adipocyte 2 in PBS, P1i treated, and unwounded skin. **C**. Differential interaction maps in unwounded skin (top left), PBS treated skin (top right), and P1i treated skin (bottom). **D**. Differential interaction maps in unwounded vs. P1i-treated wounds (left) at POD 14. **E**. Bar graphs showing interactions of adipocyte and fibroblast cells in unwounded, PBS, and P1i treated wounds. **F**. Bar graphs showing interactions of adipocyte and other cell types in unwounded, PBS, and P1i treated wounds.

#### Supplementary Figure 31

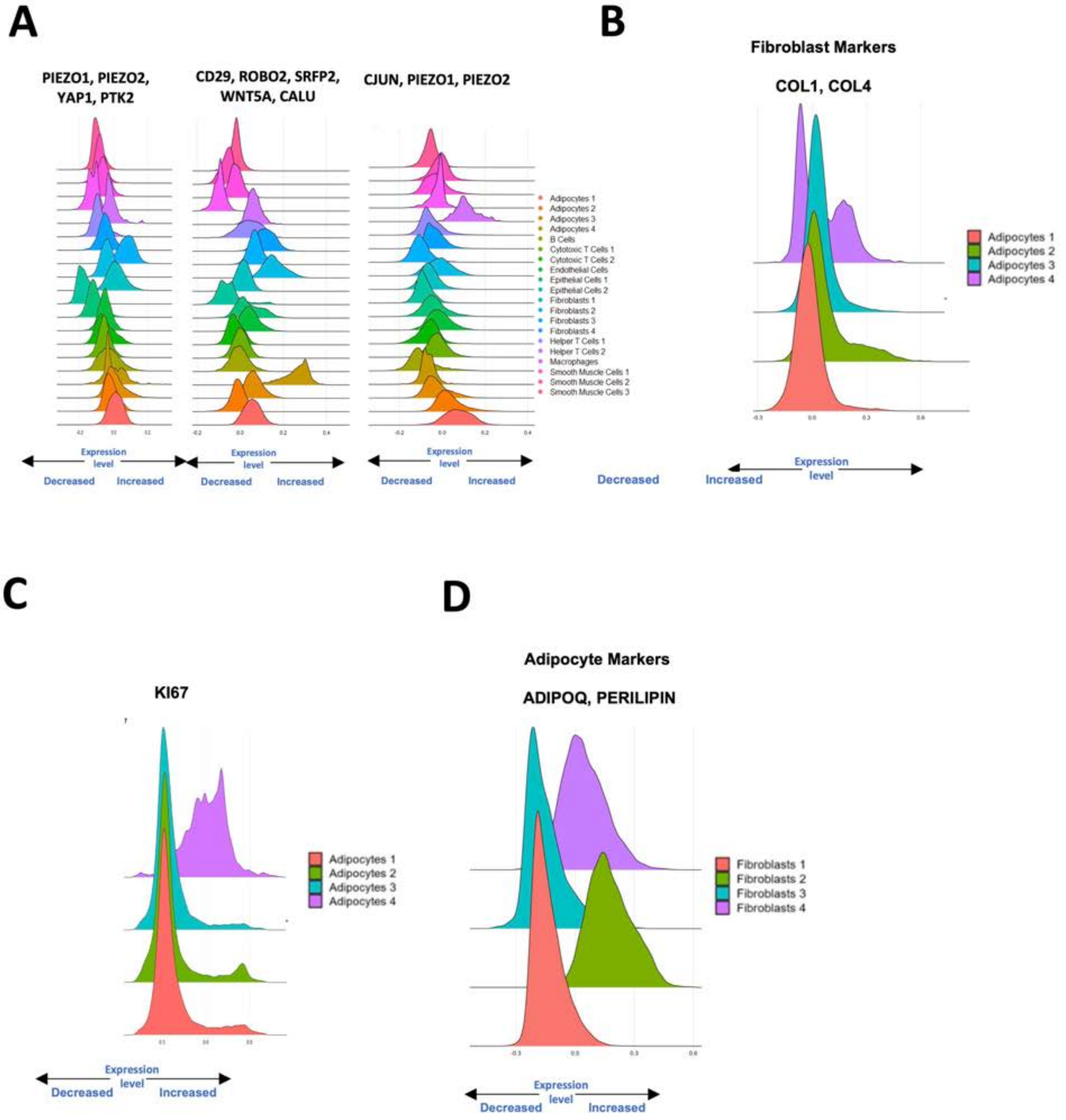

**Figure S31: CODEX analysis of human xenografts. A.** Histogram of co-expression of either $FAK^+PIEZO1^+PIEZO2^+YAP^+$ (left), $CD29^+ROBO2^+SFRP2^+WNT5A^+CALU^+$ (middle), and $CJUN^+PIEZO1^+PIEZO2^+$ (right) in all cell types. **B**. Histogram of $COL1^+$ $COL4^+$ in adipocyte clusters. **C**. Histogram of Ki67 in adipocyte clusters. **D**. Histogram of $ADIPOQ^+PLIN1^+$ in fibroblast clusters.

#### Supplementary Figure 32

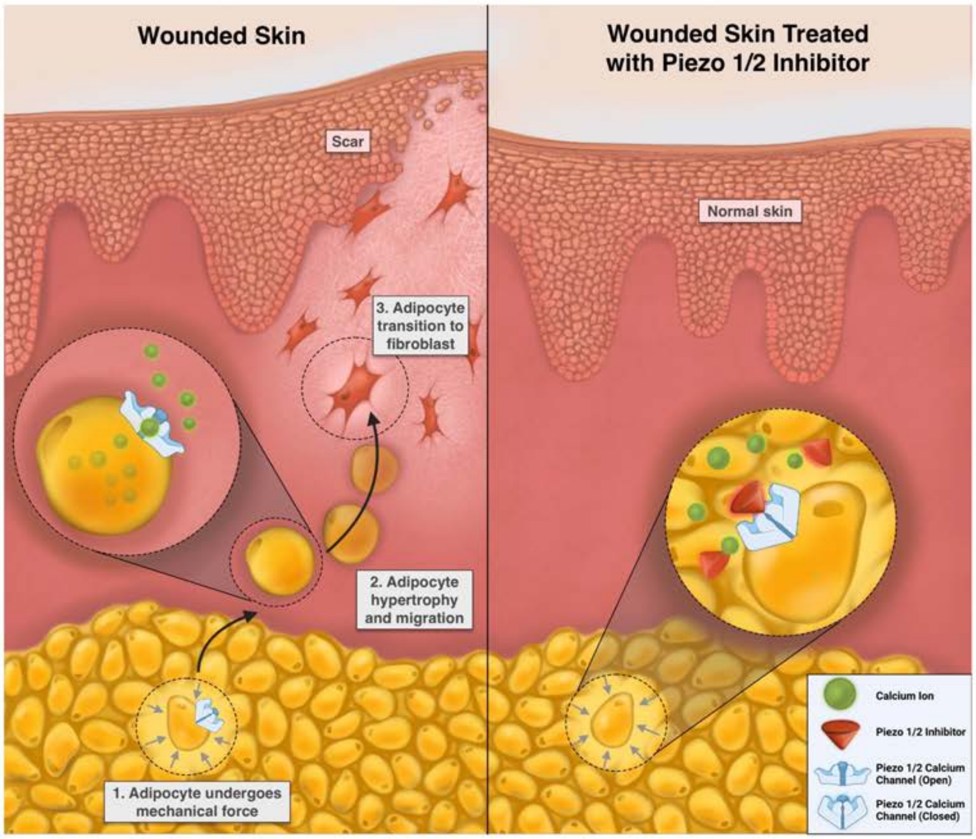

**Figure S32: Schematic illustrating the mechanism by which adipocytes convert to fibroblasts under mechanotransduction to cause scarring.**

#### Materials and Methods

##### Mice:

Transgenic mouse strains (acquired from Jackson Laboratories): **Adipoq-cre/ERT2** (B6;129-*Adipoqtm1Chan*/J, Stock: 08195), **Col 1** (B6.Cg-Tg(Col1a1-cre/ERT2)1Crm/J Stock: 016241), **mTmG** (B6.129(Cg)-*Gt(ROSA)26Sortm4(ACTB-tdTomato,-EGFP)Luo*/J Stock: 007676), **Piezo1**$^{\mathbf{fl/fl}}$ (*Piezo1tm2.1Apat*/J Stock: 029213)*,* **Piezo2**$^{\mathbf{fl/fl}}$ (B6(SJL)-*Piezo2tm2.2Apat*/J, Stock: 027720)*,* **Piezo2-EGFP-IRES-Cre** (B6(SJL)-*Piezo2tm1.1(cre)Apat*/J, Stock: 027719)*,* **Piezo1**$^{\mathbf{p1\text{-}tdT}}$ (B6;129-*Piezo1tm1.1Apat*/J, Stock: 029214)*,* **FAK**$^{\mathbf{fl/fl}}$ (B6.129P2(FVB)-*Ptk2tm1.1Guan*/J, Stock: 031956)*,* **YAP**$^{\mathbf{fl/fl}}$ (*Yap1tm1.1Dupa*/J*,* Stock: 027929), **ROSA26**$^{\mathbf{iDTR}}$ (*C57BL/6-Gt(ROSA)26Sortm1(HBEGF)Awai*/J*,* Stock: 007900), **CD1** (C.129S2-*Cd1tm1Gru*/J*,* Stock: 003814), **B6** (C57BL/6J, Stock: 000664). Mice were housed at the Stanford University Comparative Medicine Pavilion per Stanford APLAC guidelines, under the supervision of the Veterinary Service Center (VSC). All animals were genotyped under Jackson Laboratory instructions, using Transnetyx's Automated Genotyping PCR services. *ROSA26*$^{mTmG}$ mice utilize a dual-fluorescence reporter system that irreversibly substitutes Tomato red fluorescent protein (RFP) with membrane-bound green fluorescent protein (GFP) after recombination. *Adipoq*$^{Cre\text{-}ERT2}$ mice were crossed with *ROSA26*$^{mTmG}$ reporter mice to trace Adiponectin-lineage-positive cells, as defined by GFP expression. Rainbow (*ROSA26*$^{VT2/GK3}$) mice were gifted by the Weissman Laboratory, Stanford University School of Medicine.

##### Dorsal excisional wounding:

*Induction:*

Intraperitoneal tamoxifen injections (90% corn oil/ethanol v/v; 200 mg/kg body weight) were used to induce *Adipoq$^{Cre/ERT2}$;ROSA26$^{mTmG}$* mice (*n* = 6 mice per experimental group) every day for 5 consecutive days prior to surgery. Topical tamoxifen (150µL) was applied for 3 consecutive days following surgery.

*Wounding:*

For wounding, anesthesia was induced and maintained with 1-3% isoflurane at a flow rate of 2L/min. Adequate anesthesia was confirmed with the loss of hind-limb reflex to nociceptive stimuli. Dorsal skin was sterilized with Betadine Surgical Scrub Veterinary (Avrio Health L.P.™, Stamford, CT) followed by sterile alcohol prep pads (FisherScientific™, Pittsburgh, PA). Next, four 6 mm full-thickness circular wounds were made through the panniculus carnosus with sterile scissors and forceps. Wounds were equally spaced on the dorsum of each animal at least 4 mm lateral to the midline and stented open using 10 mm diameter silicone rings. Rings were secured using Krazyglue™ and 6 simple interrupted nylon monofilament 4-0 sutures (Dynarex™, Orangeburg, NY). Wounds were dressed using 3M Tegaderm Transparent Film 1626w dressings (3M™, Cat:1626W). Dressing changes took place every 48 hours under anesthesia. 200 ng topical dipheria toxin (DT) in 30 µL PBS (or 30 µL PBS control) was injected intradermally every day over 3 days to ablate adipocytes in *Adipoq$^{Cre\text{-}ERT2/Awai}$* mice. For mice receiving treatments with inhibitors of mechanosensitive ion channels, treatments consisted of a single administration at POD 0, via local intradermal injections into the wound edge; PBS was injected for vehicle controls. 30 µL (0.2mg/mL of inhibitor) was delivered per wound area of Piezo1 inhibitor (GsMTx4 [500nM], Tocris, Cat: 4912) or Piezo 2 inhibitor (D-GsMTx4 [500nM], Tocris, Cat: 4912) (n=6).

*Harvest:*

Wounds were re-epithelialized by postoperative day 14 (POD 14), at which time the wounds were harvested with dissecting scissors and processed for histology. Dissection followed fascial planes. Surrounding skin was also harvested for use as unwounded controls. Mice were euthanized by $CO_2$ narcosis and cervical dislocation. Harvested skin for Fluorescence-activated cell sorting (FACS) was mechanically digested using dissecting scissors to finely mince each specimen. Harvested skin for use in histology and immunofluorescent (IF) staining was placed in tissue embedding cassettes.

*Transplant:*

Fat pads were harvested from mTomato$^+$ (*mTmG*) mice and underwent mechanical digestion as above. Tissue was then enzymatically digested in a Collagenase II and IV solution (1:1, 1500U/ml in Dulbecco's Modified Eagle Medium [DMEM]). Samples were placed on an oscillating plate at 150 rpm for 40 minutes at 37°C. At the end of digestion, FACS buffer (see FACS section below for composition) was added to stop enzyme activity and solution was strained using a 70um cell strainer. Tomato$^+$ adipocytes were then injected intradermally into a wildtype (B6) mouse, which was prepared 24 hours prior to injection as described in the section above (*Dorsal Excisional Wounding)*. Then, 48 hours after injection, wildtype mice ($n$ = 4) were wounded in the engrafted area. Harvests took place at POD 14.

**Hypertrophic Scarring (HTS) Model:**

Mice were prepared as described in the section above (*Dorsal Excisional Wounding).* Linear incisions were made on the dorsum of the mice approximately 20 mm long. Incisions were closed using simple interrupted nylon monofilament 4-0 sutures (Dynarex™, Orangeburg, NY). A loading device consisting of 22 mm expansion screws and Luhr plate supports was placed over each wound on POD 4. The device was secured using Krazyglue™ and sutured in place. Tension was increased by 2mm expansion of the loading device every 2 days for 10 days. Mice with attached devices and no expansion protocol, as well as incision only with no device attached, also served as controls. Both wounded and control samples were harvested on POD 18 (n=6).

**Xenograft model:**

*Animals:*

For the xenograft model, adult immunocompromised CD-1 nude mice were purchased from The Jackson Laboratory (Bar Harbor, ME). Mice were maintained at the Stanford University Research Animal Facility, SIM-1 Barrier Facility in accordance with Stanford University guidelines. All experiments were performed under an approved APLAC protocol (APLAC #11048). 3 mice were used for each group.

*Human foreskin samples:*

Discarded postnatal human foreskin samples were collected after circumcision at the Lucille Packard Children's Hospital at Stanford under a Stanford University Institutional Review Board (IRB) approved protocol (IRB #45219). Samples were not used if collected >8 hours after circumcision; samples were kept in Dulbecco's Modified Eagle Medium (ThermoFisher Scientific™, Waltham, MA) on ice until grafting. Foreskin was prepared by removal of residual muscle and divided into 1cm sections to be grafted.

*Grafting protocol:*

Nude mice were induced and maintained with 2-4% isoflurane (Product:502017, MWI Veterinary Supply Co.®; Boise, ID) at a rate of 2L/min. While anesthetized, Betadine Surgical Scrub Veterinary (Avrio Health L.P.™, Stamford, CT) and sterile alcohol prep pads (FisherScientific™, Pittsburgh, PA) were used to prepare the dorsum. A 1.2 cm full-thickness excision of dorsal skin was created with forceps and sharp scissors. The foreskin graft was then placed into the full-thickness wound. 5-0 monofilament Nylon sutures (Medtronic, Minneapolis, MN) were circumferentially placed in a simple interrupted fashion and equal cross tension was maintained across the graft during placement. Tegaderm film dressings (3M™, Saint Paul, MN) served as pressure dressings post-operatively. Dressings were kept for five days post-operatively before being removed.

*Wounding protocol:*

At two weeks post-operatively, complete engraftment was established and the nylon sutures were removed. A 4mm punch biopsy was then used to create a full thickness wound within the graft using the described sterile technique. For mice receiving treatments with inhibitors of mechanosensitive ion channels, treatments were administered via local intradermal injections into the wound edge; PBS was injected for vehicle controls. 30 µL (0.2mg/mL of inhibitor) was delivered per wound area of Piezo1 and Piezo 2 inhibitors at POD 0. Wounds were dressed with Tegaderm film dressings (3M™, Saint Paul, MN) and changed every 48 hours for 5 days. Wounds were observed until POD 14 at which point they were harvested for analysis.

**Wound curve analysis:**

Gross measurements were taken using Adobe Photoshop 22.5.1 (Adobe Systems, San Jose, CA) to determine the percent closure of the wound until POD 14 (n= 6). Scar size was measured relative to the inner circumference of the silicone ring. Percentage of the original wound is plotted from baseline day 0 to day 16.

**EdU *in vivo*:**

Mice received intraperitoneal injections of EdU (5-ethynyl-2′-deoxyuridine) at a concentration of 100mg/kg 24 hours before wounding and again 24 hours before euthanasia (n = 6). For FACS analysis, cells were extracted using digestion methods previously described and stained using the Click-iT™ Plus EdU Alexa Fluor™ 647 Fluorescence-activated cell sorting Assay Kit (ThermoFisher, Cat:C10634). EdU incorporation was analyzed using a BD FACSAria II. For Immunohistochemical analysis of EdU incorporation in vivo, skin was harvested, fixed, and sectioned as previously described. Slides with skin sections were stained using Click-iT™ EdU Cell Proliferation Kit for Imaging, Alexa Fluor™ 647 dye (ThermoFisher, Cat:C10337), and fluorescent images of live cells were taken with an LSM880 inverted confocal microscope.

**Tensile strength testing:**

Skin from wounded and unwounded mice at day 14 were tested using an Instron 5565 using a 100 N load cell. Dorsal wounds were excised with sharp surgical scissors and carefully cut into tapered 4mm by 15mm pieces. Tissue pieces were subsequently anchored with grips such that the middle of the wound was positioned centrally between the grips. The tissue was slowly separated (1% increase per second) until failure (defined by a clear drop in measured stress as tension increased).

Young's Modulus was determined by taking the slope of the linear portion of the stress-strain curve.

**Harvesting cells for fluorescence-activated cell sorting (FACS):**

Following $CO_2$ euthanasia, dorsal skin wounds were dissected and washed once in PBS. Wounded tissue was then diced into a fine consistency using sharp surgical scissors. Minced skin tissue was then enzymatically digested using a 1:1 ratio of Collagenase Type IV (ThermoFisher, Cat:17104019) and Collagenase Type II (ThermoFisher, Cat:17101015) at a concentration of 1500U/ml in DMEM (ThermoFisher, Cat:10569010) for 90 minutes at 37°C. Samples were continuously agitated at 150rpm during enzymatic digestion. Enzyme activity was stopped using FACS buffer, and digested tissue was strained through 70um cell strainers. Cells were pelleted at 1500rpm for 5 minutes at 4C and then resuspended in 150uL of FACS buffer for primary antibody staining. For lineage negative (Lin-) FACS analysis, the following primary antibodies were used: Tie2 (ThermoFisher, Cat:13598782), CD45 (ThermoFisher, Cat:48045182 or ThermoFisher, Cat:13045181), CD31 (ThermoFisher, Cat:RM52280 or ThermoFisher 13031181), CD324 (ThermoFisher, Cat:13324982), CD326 (ThermoFisher, Cat:45592185), Ter119 (ThermoFisher, Cat:14592182). For fibroblast subpopulation analysis, the following primary antibodies were used: D1k1 (R&D systems, Cat:FAB8635A or FAB8634N), Sca1(Ly-6A/E) (BioLegend, Cat:108133), and CD26 (ThermoFisher, Cat:45026182 or Biolegend 137809). Cells were stained with primary for 30 minutes on ice. Following primary antibody staining, samples were washed with 500uL of FACS buffer, spun at 1500rpm for 5 minutes (4C), and resuspended in 150uL of FACS buffer for secondary antibody staining using either Streptavidin-eFluor450 (ThermoFisher, Cat: 48431782) or Streptavidin-Alexa Fluor 647 (S21374). Cells were stained with secondary for 20 minutes on ice. Cells were washed again with 500uL of FACS buffer, and DAPI (4′,6-diamidino-2-phenylindole) (Biolegend, Cat: 422801) was added to label dead cells. A BD II FACS Aria machine was used for FACS sorting and analysis.

**Histology and immunofluorescent staining:**

*Fixation:*

Fluorescent tissues from transgenic mice were fixed in 4% paraformaldehyde (PFA) solution in PBS (Electron Microscopy Sciences, Cat: 15710) for 24h at 4°C. All other samples were fixed in 10% neutral buffered formalin (NBF; ThermoFisher Scientific™, Waltham, MA) for 24 hours at room temperature.

*Cryosectioning:*

Fixed samples were soaked in 30% sucrose dissolved in PBS at 4°C. After one week, samples were removed from the sucrose solution and embedded as tissue blocks using Tissue Tek O.C.T. (Sakura Finetek, Torrance, CA) over dry ice and 100% ethanol to achieve rapid freezing. Frozen blocks were mounted on a Thermo Scientific CryoStar NX70 cryostat, and 8 µm-thick sections were transferred to Superfrost/Plus adhesive slides (ThermoFisher Scientific™, Waltham, MA).

*Paraffin sectioning:*

Automated Tissue Processor (ThermoFisher Scientific™, Waltham, MA) was used to dehydrate samples in a gradient of alcohols. Tissue was then embedded using ThermoFisher Histostar Tissue Embedding station. Paraffin blocks were trimmed as necessary and cut as 8 µm-thick sections.

Paraffin ribbons were placed in a water bath at 40°C and mounted onto Superfrost/Plus adhesive slides (ThermoFisher Scientific™, Waltham, MA). Sections were baked at 50°C overnight.

*Staining:*

Hematoxylin and eosin (Cat:H-3502; Vector Laboratories, Burlingame, California), Masson's Trichrome (ab150686; Abcam®, Waltham, MA), Picro-sirius Red (ab150681; Abcam®, Waltham, MA), and Oil Red O (Sigma-Aldrich™, St. Louis, MO) stains with standard protocols were used. Cryosection samples were first dehydrated using a slide rack, submerged into 1% PBS for 10 minutes, followed by 30% ethanol (EtOH), 50% EtOH, 70% EtOH, 95% EtOH, and 100% EtOH for 15 minutes each. Paraffin sections were hydrated prior to staining by placement in xylene for 20 minutes, followed by 10 minutes each of 100% EtOH, 95% EtOH, 70% EtOH, 50% EtOH, and 30% EtOH. Slides were then submerged in running tap water for 10 minutes.

For immunofluorescent staining, slides were washed twice in Tween 20 (Sigma-Aldrich™, St. Louis, MO) followed by one wash in PBS. Slides were then blocked for 1 hour with Power Block (Biogenex™, Fremont, CA) prior to addition of the following primary antibodies: Abcam ab40794 (anti-FAK), Abcam ab3526 (anti-perilipin), Abcam ab5694 (anti-α-SMA), Abcam ab181595 (anti-CK14), Abcam ab52625 (anti-CK19), Abcam ab59436 (anti-collagen type III), Abcam ab51317 (anti-Sca1), Abcam ab197896 (anti-S100A4), ThermoFisher Scientific MA1-26771 (anti-collagen type I), ThermoFisher Scientific PA5-16571(anti-PDGFRα), ThermoFisher Scientific PA3-821A (anti-PPARγ), R&D systems anti-mAcrp30 (anti-adiponectin), Proteintech 15939-1-AP (anti-Piezo1), Novus Biologicals NBP1-78624 (anti-Piezo2), Santa Cruz Biotechnology SC-101199 (anti-YAP1), Santa Cruz Biotechnology SC-7309 (anti-CD36), Sigma-Aldrich AB9260 (anti-Ki-67).

Slides were then incubated for 1 h with Alexa Fluor 488, 594, or 647-conjugated anti-rabbit, anti-rat, or anti-mouse antibodies (Invitrogen, Waltham, MA). Finally, slides were mounted in Fluoromount-G mounting solution with or without DAPI (ThermoFisher Scientific™, Waltham, MA). Brightfield images were acquired with a Leica CTR4000 microscope, while fluorescent images were acquired with a LSM880 inverted confocal, Airyscan, AiryscanFAST, GaAsP detector upright confocal microscope.

Hematoxylin and eosin staining started with submerging slides into Hematoxylin for 10 minutes. Slides were then submerged into tap water for 5 minutes, then dipped into eosin 12 times. DI water baths were prepared and slides were submerged until water was clear of eosin. Slide racks were then dipped into 70% EtOH 10 times, followed by 1 minute in 95% EtOH then 100% EtOH. Finally slides were dipped into Xylene 8 times until they were mounted on Superfrost/Plus adhesive slides (ThermoFisher Scientific™, Waltham, MA) with Permount Mounting Medium (Electron Microscopy Sciences™, Hatfield, PA).

For Trichrome staining, Bouin's solution was added to the samples for 60 minutes in a humidity chamber. To create humidity chambers, slide boxes were lined with wet paper towels. Next, slides were submerged into running tap water for 5 minutes. Working Weigert's Iron Hematoxylin was then added to samples for 5 minutes, followed by submersion into running tap water for 5 minutes. Biebrich Scarlet /Acid Fuchsin Solution was then added to samples for 4 minutes. After slides were submerged in running tap water for 5 minutes, Phosphomolybdic/Phosphotungstic Acid was added to the sample for 45 minutes. Phosphomolybdic/Phosphotungstic Acid was removed from slides and Aniline Blue Solution was added for 4 minutes without a washing step. After Aniline Blue staining, slides were submerged into running tap water for 5 minutes. Correct staining was confirmed using Leica CTR4000 microscope. Finally, slides were dipped into 1% Glacial Acetic Acid Solution then running water 12 times each, followed by 15 dips each in 95% EtOH then 100% EtOH. Slides were submerged into Xylene 8 times prior to mounting with Permount.

Dehydration/rehydration steps were not completed for Picrosirius red staining. Slides were first washed 3 times with PBS and then submerged into running tap water for 1 minute. Picrosirius red was added to slides for 60 minutes in a humidity chamber, previously described. After the completion of the 1 hour stain, slides were dipped into 2 different changes of 0.5% Glacial Acetic Acid 10 times (20 dips total), followed by 10 times into 2 different changes of 100% ETOH (20 dips total), and 8 times into Xylene. Slides were then mounted with Permount.

*Picrosirius red stained histologic analysis:*

Analysis of picrosirius red stained tissue sections took place using an image-processing algorithm. The algorithm profiles 26 ultrastructural features to provide a quantitative comparison of extracellular matrices.[11] Each group ($n$ = 6) was randomly imaged at 100 separate locations at 40x. Color deconvolution following previously described methods(Ruifrok and Johnston, 2001) was performed to characterize each stain by absorbance in three RGB channels. Ortho-normal transformation was then used to determine each color's contribution to the captured image. Red and green images were produced, representing mature and immature ECM fibers, and analyzed as separate groups. A Matlab script was used to achieve analysis, including noise reduction, preferential selection for smooth regions with low variance, and "skeletonization" of images to characterize the fiber networks. The algorithm also allowed for measurement of fiber length, width, persistence, alignment, and overall dimensionality.

Oil Red O Staining was completed using frozen sections. Sections were first fixed in formalin for 15 minutes, followed by a wash step in running tap water for 5 minutes. Slides were then rinsed in 60% isopropanol prior to a 15-minute staging in Oil Red O working solution. Working solution was made from 30 ml of the stock stain and 20 ml of distilled water. Stock stain was made using 0.5g of Oil Red O (Sigma-Aldrich™, St. Louis, MO) dissolved in 100 ml of isopropanol using a warm water bath. Filter paper was then used to add stock stain to a 50 ml conical. Stains were completed on the same day that a working solution was made using Coplin Staining Jars.

**RNAscope:**

RNAscope® 2.5 HD Assay in situ hybridization assay was conducted starting with a permeabilization step. Paraffin-embedded tissue sections were pretreated to permeabilize cells and expose target RNA. Next, RNAscope® and double-Z design probes were hybridized to the target sequence. Signals were amplified using detection reagents and labeled fluorescent probes were added to bind to each amplifier region. Target probes, amplifiers and label probes were sourced from Advanced Cell Diagnostics, Hayward, CA. Samples were imaged using a bright field microscope with a Leica CTR4000 microscope and positive pixels were quantified in Imagej.

**Cell culture:**

*Mouse cell line:*

Embryonic fibroblast cell line 3T3-L1 (CL-173, ATCC) was purchased and revived per manufacturer instructions. Following revival, cell expansion was monitored, and media was replaced every other day. Cells were passaged once confluency exceeded 70%. Cells used for experiments were between passages 2-5. Fibroblasts were cultured in DMEM + Glutamax media (ThermoFisher, Cat: 10569010) enriched with 10% fetal bovine serum (ThermoFisher, Cat: 10082147) and 1% Antibiotic-Antimycotic (ThermoFisher, Cat:15240062) at 37°C and 5% $CO_2$.

*Human primary adipocytes:*

Tissue was collected according to approved Stanford University IRB protocols. Human Lipoaspirate samples were washed in PBS and minced using sterile scissors. Samples were then enzymatically digested using Collagenase Type I (ThermoFisher, Cat: 17018029) at a concentration of 1500U/ml in DMEM (ThermoFisher, Cat: 10569010) for 45 minutes at 37°C. Samples were continuously agitated at 150rpm during enzymatic digestion. Enzyme activity was stopped using fetal bovine serum (FBS) enriched media, and digested tissue was strained through 300um and 100um cell strainers successively. Filtered samples were then spun at 1500rpm for 5 minutes at 4°C. Mature adipocytes were isolated from the supernatant.

**Differentiation of embryonic fibroblast cells to adipocytes:**

Embryonic fibroblast cell line 3T3-L1 (CL-173, ATCC) was treated with Adipogenic medium containing DMEM + Glutamax media (ThermoFisher, Cat: 10569010) enriched with 10% fetal bovine serum (ThermoFisher, Cat: 10082147), 1% Antibiotic-Antimycotic (ThermoFisher Cat:15240062), 10ng/mL Insulin (Sigma, Cat:0516), 500mM 3-isobutyl-1-methylxanthine (FisherScientific, Cat:AC228420010), 1mM Dexamethasone (Sigma, Cat:1756), and 1mM Rosiglitazone (Stem Cell Technologies, Cat:72622). Cells were incubated at 37°C and 5% $CO_2$. Cells were treated for 10 days and media was replaced every other day. Adipogenic differentiation protocol was based on published protocol(Guasti et al., 2012) and was validated in this study via

ICC. In order to validate the differentiation protocol, embryonic fibroblast cells (3T3-L1, ATCC) were seeded in 24 well plates at 20,000 cells per well. Once confluent, wells received either adipogenic differentiation media or standard fibroblast media for 10 days. Cells were then fixed (4% PFA for 24 hours at 4ºC) immunocytochemically stained with anti-Adiponectin (Novus Biologicals, Cat: AF1119) and anti-Collagen I (ThermoFisher Cat: MA1-26771) antibodies. Fluorescent images were taken with a LSM880 inverted confocal microscope.

**Proliferation and apoptosis:**

P1i and P2i treated adipocytes (and fibroblast controls) were examined for treatment toxicity using proliferation and apoptosis assays. Embryonic fibroblast cells (3T3-L1, ATCC) were seeded in 24 well plates at 20,000 cells per well. Once confluent, wells received either adipogenic differentiation media or standard fibroblast media for 10 days. Then, cells were treated with either 0.5x, 1x, or 2x the standard concentration of Piezo1 or Piezo2 inhibitors [500nm]. After 7 days of treatment, cells were fixed or harvested for analysis of potential toxicity. For proliferation, EdU (5-ethynyl-2′-deoxyuridine) incorporation in vitro was investigated on differentiated adipocytes and undifferentiated fibroblasts using the Click-iT™ EdU Cell Proliferation Kit for Imaging, Alexa Fluor™ 488 dye (ThermoFisher, Cat:C10337). Proliferation was detected according to the manufacturer's guidelines. To evaluate apoptosis, the Annexin V Apoptosis Detection Kit-AF647 (Abcam, Cat:219919) was used according to the manufacturer's guidelines.

**Gels:**

*Human Gels:*

Human adipocytes (collected and processed according to Cell Culture-*Human Primary Adipocytes*) were used to generate adipocyte-populated collagen I hydrogels as per a previously published protocol.(Chen et al., 2018) In short, a solution of 2mg/mL collagen I (Advanced Biomatrix, Cat:5005), 0.8x MEM (ThermoFisher, CAT:11430030) in 16mM HEPES (Sigma, CAT:83264), and suspended human adipocytes (concentration 250,000 cells/mL) in DMEM (ThermoFisher, Cat:10569010) were pipetted into cruciform-shaped PDMS (DOW, Cat:2646340) molds and allowed to gelate before removing the mold. Hydrogels were constrained by metal pins and fibroblast media containing DMEM + Glutamax media (ThermoFisher, Cat: 10569010) enriched with 10% fetal bovine serum (ThermoFisher, Cat:10082147) and 1% Antibiotic-Antimycotic (ThermoFisher Cat:15240062) was added. After 24 hours, formed hydrogels were subjected to either 10% equibiaxial strain or no strain. Strain was achieved through movement of metal anchor pins in each of the four arms of the hydrogel. Strain quantification was confirmed by Fiji analysis of $TiO_2$ dye marks on the central region of each hydrogel. Media containing small molecule mechanotransduction inhibitors to Piezo1 (GsMTx4 [500nM], Tocris, Cat: 4912) Piezo2 (D-GsMTx4 [500nM], Tocris, Cat: 4912), FAK (PF573228 [10uM], Tocris, Cat: 3239) YAP (Verteporfin [300nM], Sigma, Cat: SML0534), or Vehicle Control (PBS) was added. Media was replaced every other day. Hydrogels were kept for 7 days. For IHC analysis of gel contents, gels were fixed whole in 4% PFA (ChemCruz, Cat:281692) after being washed in PBS. For FACS and PCR analysis, cells were harvested by enzymatic digestion using Collagenase Type I (ThermoFisher, Cat:17018029) at a concentration of 1500U/ml in DMEM (ThermoFisher, Cat:10569010) for 25 minutes at 37°C. Enzyme activity was stopped using FBS enriched media. Cells were spun, and the pellet was processed for FACS or resuspended in TRIzol for PCR.

*Mouse gels:*

Mouse fibroblasts (collected according to Cell Culture-*Mouse Cell line*) were used to generate adipocyte-populated collagen I hydrogels as per a previously published protocol.(Chen et al., 2018) In short, a solution of 2mg/mL collagen I (Advanced Biomatrix, Cat:5005), 0.8x MEM (ThermoFisher, Cat:11430030) in 16mM HEPES (Sigma, CAT: 83264), and suspended mouse adipocytes (concentration 500,000 cells/mL) in DMEM (ThermoFisher, Cat: 10569010) were pipetted into cruciform-shaped PDMS (DOW, Cat:2646340) molds and allowed to gelate before removing the mold. Hydrogels were constrained by metal pins and Adipogenic medium containing DMEM + Glutamax media (ThermoFisher, Cat:10569010) enriched with 10% fetal bovine serum (ThermoFisher, Cat:10082147), 1% Antibiotic-Antimycotic (ThermoFisher Cat:15240062), 10ng/mL Insulin (Sigma, Cat:0516), 500mM 3-isobutyl-1-methylxanthine (FisherScientific, Cat:AC228420010), 1mM Dexamethasone (Sigma, Cat:1756), and 1mM Rosiglitazone (Stem Cell Technologies, Cat:72622) was added. After 10 days of differentiation, formed hydrogels were subjected to either 10% equibaxial strain or no strain. Strain was achieved through movement of metal anchor pins in each of the four arms of the hydrogel. Strain quantification was confirmed by Fiji analysis of $TiO_2$ dye marks on the central region of each hydrogel. Media containing small molecule mechanotransduction inhibitors to Piezo1 (GsMTx4 [500nM], Tocris, Cat: 4912) Piezo2 (D-GsMTx4 [500nM], Tocris, Cat: 4912), FAK (PF573228 [10uM], Tocris, Cat: 3239) YAP (Verteporfin [300nM], Sigma, Cat: SML0534), or Vehicle Control (PBS) was added. Media was replaced every other day. Hydrogels were kept for 7 days. For ICC analysis of gel contents, gels were fixed whole in 4%PFA (ChemCruz, Cat:281692) after being washed in PBS. For FACS and PCR analysis, cells were harvested by enzymatic digestion using Collagenase Type I (ThermoFisher, Cat: 17018029) at a concentration of 1500U/ml in DMEM (ThermoFisher,

Cat:10569010) for 25 minutes at 37°C. Enzyme activity was stopped using FBS enriched media. Cells were spun, and the pellet was processed for FACS or resuspended in TRIzol for PCR.

*Immunocytochemistry processing for gels:*

Following 24 hour fixation, gels were carefully dissected into 2mm x 2mm pieces and placed into wells of a 96 well plate. Gentle agitation was provided to the plates during staining via a platform shaker at 25°C. Gel pieces were washed in PBS for 30 minutes. Gels were then permeated with 0.1% TritonX-100 in PBS (ThermoFisher, Cat:85111) for 30 minutes. Gels were then blocked with PowerBlock (BioGenex, Cat:HK085-GP) for 1 hour. Following blocking, primary antibody was made up in a 1:1 solution of 0.1% TritonX-100 and PowerBlock. Primary staining lasted 1 hour, followed by two 15 minute washes in 0.025% Tween 20 in PBS (Sigma, Cat:P1379) and a single 15-minute wash in PBS. Secondary antibody containing Alexa Fluor 488, 594, or 647 against the primary antibody was added to a 1:1 solution of 0.1% TritonX-100 and Powerblock. Secondary antibody staining lasted 45 minutes, followed by two 30 minute washes in 0.025% Tween 20 in PBS (Sigma, Cat:P1379) and a single 30-minute wash in PBS. Finally, gel pieces were mounted on a slide with Fluoromount-G mounting solution with DAPI (ThermoFisher Cat:00495952). Fluorescent images were taken with an LSM880 inverted confocal microscope.

The following primary antibodies were used for staining the gels: Abcam ab40794 (anti-FAK), Abcam ab3526 (anti-perilipin), ThermoFisher Scientific MA1-26771 (anti-collagen type I), R&D systems anti-mAcrp30 (anti-adiponectin), Proteintech 15939-1-AP (anti-Piezo1), Novus Biologicals NBP1-78624 (anti-Piezo2), Santa Cruz Biotechnology SC-101199 (anti-YAP1), Santa Cruz Biotechnology SC-7309 (anti-CD36).

*Calcium imaging of gels:*

Calcium imaging was used to determine in vitro $Ca^{2+}$ flow in hydrogel-bound adipocytes with or without mechanical strain and with or without Piezo1 inhibitor treatment. A Fluo-4, AM cell permeant kit (ThermoFisher, Cat: F-14201) was used per the manufacturer's instructions. Fluorescent images of live cells were taken with an LSM880 inverted confocal microscope. Imagej was used for image quantification.

*shRNA knockdown of Piezo1 in gels:*

Short hairpin RNA (shRNA) knockdown of Piezo1 gene expression in human primary adipocytes was performed per manufacturer guidelines (Santa Cruz, US). Human primary adipocytes were collected as described in Cell Culture-*Human Primary Adipocytes* and populated into collagen 1 hydrogels as described in Gels-*Human Gels.* Twenty-four hours after the formation of the gels, shRNA knockdown, scrambled control, or vehicle control was delivered. Following knockdown, gels were subjected to 0 or 10% equibaxial strain. Seven days after strain, gels were harvested for FACS analysis or fixed in 4% PFA for Immunocytochemistry. Other than replacing traditional culture conditions in flasks/wells with a hydrogel populated system, no other modifications were made to the manufacturer's protocol.

*ELISA:*

Media samples from mouse adipocyte-populated collagen 1 hydrogels, human adipocyte-populated collagen 1 hydrogels, Mouse Adiponectin, Mouse Collagen 1, Human Collagen 1, and Human Adiponectin were tested using Abcam mouse/human enzyme linked immunoassay (ELISA) kits (Abcam, Cat: ab108785, ab210579, ab229389, USA) according to manufacturer's protocol. Standards and samples were placed into wells coated with Collagen 1 or Adiponectin specific antibodies. Enzyme-linked polyclonal antibodies were then added to wells, followed by the addition of substrate. A spectrophotometer captured absorbances, and concentrations of protein target were calculated and modeled using Prism.

**RT-qPCR:**

RNeasy mini kit (Qiagen LLC, Germantown MD) was used for RNA extraction for real time quantitative polymerase chain reaction (qPCR). Transcription was then performed using Moloney murine leukemia virus reverse transcriptase. Following standard manufacturers' protocols, an ABI Prism PCR7500 sequence detection system (Applied Biosystems, Waltham, MA) using TaqMan expression assays (ThermoFisher CA).

Primer sequences were used as follows:

| **Mouse** | **Human** |
|---|---|
| Adipoq :Mm00456425_m1 | ADIPOQ:Hs00605917_m1 |
| Col1a1: Mm00801666_g1 | COL1A1: Hs00164004_m1 |
| Gapdh: Mm99999915_g1 | GAPDH: Hs02786624_g1 |

**Single-cell sequencing:**

Dermal wounds were harvested at POD 14 and mechanically digested with sharp dissecting scissors (n = 6 per condition, performed in 3 independent experiments). Tissue was then added to an enzymatic digestion consisting of Collagenase II (ThermoFisher, Cat:17101015) and IV (ThermoFisher, Cat:17104019) in DMEM-F12 (GIBCOTM, Fischer Scientific, Hampton, NH).

Samples were added to an orbital shaker at 150 rpm for 90 minutes at 37ºC. FACS buffer was added to quench the digest and samples were filtered through 70 µm cell strainers. Samples were then centrifuged at 1500 for 5 minutes at 4°C. Cell suspensions were labeled with TotalSeq Series B hashtag oligonucleotide-labeled antibodies (BioLegend). Samples were then centrifuged and resuspended with 0.04% UltraPure BSA (Thermo Fisher, Waltham, MA); cell counts were completed. A second filtration step took place using 40 µm cell strainers. Quality control and single cell RNAseq were performed on unsorted cells using the 10x Chromium Single Cell platform (Single Cell 3' v3, 10x Genomics, USA) at the Stanford Functional Genomics Facility (SFGF), Stanford University, Palo Alto.

Base calls were converted to reads using the Cell Ranger (10X Genomics; version 3.1) implementation mkfastq and then aligned against the Cell Ranger mm10 reference genome, available at: http://cf.10xgenomics.com/supp/cell-exp/, using Cell Ranger's count function with SC3Pv3 chemistry and 5,000 expected cells per sample, as previously described. Hashtag oligos (HTOs) for samples were demultiplexed using Seurat's implementation HTODemux as previously described. For both datasets (Figure 2 and Figure 5), a maximum percent mitochondrial RNA cutoff of 15% was employed.(Mascharak et al., 2022) For Figure 2, cutoffs of 7,500 maximum unique genes and 85,000 maximum RNA counts were used. For Figure 5, cutoffs of 6,500 maximum unique genes and 65,000 maximum RNA counts were used. This resulted in 4,116 cells for Figure 2 and 45,725 cells for Figure 5 (12,041 of which were later classified as Fibroblasts).

Unique molecular identifiers (UMIs) from each cell barcode were retained for all downstream analysis, normalized with a scale factor of 10,000 UMIs per cell, and subsequently natural log transformed with a pseudocount of 1 using the R package Seurat (version 4.0.5).(Chen et al., 2013) The first 15 principal components of the aggregated data were then used for uniform manifold approximation and projection (UMAP) analysis.(Mascharak et al., 2022) Cell annotations were ascribed using SingleR (version 3.11) against the Mouse-RNAseq reference dataset, available at https://rdrr.io/github/dviraran/SingleR/man/mouse.rnaseq.html. Cell-type marker lists were generated using Seurat's native *FindMarkers* function with a log fold change threshold of 0.25 using the ROC test to assign predictive power to each gene. The 200 most highly ranked genes from this analysis for each cluster were used to perform gene set enrichment analysis in a programmatic fashion using EnrichR (version 2.1).(Chen et al., 2013)

The scRNA sequencing data generated during this study has been sent to the NCBI's Gene expression Omnibus for deposition and will be accessible on request.

**Pseudotime analysis:**

Pseudotime analysis was performed using the Monocle 3 package in R (version 3 0.2.0).(Trapnell et al., 2014) Counts for individual cells were preprocessed using PCA with 15 dimensions following log-normalization. Dimensional reduction was performed using a UMAP reduction with min_dist = 0.5, n_neighbors = 30, and repulsion.strength = 2.0. Cells were then clustered using Monocle 3's Louvain implementation with a resolution of 1e-5. A principal graph was then learned from the reduced dimension space using reversed graph embedding with default parameters, and cell order selection was made from the two elements at either end of the trajectory. Pseudotime trajectory heatmaps were created using the Monocle 2 package in R.

**CytoTRACE analysis:**

We utilized the recently developed bioinformatics tool CytoTRACE to compare differentiation states among cells in our dataset (https://cytotrace.stanford.edu/).(Gulati et al., 2020) This tool analyzes the number of uniquely expressed genes per cell, as well as other factors like distribution of mRNA content and number of RNA copies per gene, to calculate a score assessing the differentiation and developmental potential of each cell (lowest differentiation and highest developmental potential at 1; highest differentiation and lowest developmental potential at 0). Cells are then ordered by their predicted differentiation status. CytoTRACE analysis was performed using default parameters for each fibroblast in our dataset.

**GeneTrail analysis:**

Using GeneTrail 3, an over-representation analysis (ORA) was performed for each cell using the 500 most expressed protein-coding genes on gene sets from Gene Ontology.(Gerstner et al., 2020) P values were adjusted using the Benjamini-Hochberg procedure, and gene sets were required to have between 2 and 1,000 genes. Stacked violin plots were generated using the Scanpy package.(Wolf et al., 2018)

**CellChat receptor-ligand analysis:**

To evaluate the potential for interactions between different cell types in our dataset, we applied the recently developed CellChat platform.(Jin et al., 2021) This was implemented using our scRNA-seq Seurat object in R, in conjunction with the standalone CellChat Shiny App for its Cell-Cell Communication Atlas Explorer. Cells were binned according to the SingleR-defined cell type classifications, with fibroblast cells subsetted based on their location within either the scarring or regenerative pseudotime arms. Default parameterizations were used throughout, and Secreted Signaling, ECM-Receptor, and Cell-Cell Contact relationships were considered.

**RNA velocity analysis:**

RNA velocity analysis was performed using the scVelo package.(Bergen et al., 2020) scVelo uses a likelihood-based dynamical model to solve the full transcriptional dynamics of spliced and unspliced mRNA kinetics of each gene. RNA velocity analysis allowed us to identify transient cellular states in our dataset and to predict the directional progression of transcriptomic signatures along the identified trajectories. These predictions are based on gene-specific rates of transcription, splicing, and degradation of mRNA to estimate each cell's position in their own underlying differentiation process. The RNA velocity across all genes is then projected as a stream of arrows on the UMAP embedding.

**CODEX spatial analysis:**

To spatially phenotype the mouse specimens, we used Co-Detection by Indexing (CODEX), a novel assay in which markers are labeled with oligonucleotide-conjugated antibodies and iteratively imaged between cyclic additions and washouts of dye-labeled oligonucleotides. A custom CODEX panel was designed to assess wound cells within the tissue see **Table 1**). In brief, primary antibodies were individually barcoded and validated using the commercial supplier's protocols. OCT for mouse or paraffin blocks for human xenograft (*n* = 3 per group) were sectioned at 8 µm thickness onto coverslips for CODEX antibody staining. Antigens were retrieved by standard citrate-EDTA processing prior to addition of CODEX antibodies. Using a CODEX-integrated Keyence BZ-X instrument (Akoya Biosciences) image acquisition was then performed. Using software from Akoya Biosciences the raw images were process, with cell segmentation, and rendering.

The CODEX was visualized using Akoya Biosciences Multiplex Analysis Viewer (MAV) in ImageJ. The resulting .fcs files were then concatenated in FlowJo and imported into the Monocle3 and STvEA R packages for further analysis. After debris removal, the processed UMAP manifold was analyzed through Monocle3 with a post-manifold threshold of >10,000 cells per cluster. Analysis of the protein staining patterns was then used to assign cell types. The cell interactions were then inferred using STvEA for cell types at >2.5% of total abundance at k=20 nearest neighbors to quantify cell spatial interactions, and differential interaction maps were generated using ggraph scores.

**Visium spatial transcriptomic analysis:**

Wound specimens were rapidly harvested and flash frozen in OCT. Using the Visium Tissue Optimization Slide and Reagent Kit, permeablization time was optimized at a thickness at 10um per section and 37 minutes for mouse tissue. Following cryo-sectioning at -20 degrees onto gene expression slides the expression slide and reagent kit was used to produce sequencing libraries. The libraries were then sequences using NextSeq (Illumina). Following demultiplication the raw FASTQ files and histology images were processed by sample with the Space Ranger software for genome alignment. The raw spaceranger output files for each sample was then read into a Seurat class object in R using Seurat's Load10x function. Data was normalized using the SCT transform with default parameters. To ascertain the integration of our scRNAseq and Visium spatial analysis we employed the FindTransferAnchors() function from Seurat, which allowed the alignment of data using the two datasets. This cross platform linkage is performed serially in an unconstrained and constrained fashion.

**Statistical methods:**

Statistical testing was performed in GraphPad Prism v9 unless otherwise stated. For two-group comparisons, unpaired t-tests were used. For multi-group analysis, one-way ANOVAs were used with Tukey's *post hoc* corrections to compare groups; $p<0.05$ conferred statistical significance for all tests.
